# Identification, Characterization and Synthesis of Natural Parasitic Cysteine Protease Inhibitors – More Potent Falcitidin Analogs

**DOI:** 10.1101/2021.10.30.466580

**Authors:** Stephan Brinkmann, Sandra Semmler, Christian Kersten, Maria A. Patras, Michael Kurz, Natalie Fuchs, Stefan J. Hammerschmidt, Jennifer Legac, Peter E. Hammann, Andreas Vilcinskas, Philip. J. Rosenthal, Tanja Schirmeister, Armin Bauer, Till F. Schäberle

## Abstract

Protease inhibitors represent a promising therapeutic option for the treatment of parasitic diseases such as malaria and human African trypanosomiasis. Falcitidin was the first member of a new class of inhibitors of falcipain-2, a cysteine protease of the malaria parasite *Plasmodium falciparum*. Using a metabolomics dataset of 25 *Chitinophaga* strains for molecular networking enabled identification of over 30 natural analogs of falcitidin. Based on MS/MS spectra, they vary in their amino acid chain length, sequence, acyl residue, and *C*-terminal functionalization; therefore, they were grouped into the four falcitidin peptide families A-D. The isolation, characterization and absolute structure elucidation of two falcitidin-related pentapeptide aldehyde analogs by extensive MS/MS spectrometry and NMR spectroscopy in combination with advanced Marfey’s analysis was in agreement with the *in silico* analysis of the corresponding biosynthetic gene cluster. Total synthesis of chosen pentapeptide analogs followed by *in vitro* testing against a panel of proteases revealed selective parasitic cysteine protease inhibition and additionally low-micromolar inhibition of α-chymotrypsin. The pentapeptides investigated here showed superior inhibitory activity compared to falcitidin.

## INTRODUCTION

Parasitic diseases such as malaria and sleeping sickness (human African trypanosomiasis, HAT) are poverty-associated diseases that have a massive impact on human life, especially in tropical areas.^1,2^ Protozoan parasites of the *Trypanosoma* genus, transmitted by tsetse flies, are responsible for HAT.^2^ With fewer than 3000 cases in 2015, the number of reported cases of HAT has reached a historically low level in recent years.^2^ On the contrary, 229 million malaria cases and 409,000 deaths were estimated in 2019.^3^ *Plasmodium falciparum* is the most virulent malaria parasite, causing nearly all deaths^3^ as well as most drug-resistant infections.^4,5^ Resistance to most available antimalarials, including artemisinin-based combination regimens that are the standard treatment for falciparum malaria, is of great concern. Hence, there is a great need to discover new active compounds to serve as lead structures for the development of new antimalarial drugs.^4,5^

Papain-family cysteine proteases represent interesting therapeutic targets against several infectious diseases, including malaria (falcipains) and HAT (rhodesain).^6^ Synthetic or natural product-based small molecule inhibitors were investigated and proved falcipains and rhodesain to be promising targets.^7,8^ However, developing compounds that target these cysteine proteases into drugs has proved challenging, in part due to the difficulty of achieving selectivity.^9^ The acyltetrapeptide falcitidin and its synthetic analogs were described as first members of a new class of cysteine protease inhibitors. In a falcipain-2 assay the natural compound showed an IC50 value in the low micromolar range and the optimized synthetic analogs a sub-micromolar IC50 activity against *Plasmodium falciparum* in a standard blood-cell assay. Instead of reactive groups that covalent-reversibly or irreversibly bind to active-site cysteines, these compounds share a *C*-terminal amidated proline.^10,11^

High structural diversity in combination with diverse biological activities make natural products a valuable source in the search for new lead candidates for drug development. Technological improvements in resolution and accuracy of spectrometry methods followed by complex and large dataset analysis allowed the application of metabolomics techniques based on e.g. ultra-high-performance liquid chromatography in line with high-resolution tandem mass spectrometry (UHPLC-HRMS/MS) for the discovery and characterization of these metabolites present in a given sample.^12^ Data visualization and interpretation using tandem mass spectrometry (MS/MS) networks (molecular networking) allows automatic annotation of MS/MS spectra against library compounds as well as the identification of signals of interest and their analogs.^13^

In this study, molecular networking was applied to analyze a previously generated dataset based on bacterial organic extracts originated from 25 *Chitinophaga* strains (phylum Bacteroidetes) in which falcitidin was annotated.^14^ We aimed to investigate if additional natural analogs were produced by these strains. In total, more than 30 natural analogs were discovered, biosynthesized by *C. eiseniae* DSM 22224, *C. dinghuensis* DSM 29821, and *C. varians* KCTC 52926. They were classified into four peptide families (A-D) of falcitidin-like natural products based on variations in their amino acid chain length, sequence, acyl residue, and *C*-terminal functionalization. The analysis of MS/MS spectra revealed each family to contain truncated di- or tripeptides and larger pentapeptides, including molecules with classical *C*-terminal aldehyde moieties which are supposed to covalent-reversibly react with the active-site cysteine and serine residues of proteases.^9^ A gene encoding a multimodular non-ribosomal peptide synthetase, responsible for the production of the pentapeptide aldehydes, was identified *in silico* in selected *Chitinophaga* genomes. After isolation and structure elucidation of two natural aldehyde analogs, total synthesis of all four pentapeptide aldehydes and their carboxylic acid and alcohol derivatives was successfully achieved using a solid-phase peptide synthesis (SPPS) split approach, followed by functional *C*-terminal group interconversion. This enabled activity testing against a panel of proteases: the cysteine proteases cathepsin B and L, falcipain-2 and 3, and rhodesain, as well as the serine proteases α-chymotrypsin, matriptase-2, and the transpeptidase of *Staphylococcus aureus* sortase A. Besides a low-micromolar IC_50_ activity against α-chymotrypsin, all four tested pentapeptide aldehydes had improved inhibitory activity, compared to the reference falcitidin, against falcipain-2, and three of them also showed activity against rhodesain. In summary, the pentapeptide aldehydes display specific inhibition of the parasitic cysteine proteases falcipain-2 and −3 compared to no inhibition of human cathepsin B and L.

## RESULTS AND DISCUSSION

### Discovery of falcitidin analogs

Exploration of the genomic and metabolomic potential of the Bacteroidetes genus *Chitinophaga* revealed a high potential to find chemical novelty.^14^ In the framework of a previous metabolomics’ study of *Chitinophaga*, an inhibitor of the antimalarial target falcipain-2, falcitidin (**1**),^10^ was detected.^14^ Originally wrongly described as myxobacterium-derived tetrapeptides produced by *Chitinophaga* sp. Y23, falcitidin and its synthetic analogs are first members of a new class of cysteine protease inhibitors without a reactive group that covalent-reversibly or irreversibly binds to the active site cysteine residue.^10,11^ Here, molecular networking was carried out to analyze the chemical profiles of all 25 previously cultured *Chitinophaga* strains (Figure 1a). This facilitated the discovery of more than 30 natural analogs of falcitidin (**1**, *m/z* 548.3563 [M+H]^+^, molecular formula C_27_H_46_N_7_O_5_) (Figure 1b) biosynthesized by *C. eiseniae* DSM 22224, *C. dinghuensis* DSM 29821, and *C. varians* KCTC 52926. Analysis of the corresponding MS/MS spectra led to the assignment of four peptide families that differed in the amino acid chain at position 2 and the acyl residue of the peptide. Members of families A and B shared a fragment ion of *m/z* 222.1237 [M+H]^+^ corresponding to the molecular formula C_11_H_16_N_3_O_2_^+^. The fragment can be assigned to the described isovaleroyl (iVal) residue attached to the first amino acid histidine.^10^ In contrast, a fragment ion of *m/z* 256.1084 [M+H]^+^, corresponding to the molecular formula C_14_H_14_N_3_O_2_^+^, was detected in peptides of families C and D (Figure S1-4). A difference within the acyl moiety was assumed due to the identical number of three nitrogen atoms, also indicating a histidine at position one. Moreover, the MS/MS fragmentation pattern of family members A and B as well as C and D varied in position 2 of the amino acid chain, either carrying valine or isoleucine. Within each peptide family, members differed in chain length from two to five amino acids and their *C*-terminal functional group (Figure 1c). Interestingly, a corresponding falcitidin analog with the unusual *C*-terminal amidated (CONH_2_-group) proline, indicated by a neutral loss of 114.0794 Da, was found in each family. Falcitidin (iVal-H-I-V-P-CONH_2_) itself belongs to family B. Its analogs with the same neutral loss were assigned to the UHR-ESI-MS ion peaks at *m/z* 534.3402 [M+H]^+^ (C_26_H_44_N_7_O_5_, family A), *m/z* 568.3248 [M+H]^+^ (C_29_H_42_N_7_O_5_, family C), and *m/z* 582.3408 [M+H]^+^ (C_30_H_44_N_7_O_5_, family D). Additionally, within all families tetrapeptide-analogs carrying a *C*-terminal unmodified (COOH-group) proline and analogs with an aldehyde moiety (CHO-group) indicated by a neutral loss of 99.0686 Da (C5H9NO) were found. Besides these tetrapeptide analogs of falcitidin, truncated di- or tripeptides with *C*-terminal COOH-groups and larger pentapeptide analogs were predicted based on MS/MS spectra. The spectra of the pentapeptides, which were found in all families, revealed either phenylalanine or leucine/isoleucine in the *C*-terminal position. Besides an unmodified *C*-terminal phenylalanine (COOH-group), additional analogs carrying a *C*-terminal phenylalaninal (CHO-group) and phenylalaninol (CH_2_OH-group) were predicted. The last two analogs were proposed based on the neutral losses of 149.0841 Da (C_9_H_11_NO) and 151.1006 Da (C_9_H_13_NO), respectively. For analogs with a leucine/isoleucine at position five only analogs with a CHO- and CH2OH-group were detected, with neutral losses of 115.1031 Da (C_6_H_13_NO) and 117.1164 Da (C_6_H_15_NO), respectively (Figure 1c, Figure S1-4).

**Figure 1.**
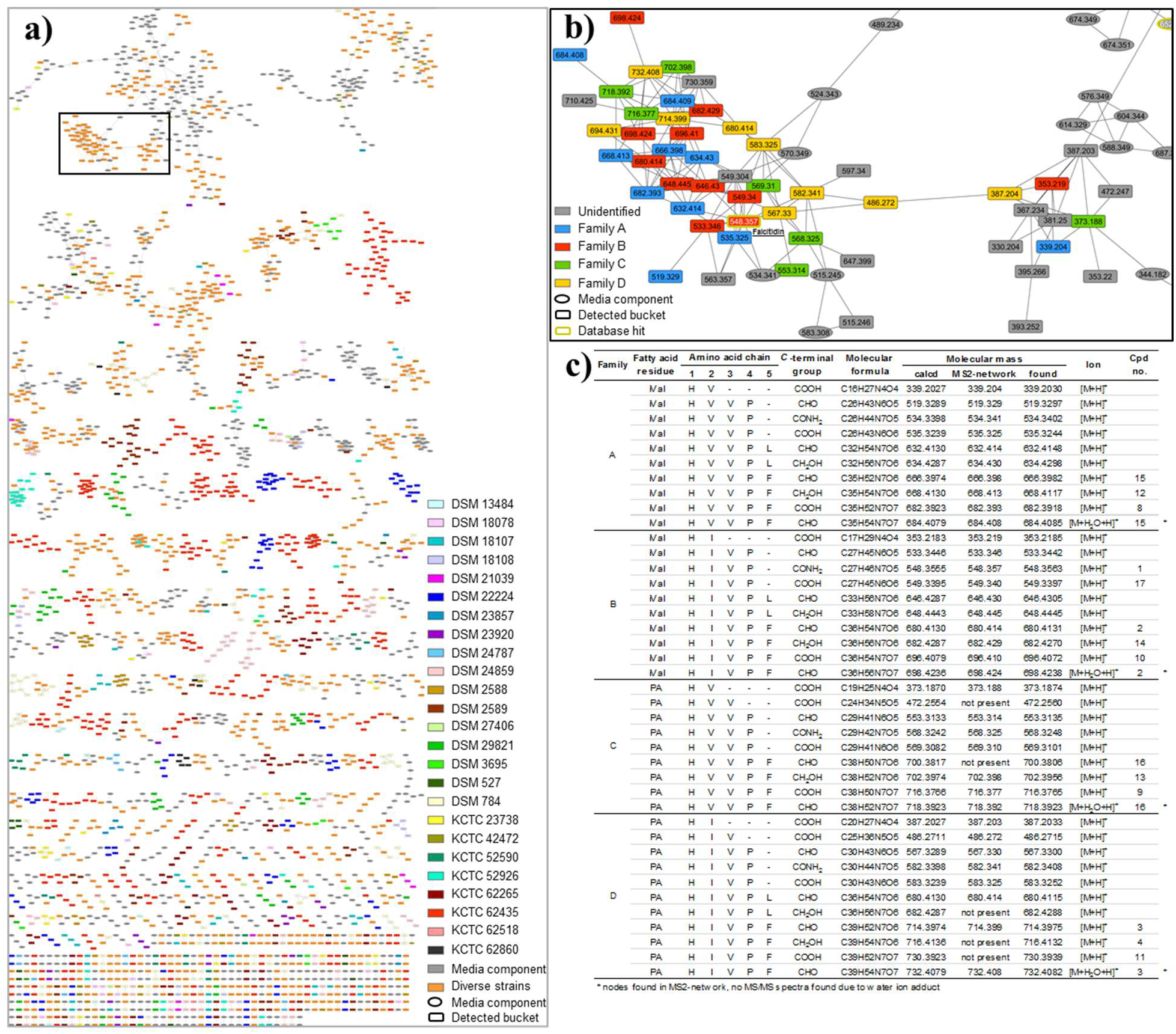
MS^2^-networking analysis revealed more than 30 natural falcitidin analogs. **A**: Complete MS^2^-network based on extracts of 25 *Chitinophaga* strains. **B**: Falcitidin network color-coded according to the four molecule families A-D. **C**: Overview of molecular features of all identified natural falcitidin analogs.

### Natural compound isolation and spectroscopic analysis

In order to confirm the proposed structures of the pentapeptide falcitidin analogs, their functional *C*-terminal groups and the unknown acyl chain of peptide families C and D, isolation and spectroscopic analysis of natural pentapeptides with *m/z* 680.4131 [M+H]^+^ (**2**, C_36_H_54_N_7_O_6_), *m/z* 714.3975 [M+H]^+^ (**3**, C_39_H_52_N_7_O_6_), and *m/z* 716.4132 [M+H]^+^ (**4**, C_39_H_54_N_7_O_6_) was achieved. Isolation was performed by adsorption of the metabolites on XAD16N resin and sequential C18-RP-HPLC followed by C18-RP-UHPLC fractionations starting from a 20 L fermentation of *C. eiseniae* DSM 22224. Based on the analysis of several two-dimensional NMR spectra, including DQF-COSY, TOCSY, ROESY, multiplicity edited-HSQC, and HMBC experiments, the structure of the isolated compound **2** was determined to be the phenylalaninal extended version of falcitidin (iVal-H-I-V-P-F-CHO), determined from the additional aromatic signals δ_H_ = 7.27 and 7.23 ppm, as well as the aldehyde protons at δ_H_ = 9.47 and 9.41 ppm and δ_C_ = 200.3 ppm (Table 1, Figure S19). The presence of two signals for several protons and carbon atoms indicated an epimerization of the stereogenic center of the *C*-terminal phenylalaninal. The analysis of the NMR data of the isolated compound **3** revealed the same peptide sequence. However, instead of the isovaleroyl moiety, compound **3** contains a phenyl acetyl (PA) residue at the *N*-terminus, leading to the final sequence PA-H-I-V-P-F-CHO (Table 2, Figure S21). Again, the protons of the terminal aldehyde moiety gave rise to a set of two signals (e.g. δ_H_ of aldehyde proton = 9.46 and 9.41 ppm), indicating the epimerization of its stereogenic center. The NMR sample of compound **4** contained a mixture of **4** and aldehyde **3**, not allowing for a complete assignment of the NMR signals. The phenylalaninol moiety, as indicated by MS/MS results, was confirmed by the presence of a second methylene group adjacent to C_α_ (δ_H_ = 3.27 ppm, δ_13C_ = 61.8 ppm), which in the COSY spectrum is correlated with an exchangeable proton at 4.7 ppm (OH). The reduction of the aldehyde function also leads to a significant high-field shift of the H_α_ and the amide proton (Figure S15), verifying the structure of compound **4** to be PA-H-I-V-P-F-CH_2_OH (Figure 2).

**Figure 2.**
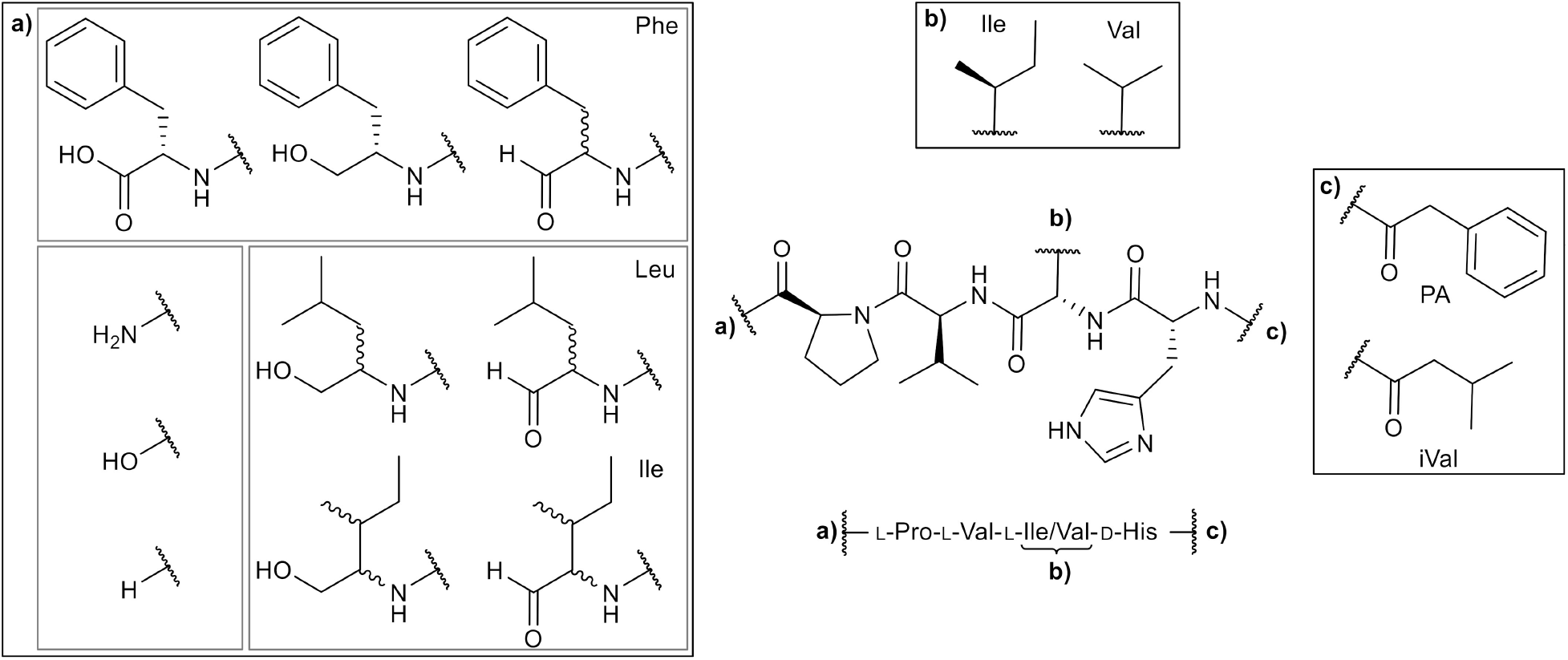
Overview of chemical structures of natural falcitidin analogs.

**Table 1.** NMR data comparison for **2** in DMSO-*d*_6_ for natural isolated one (700 MHz, 176 MHz) and synthetic one (600 MHz, 100 MHz). Due to the presence of two diastereomers (see above) two sets of signals are obtained. Therefore, for some signals two chemical shift values are given.^a^

| Position | $\delta$ $^{13}\text{C}$ [ppm] | | $\delta$ $^1\text{H}$ [ppm]; (m, $\int$, $J$) | |
| --- | --- | --- | --- | --- |
|  | natural isolated | synthetic | natural isolated | synthetic |
| Phenylalaninal |  |  |  |  |
| **CHO** | 200.3 | 200.3 | 9.47, 9.41 | 9.47 (s, 0.2H)<br>9.41 (s, 0.2H) |
| **NH** | - | - | 8.38 | 8.36 (d, 0.2H, $J = 7.7$ Hz)<br>8.34 (d, 0.2H, $J = 7.0$ Hz) |
| **α-CH** | 59.5 | 59.7 / 59.5 | 4.30 | 4.35-4.24 (m, 2H, overlay)<br>4.24-4.16 (m, 2H, overlay) |
| **β-CH<sub>2</sub>** | 33.5 | 33.6 / 33.3 | 3.13, 2.74 | 3.15-3.07 (m, 1H)<br>2.91-2.83 (m, 2H, overlay)<br>2.78-2.69 (m, 2H, overlay) |
| **γ-C<sub>quart</sub>** | 137.6 | 137.6 | - | - |
| **CH<sub>arom</sub>** | 129.3,<br>128.1,<br>126.3<br>126.2 | 129.3, 129.2,<br>129.1, 128.9,<br>128.2, 128.1,<br>126.24, 126.19 | 7.23, 7.27, n.a. | 7.29-7.12 (m, 5H) |
| Proline |  |  |  |  |
| **CO** | ~172.0 | 172.1 / 172.0 | - | - |
| **α-CH** | ~59.1 | 59.1 / 59.0 | 4.29 | 4.35-4.24 (m, 2H, overlay) |
| **β-CH<sub>2</sub>** | 29.3 | 29.31 / 29.28 | 1.92, 1.54 | 2.02-1.83 (m, 5H, overlay)<br>1.83-1.70 (m, 2H, overlay) |
| γ-CH<sub>2</sub> | ~24.4 | 24.44 / 24.38 | 1.79, 1.74 | 1.60-1.51 (m, 1H)<br>2.02-1.83 (m, 5H, overlay)<br>1.83-1.70 (m, 2H, overlay) |
| δ-CH<sub>2</sub>N | ~47.0 | 47.00 / 46.96 | 3.75, 3.53 | 3.78-3.68 (m, 1H)<br>3.57-3.47 (m, 1H) |
| Valine |  |  |  |  |
| CO | 169.6 | 169.58 / 169.55 | - | - |
| NH | - | - | 8.05 | 8.00-7.94 (m, 2H, overlay) |
| α-CH | 55.8 | 55.7 | 4.25 | 4.35-4.24 (m, 2H, overlay) |
| β-CH | 29.7 | 29.7 | 1.98 | 2.02-1.83 (m, 5H, overlay) |
| γ-CH<sub>3</sub> | 19.0 | 19.03 / 19.00 | 0.88 | 0.90-0.84 (m, 6H, overlay) |
|  | 18.5 | 18.45 / 18.41 | 0.86 |  |
| Isoleucine |  |  |  |  |
| CO | 170.7 | 170.7 | - | - |
| NH | - | - | 7.82 | 7.69-7.63 (m, 1H) |
| α-CH | ~56.3 | 56.32 / 56.29 | 4.25 | 4.24-4.16 (m, 2H) |
| β-CH | ~36.9 | 36.9 / 36.8 | 1.65 | 1.70-1.60 (m, 2H) |
| γ-CH<sub>2</sub> | 24.0 | 24.0 | 1.25 | 1.30-1.22 (m, 1H) |
|  |  |  | 0.93 | 0.98-0.90 (m, 1H) |
| γ-CH<sub>3</sub> | 15.2 | 15.2 | 0.71 | 0.76-0.67 (m, 6H, overlay) |
| δ-CH<sub>3</sub> | 11.0 | 11.0 | 0.74 | 0.76-0.67 (m, 6H, overlay) |
| Histidine |  |  |  |  |
| CO | 170.0 | 170.9 | - | - |
| NH | - | - | 8.16 | 8.00-7.94 (m, 2H, overlay) |
| α-CH | ~51.7 | 52.7 | 4.71 | 4.61-4.55 (m, 1H) |
| β-CH<sub>2</sub> | ~27.9 | 29.9* | 3.03 | 2.91-2.83 (m, 2H, overlay) |
|  |  |  | 2.85 | 2.78-2.69 (m, 2H, overlay) |
| γ-C<sub>quart</sub> | n.a. | 134.2 | - | - |
| δ-CH<sub>arom</sub> | ~116.7 | n.o. | ~7.2 | 6.76 (s, 1H) |
| ε-CH<sub>arom</sub> | 133.8 | 134.5 | ~8.7 | 7.50 (s, 1H) |
| ε-NH | - | - | - | n.o. |
| Isovaleroyl |  |  |  |  |
| CO | 171.6 | 171.4 | - | - |
| CH<sub>2</sub> | 44.3 | 44.5 | 1.96 | 2.02-1.83 (m, 5H, overlay) |
| CH | 25.4 | 25.5 | 1.89 | 2.02-1.83 (m, 5H, overlay) |
| CH<sub>3</sub> | 22.2 | 22.2 | 0.80 | 0.81 (d, 3H, $J$ = 6.4 Hz) |
| CH<sub>3</sub> | 22.1 | 22.1 | 0.77 | 0.78 (d, 3H, $J$ = 6.4 Hz) |

<sup>a</sup> n.o. - not observed; n.a. - not assigned due to line broadening; \* - data obtained from HMBC or HSQC spectrum

**Table 2.**
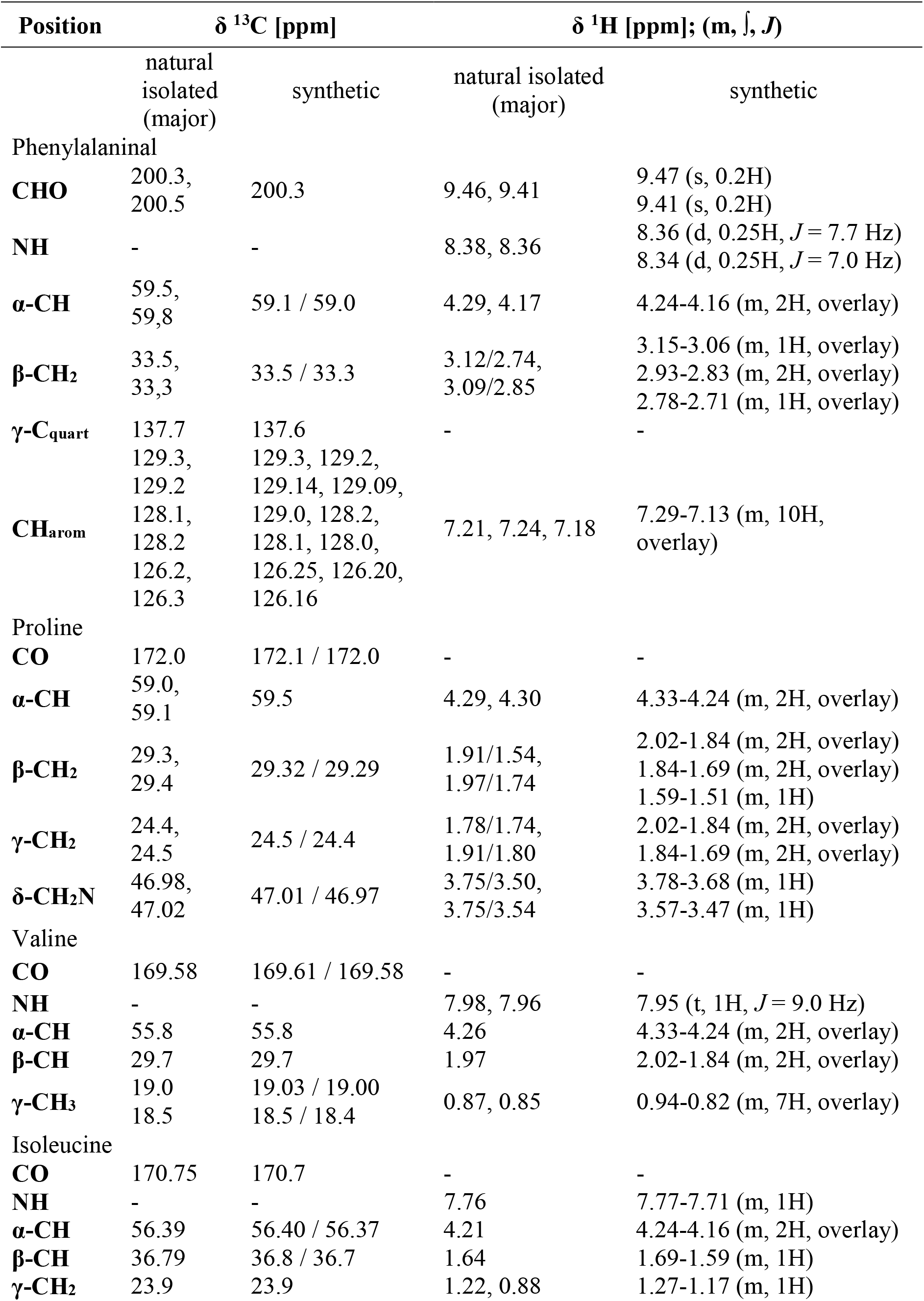
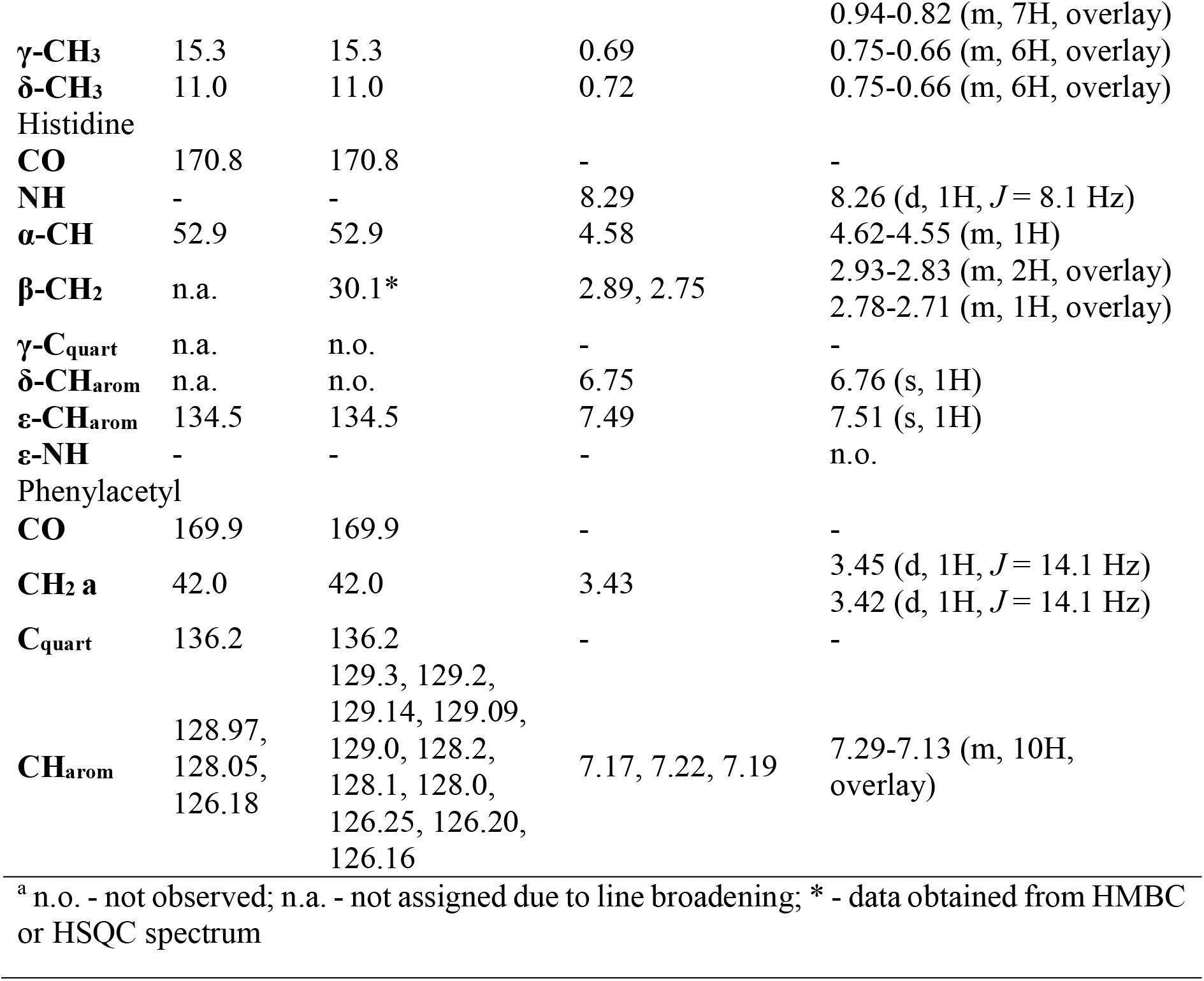
NMR data comparison for **3** in DMSO-*d*_6_ for natural isolated one (500 MHz, 126 MHz) and synthetic one (600 MHz, 100 MHz).^a^

Interestingly, UHPLC control measurements of the pure pentapeptide samples after NMR spectroscopy revealed the presence of mixtures of the pentapeptide and its corresponding tetrapeptide analog with *C*-terminal amidated proline. The NMR samples were dissolved in DMSO, afterwards dried *in vacuo*, resolved in MeOH and measured. For example within the control measurement of compound **3**, the falcitidin analog of family D *(m/z* 582.3408 [M+H]^+^, C_30_H_44_N_7_O_5_) was detected (Figure S5). Chemically, the decomposition of the pentapeptide to the unexpected amide version of the proline could be based on two hypothetical mechanisms: i) base-catalysis by the imidazole moiety of the histidine^15^ or ii) an oxidative decomposition *via* an *N*,*O*-acetal-like intermediate.^16^

Finally, advanced Marfey’s analysis was conducted to determine the absolute configuration of the amino acids.^17^ Total hydrolysis of a sample containing **3** and **4**, followed by chemical derivatization with *N_α_*-(2,4-dinitro-5-fluorophenyl)-L-valinamide (Marfey’s reagent) and UHPLC-MS comparison to reference substrates confirmed the literature known D-His-L-Val/Ile-L-Val-L-Pro configuration^10^ and the incorporation of L-phenylalanine at position five by verifying the presence of L-phenylalaninol (Figure S5). All stereogenic centers were further confirmed by total synthesis and comparison of NMR data to natural isolated compounds (Table 1 and 2).

### Identification of the biosynthetic gene cluster

The publicly available genomes of all three producers, *C. eiseniae* DSM 22224 (FUWZ01000000), *C. dinghuensis* DSM 29821 (QLMA01000000), and *C. varians* KCTC 52926 (JACVFB010000000), were scanned with antiSMASH^18^ for NRPS-type biosynthetic gene clusters (BGCs) matching the structural features of the molecules. The number and predicted substrate specificity of the A-domains, as well as precursor supply and post-assembly modifications were taken into account. BGCs congruent to the pentapeptide structure were identified in each case. Furthermore, an epimerization domain to catalyze the conversion of L- to D-amino acids is positioned in agreement with the determined stereochemistry of the molecules. Interestingly, manual search of other publicly available *Chitinophaga* genomes revealed the presence of similar BGCs in genomes of *C. varians* Ae27 (JABAIA010000000) and *C. niastensis* DSM 24859 (PYAW00000000), the latter also being included in the molecular networking analysis. However, no production was observed for *C. niastensis*. MAFFT alignment^19^ of all five BGCs allowed clear cluster boarder prediction and revealed variations in the upstream region. In conclusion, falcitidin analog biosynthesis is encoded by a single NRPS core gene 18.5 kbp in length. No additional genes encoding proteins involved in further modifications or transportation are conserved between all five BGCs (Figure 3). The NRPS starts with a starter condensation (C-starter) domain responsible for the addition of iVal or PA to the peptide core. A terminal reductase domain (TD) at the *C*-terminal end in four BGCs should be responsible for the reductive release process.^20^ The presence of alcohol and aldehyde pentapeptides suggests a similar reduction as e.g. shown in the biosynthesis of the siderophore myxochelin.^21–23^ The (peptidyl)acyl thioester attached to the carrier protein could be reduced first to an aldehyde and then to an alcohol via a four-electron reduction during the product release.^24,25^ The BGC in the genome of *C. varians* Ae27 carries a NAD_binding_4 domain instead of the TD. This domain family is sequence-related to the *C*-terminal region of the male sterility protein in arabidopsis species^26^ and a jojoba acyl-CoA reductase.^27^ The latter is known to catalyze a similar reduction reaction, with the formation of a fatty alcohol from a fatty acyl substrate.^27^

**Figure 3.**
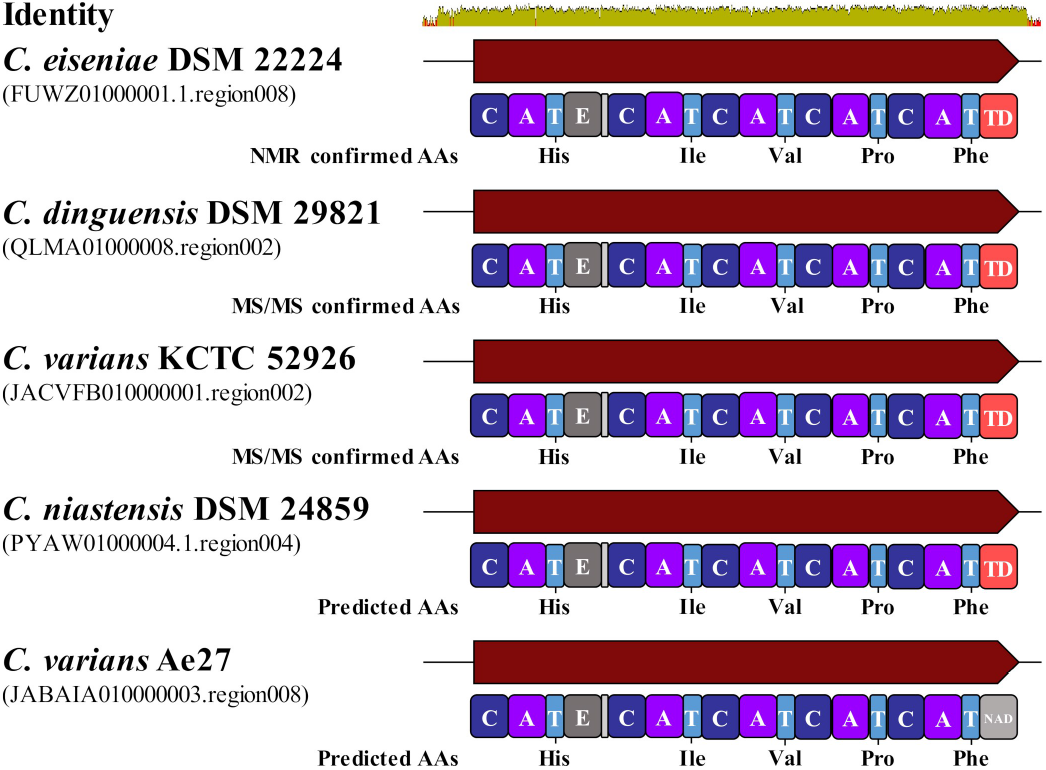
Biosynthetic gene clusters for falcitidin-like pentapeptides found in various *Chitinophaga* strains. Nucleotide alignment of all five BGCs using MAFFT alignment^19^ allowed clear cluster boarder prediction and revealed variations in the upstream region. AA, amino acids; C, Condensation domain; A, Adenylation domain; T, Peptidyl-carrier protein domain; E, Epimerization domain; TD, Terminal reductase domain; NAD, NAD_binding_4 domain.

A BGC congruent to the pentapeptide structure and the presence of the unusual tetrapeptide analog after NMR study of its corresponding pentapeptide aldehyde points toward falcitidin and its identified analogs from families A, C and D with their unusual *C*-terminal amidated proline to be degradations products. Biochemically a cleavage by a carboxypeptidase in the presence of ammonia could result in the *C*-terminal amide group of the proline, which can explain the tetrapeptides in the extract. Chemically, the unusual *C*-terminal amide group instead of the expected carboxylic acid could be based either on a base-catalyzed^16^ or an oxidative decomposition.^15^ These hypotheses and their mechanisms will need to be evaluated in further studies.

### Total synthesis of pentapeptide falcitidin analogs

The scarcity of natural material and difficulties in isolation due to co-elution of aldehydes and alcohol analogs made it necessary to synthesize the most promising pentapeptide phenylalanine-aldehydes for biological testing. The application of a split approach solid-phase peptide synthesis (SPPS)^28^ followed by functional group interconversion gave access to a variety of falcitidin pentapeptide analogs of family A, B, C and D (Scheme 1).

**Scheme 1.**
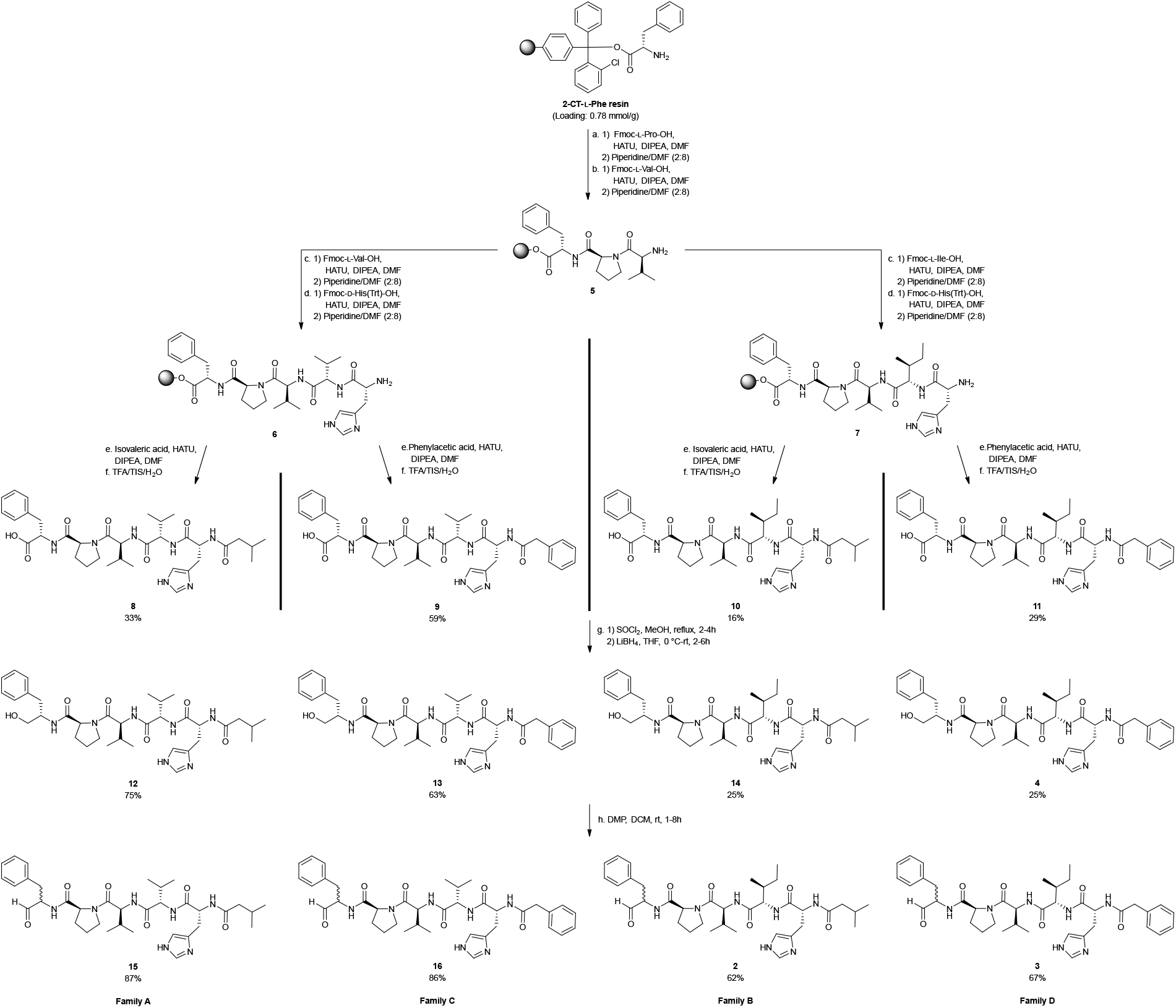
SPPS split approach (a-f) and functional group interconversions (g-h) to chosen falcitidin pentapeptide analogs of family A, B, C and D. Standard SPPS conditions apply and reactions were carried out at room temperature if not noted otherwise. Reduction of the acid to the alcohol was achieved *via* the corresponding methyl ester. The aldehyde was obtained by Dess-Martin oxidation; the stereogenic center of phenylalanine could not be retained. Detailed conditions can be found in the Supporting Information.

Splitting the synthesis after the third amino acid and for the attachment of the two different fatty acids allows the synthesis of all four chosen acid analogs (**2**, **3**, **15**, and **16**) based on 2-chlorotrytil-L-Phe resin (2-CT-L-Phe). The functionalization of the peptide acid yields the corresponding methyl ester, the alcohol and the aldehyde. Reduction to the alcohol was done *via* the methyl ester, after direct reduction of the acid could not be achieved. The methyl ester was gained by conversion with thionyl chloride in methanol, followed by the reduction with LiBH4 in THF. For the oxidation to the aldehyde Dess-Martin periodinane (DMP) was chosen as a very mild reagent. We found, that small traces of methanol as a stabilizer in DCM as the solvent made the reaction take longer, up to eight hours instead of one, and was responsible for incomplete conversion. Epimerization of the stereogenic center of the aldehydes could not be avoided, nor was it necessary to be avoided based on the data for the natural isolated compounds. They were obtained as diasteromers for the phenylalanine moiety of roughly equal proportions based on NMR data (Figure S17, S18, S20, S22). Overall yields of the SPPS for the four different chains varied between 16-59%. The reduction yielded the corresponding alcohols over two steps with 25-75% yield and oxidation to the aldehydes achieved 62-87% yield (Scheme 1).

For reference purposes, falcitidin (**1**) was synthesized using the same SPPS approach instead of the literature known liquid phase method.^11^ This afforded acid analog **17** with a yield of 78% and, after amidation, falcitidin (**1**) with a 41% over all yield (Scheme 2). Additionally, the acid analog **17** and falcitidin (**1**) themselves present great functional groups for further derivatization and the introduction of different war heads, like a nitrile or azide group, to increase the potency as a potential inhibitor.^29,30^

**Scheme 2.**
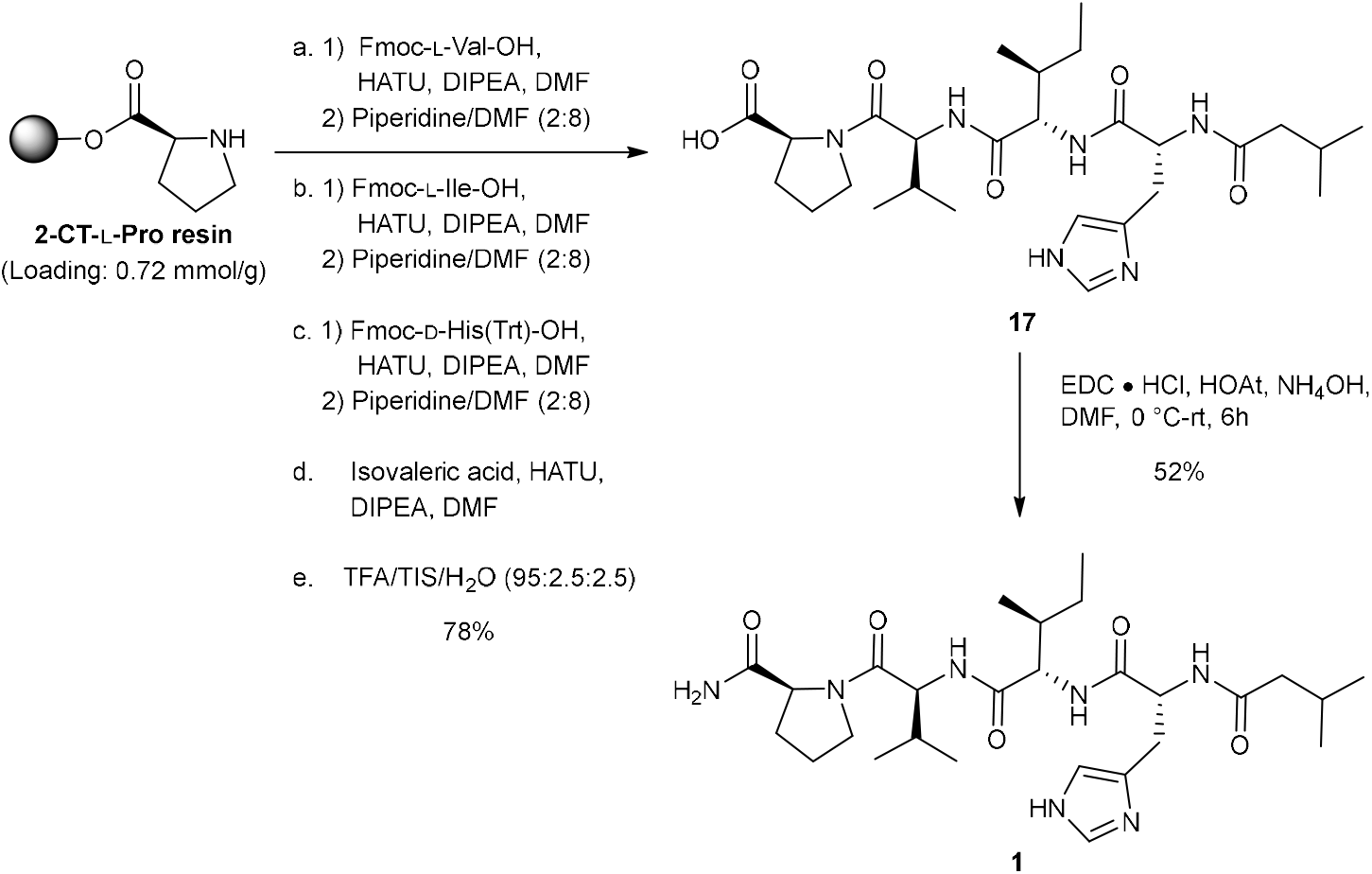
Synthesis of falcitidin (**1**) *via* acid analog **17**.

### Bioactivity

Falcitidin (**1**) was previously reported to display an IC_50_ of 6 μM against falcipain-2.^10^ Therefore, the falcitidin pentapeptide aldehyde analogs **2**, **3**, **15** and **16** were tested in a similar *in vitro* assay against falcipain-2 together with **1** as control. All four aldehydes were active with an IC_50_ of 41.5 μM (**15**) to 23.7 μM (**2**). However, the originally described activity of **1** could not be reproduced, with an IC_50_ >50 μM. This indicated differences in assay sensitivity, which made it difficult to put the aldehyde activities in line with the literature. Aldehydes **3** and **15** displayed activities also against the related *P. falciparum* cysteine protease falcipain-3 (66% sequence identity with falcipain-2), with an IC_50_ of 45.4 μM and 42.5 μM, respectively. Inhibition of both falcipains is of importance, since falcipain-3 is able to compensate for knockout of falcipain-2.^31^ A counter screen against human cysteine proteases cathepsin B and L and sortase A of *Staphylococcus aureus* as a surrogate for unrelated cysteine proteases revealed no activity for pentapeptide aldehydes **2**, **3**, **15** and **16** as well as the *C*-terminal acid (**11**) and alcohol (**4**) of **3**, thus demonstrating selectivity over these off-targets. Activities with an IC_50_ of 57.7 μM (**2**) to 17.1 μM (**15**) were also observed for the falcipain-homologue cysteine protease of *Tryoanosoma brucei rhodesiense* rhodesain. Most interestingly, aldehydes **3**, **15** and **16** displayed higher activities, with IC_50_s of 3.7 μM (**15**) and 1.5 μM (**16**), against α-chymotrypsin, which was tested as a prototype of human serine proteases, while the serine membrane protease matriptase-2 (TMPRSS6) was not inhibited (Table 3). To further elucidate these observations, molecular docking studies were performed.

**Table 3.**
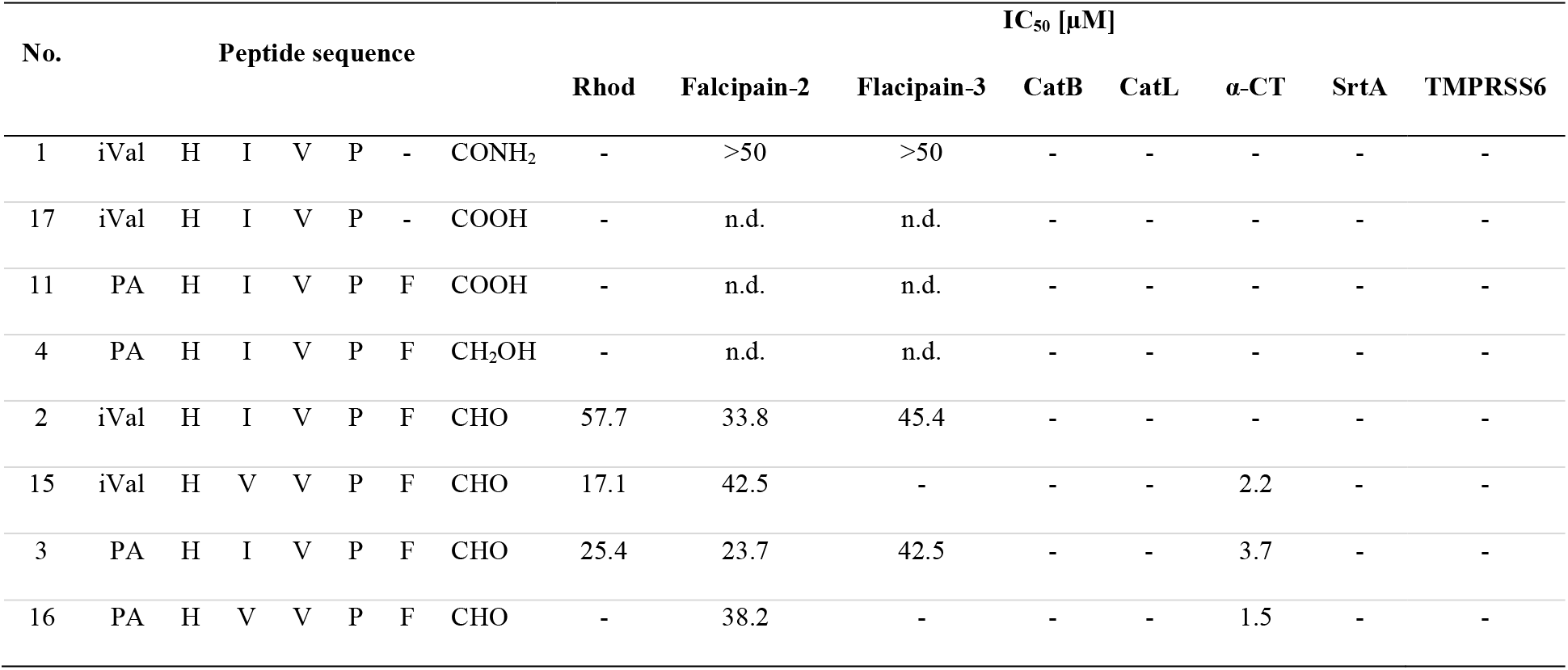
Overview of protease activity data. n.d. = not determined; − = not active

### Docking studies

Due to the high number of rotatable bonds and the associated degrees of conformational freedom in the molecules under elucidation, docking of the full-length peptides is challenging.^32–34^ Hence, truncated tripeptides with an *N*-terminal acetyl (ace)-cap to avoid a nonpresent charge of full-length inhibitors (**2**, **3**, **4**, **11**, **15** and **16**) were used for docking studies. First, a conventional non-covalent docking was performed. However, aldehydes are known electrophilic warheads and able to form covalent-reversible hemithioacetal adducts with catalytic cysteine residues.^35^ Therefore, the predicted poses with special attention to the distance between the nucleophilic sulfur of the catalytic cysteine residue and the electrophilic carbon of the aldehyde moiety were elucidated, and an additional covalent docking was performed. The tripeptides mimic the orientation of the protease substrates by addressing S3-S1 (Figure 4a and b). Based on these results (Table S15), the low inhibitory potency of the *C*-terminal alcohol (**4**) and acid (**11**) moieties as well as that for falcitidin cannot simply be explained by non-covalent interactions, as the docking scores are within a similar range or even higher when compared to the aldehydes. However, based on predicted binding poses the covalent reaction between aldehydes and catalytic cysteine residues, which might contribute to higher affinity, seems likely for molecules with proteinogenic L-Phe as P1 residue, with a distance of 3.0 Å between the electrophilic carbon atom and the nucleophilic sulfur of Cys for both falcipain-2 and falcipain-3, and the aldehyde oxygen coordinated by hydrogen bonds in the oxyanion hole, but rather unlikely for D-Phe (distance of 7.3 and 5.0 Å for falcipain-2 and falcipain-3, respectively). Additionally, only small differences between covalent and non-covalent binding modes were observed, indicating that no larger conformational changes need to take place during or after reaction. While testing against α-chymotrypsin was conducted to demonstrate selectivity over human off-target serine proteases, the high affinity is reasonable, as inhibitors with aromatic moieties deeply buried in the S1 pocket of the protease were reported previously, as well as cleavage preference after Phe.^36–38^ Aldehydes are able to form covalent reversible hemiacetal adducts with serine^39^; moreover, an increased potency is indicated by proximity of the aldehyde to the catalytic Ser-195 residue in the predicted binding modes for both L- and D-Phe at P1 (Figure 4c, Table S15).

**Figure 4.**
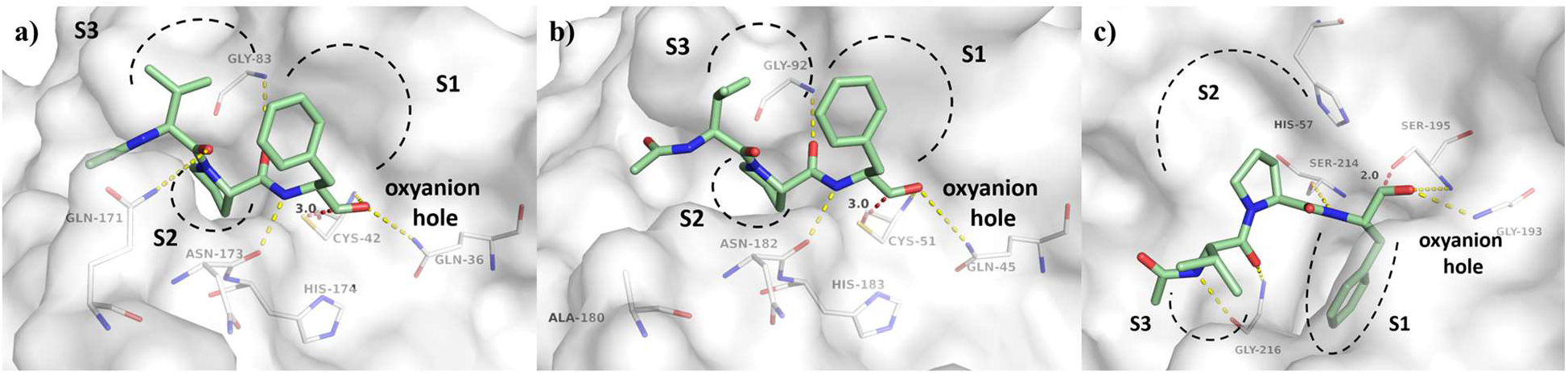
Non-covalent docking predicted binding modes of truncated peptidic protease inhibitor ace-Val-Pro-Phe-aldehyde (green carbon atoms) in complex with falcipain-2 (**A**, PDB-ID 3BPF), falcipain-3 (**B**, PDB-ID 3BPM) and chymotrypsin (**C**, PDB-ID 1AFQ). Proteases are depicted as white transparent surface, for clear view only residues forming polar interactions (yellow dashed lines) and catalytic Cys/Ser and His residues are labeled and depicted as lines. Substrate binding sites S1-S3 are schematically indicated. The distance between nucleophilic sulfur (Cys-42 in falcipain-2 and Cys-51 in falcipain-3) or oxygen (Ser-195 in chymotrypsin) of the catalytic center to the electrophilic carbon atom of the aldehyde is depicted as a dashed red line and labeled with its distance in Å.

Falcitidin (**1**) was found to display poor whole-cell activity against chloroquine-sensitive *P. falciparum* strain 3D7 (IC_50_ >10 μM), while a structure-activity-relationship (SAR) study identified a synthetic trifluoromethyl analog displaying sub-micromolar IC_50_ activity.^11^ Together with our results, this allows further development of the SAR, with the aim to increase both potency and selectivity with an improved peptidic recognition sequence. However, no general selectivity-issue against serine proteases can be expected, as matriptase-2 was not inhibited by the compounds. Additionally, the docking studies revealed a rather unlikely distance for covalent binding of D-Phe-analogs. With all compounds having been tested as mixtures of D- and L-Phe due to the aldehyde’s natural epimerization of its stereogenic center,^40,41^ further analogs with e.g. other warheads preventing epimerization such as a nitrile group^29,30^ should ideally be synthesized and tested in L-configurations only.

## CONCLUSION

In this study, metabolic networking analysis of *Chitinophaga* strains led to the discovery of over 30 *N*-acyl oligopeptides, structurally related to falcitidin, an inhibitor of the antimalarial cysteine protease falcipain-2.^10^ Isolation and structure elucidation of two novel natural pentapeptide aldehydes validated the MS/MS fragmentation pattern analysis. A BGC congruent to the pentapeptide structure was identified and indicated falcitidin and its tetrapeptide analogs carrying the *C*-terminally amidated proline to be degradation products from the described pentapeptide aldehydes. Total synthesis gave access to the most promising aldehyde analogs and allowed their biological profiling. A selective *in vitro* activity against parasitic cysteine proteases rhodesain, falcipain-2 and falcipain-3, together with a low-micromolar IC_50_ inhibition of the serine protease α-chymotrypsin was observed. This forms the basis for future studies to develop optimized derivatives with increased potency and selectivity against targeted proteases.

## EXPERIMENTAL SECTION

### General Experimental Procedures

For all UHPLC-QTOF-UHR-MS and MS/MS measurements a quadrupole time-of-flight spectrometer (LC-QTOF maXis II, Bruker Daltonics, Bremen, Germany) equipped with an electrospray ionization source in line with an Agilent 1290 infinity II LC system (Agilent Technologies, CA, Unites States) was used. C18 RP-UHPLC [ACQUITY UPLC BEH C18 column (130 Å, 1.7 μm, 2.1 x 100 mm)] was performed at 45 °C with the following linear gradient (A: H2O, 0.1% HCOOH; B: CH3CN, 0.1% HCOOH; flow rate: 0.6 mL/min): 0 min: 95% A; 0.30 min: 95% A; 18.00 min: 4.75% A; 18.10 min: 0% A; 22.50 min: 0% A; 22.60 min: 95% A; 25.00 min: 95% A. A 50 to 2000 *m/z* scan range at 1 Hz scan rate was used to acquire mass spectral data. The injection volume was set to 5 μL. MS/MS experiments were performed at 6 Hz and the top five most intense ions in each full MS spectrum were targeted for fragmentation by higher-energy collisional dissociation at 25 eV using N_2_ at 10^-2^ mbar. Precursors were excluded after 2 spectra, released after 0.5 min and reconsidered if the intensity of an excluded precursor increased by factor 1.5 or more. Data were analyzed using the Bruker DataAnalysis 4.0 software package. Specific rotation was determined on a digital polarimeter (P3000, A. Krüss Optronic GmbH, Germany). Standard wavelength was the sodium D-line with 589 nm. Temperature, concentration (g/100 mL) and solvent are reported with the determined value.

### NMR Spectroscopy

NMR spectra of natural isolated falcitidin analogs were acquired on a Bruker AVANCE 700 spectrometer (700 MHz for ^1^H, 176 MHz for ^13^C) and a Bruker AVANCE 500 spectrometer (500 MHz for ^1^H, 126 MHz for ^13^C). Both instruments were equipped with a 5 mm TCI cryo probe. For structure elucidation and assignment of proton and carbon resonances 1D-^1^H, 1D-^13^C, DQF-COSY, TOCSY (mixing time 80 ms), ROESY (mixing time 150 ms), multiplicity edited-HSQC, and HMBC spectra were acquired. NMR spectra of synthesized molecules were recorded on an AVANCE III HD 600 spectrometer (600 MHz for ^1^H, 151 MHz for ^13^C) from Bruker Biospin (Bruker Biospin GmbH, Rheinstetten, Germany). ^1^H- and ^13^C-chemical shifts are reported in ppm and were referenced to the corresponding residual solvent signal (DMSO-*d*_6_: δ_C_ = 39.52 ppm, δ_H_ = 2.50 ppm). δ_C_ shifts marked with an * (asterisk) were not observed in the ^13^C NMR spectrum, but were obtained either from HMBC or HSCQ data.

### MS/MS Networking

Molecular networking was performed following established protocols.^13,42^ In brief, parent ions are represented by a list of fragment mass/intensity value pairs within the raw data (*.d files) converted with MSConvert (ProteoWizard package32) into plain text files (*.mgf). These ions are included in the final network once they share at least six fragments (tolerance Δppm 0.05) with at least one partner ion.^43^ Deposited compounds from an *in silico* fragmented^44^ commercial database (Antibase 2017^45^) as well as our *in-house* reference compound MS/MS database were included in the final network to highlight known NPs. A visualization of the network was constructed in Cytoscape v3.6.0.^46^ Edges were drawn between scan nodes with a cosine similarity >0.7^47^.

### Strain Fermentation and Purification of Falcitidin Analogs

A pre-culture (R2A, 100 mL in 300 mL Erlenmeyer flask) of *C. eiseniae* DSM 22224 was inoculated from plate (R2A) and incubated at 28 °C with agitation at 180 rpm for 3 days. A 20 L fermentation in medium 3018 (1 g/L yeast extract, 5 g/L casitone, pH 7.0) inoculated with 2% (v/v) pre-culture was carried out separate 2 L flasks filled with 500 mL culture volume at 28 °C with agitation at 180 rpm for 4 days. The culture broth was subsequently freeze-dried using a delta 2-24 LSCplus (Martin Christ Gefriertrockungsanlagen GmbH, Osterode am Harz, Germany). The sample was extracted with one-time culture volume CH3OH, evaporated to dryness using rotary evaporation under reduced pressure, and resuspened in 3 L of 10% CH3OH/H2O. The extract was loaded onto a XAD16N column (1 L bed volume) and eluted step-wise with 10%, 40%, 60%, 80%, and 100% CH3OH (two-times bed volume each). The 80 and 100% fractions containing falcitidin analogs were pooled and the sample was adjusted to 200 mg/mL in methanol to achieve further separation using preparative C18-RP-HPLC (Synergi 4 μm Fusion-RP 80 Å (250 x 21.2 mm)) by eluting in a linear gradient increasing from 25% to 75% CH3CN (+0.1% HCOOH) in 22 min. Fractions of interest were concentrated to 100 mg/mL for semi-preparative C18-RP-HPLC (Synergi 4 μm Fusion-RP 80 Å (250 x 10 mm)) using a linear gradient from 15% to 50% CH3CN (+0.1% HCOOH) in 29 min. Final purification of samples of interest (100 mg/mL) were achieved using UHPLC on a ACQUITY UPLC BEH C18 column (130 Å, 1.7 μm, 100 x 2.1 mm) eluting in a isocratic gradient of 27.50% CH_3_CN (+0.1% HCOOH) in 18 min. In total, isolation yielded 1.5 mg of **2**, 1 mg of **3**, and 2 mg of a mixture of **3** and **4**.

(2*R*/*S*)-1-((3-methylbutanoyl)-D-histidyl-L-isoleucyl-L-valyl)-*N*-((*S*)-1-oxo-3-phenylpropan-2-yl)pyrrolidine-2-carboxamide (**2**). Amorphous, white powder; see Table 1 for ^1^H and ^13^C NMR data; positive HR-ESIMS *m/z* 680.4131 [M+H]^+^, calculated mass for C_36_H_54_N_7_O_6_^+^ 680.4130; Δ= 0.15 ppm.

(2*R*/*S*)-*N*-((*S*)-1-oxo-3-phenylpropan-2-yl)-1-((2-phenylacetyl)-D-histidyl-L-isoleucyl-L-valyl)pyrrolidine-2-carboxamide (**3**). Amorphous, white powder; see Table 2 for ^1^H and ^13^C NMR data; positive HR-ESIMS *m/z* 714.3975 [M+H]^+^, calculated mass for C_39_H_52_N_7_O_6_^+^ 714.3974; Δ= 0.14 ppm.

### Advanced Marfey’s Analysis

The absolute configuration of all amino acids was determined by derivatization using Marfey’s reagent.^17^ Stock solutions of amino acid standards (50 mM in H_2_O), NaHCO_3_ (1 M in H_2_O), and *N_α_*-(2,4-dinitro-5-fluorophenyl)-L-valinamide (L-FDVA, 70 mM in acetone; Sigma Aldrich, St. Louis, MO, Unites States) were prepared. Commercially available and synthesized standards were derivatized using molar ratios of amino acid to L-FDVA and NaHCO3 (1/1.4/8). After stirring at 40 °C for 3 h, 1 M HCl was added to obtain concentration of 170 mM to end the reaction. Samples were subsequently evaporated to dryness and dissolved in DMSO (final concentration 50 mM). L- and D-amino acids were analyzed separately using C18 RP-UHPLC-MS (A: H_2_O, 0.1% HCOOH; B: CH_3_CN, 0.1% HCOOH; flow rate: 0.6 mL/min). A linear gradient of 15-75% B in 35 min was applied to separate all amino acid standards. Total hydrolysis of the peptide sample containing **3** and **4** was carried out by dissolving 250 μg in 6 M DCl in D_2_O and stirring for 7 h at 160 °C. The sample was subsequently evaporated to dryness. Samples were dissolved in 100 μL H_2_O, derivatized with L-FDVA and analyzed using the same parameters as described before.

### Fluorometric Assays

Rhodesain (Rhod),^48–50^ *Staphylococcus aureus* sortase A (SrtA)^51^ and human matriptase-2 (TMPRSS6)^52^ were expressed and purified as published previously, cathepsins B and L (CatB, CatL; human liver, Calbiochem) and α-chymotrypsin (α-CT; Sigma Aldrich, St. Louis, MO, Unites States) were purchased. For these proteases except SrtA, fluorescence increase upon cleavage of the fluorogenic substrates was monitored without incubation with a TECAN Infinite F200 Pro fluorimeter (excitation λ = 365 nm; emission λ = 460 nm) in white, flat-bottom 96-well microtiter plates (Greiner bio-one, Kremsmünster, Austria) with a total volume of 200 μL. Inhibitors and substrates were prepared as stock solutions in DMSO to a final DMSO-content of 0.5%. Inhibitors were screened at final concentrations of 20 μM and eventually at 1 μM. IC_50_ values were determined for compounds for which an inhibition of >50% at a concentration of 20 μM was observed. All assays were performed in technical triplicates and normalized to the activity of DMSO instead of the inhibitors by measuring the increase of fluorescence signal over 10 min. The data were analyzed using GraFit V 5.0.13^53^ (Erithracus Software, Horley, UK, http://www.erithacus.com/grafit/).

Cbz-Phe-Arg-AMC (Bachem, Bubendorf BL, Switzerland) was used as a substrate for Rhod, CatB and CatL. The enzymes were incubated at room temperature in enzyme incubation buffer (Rhod: 50 mM sodium acetate pH 5.5, 5 mM EDTA, 200 mM NaCl and 2 mM DTT; CatB/L: 50 mM TRIS-HCl pH 6.5, 5 mM EDTA, 200 mM NaCl, 2 mM DTT) for 30 min. 180 μL assay buffer (Rhod: 50 mM sodium acetate pH 5.5, 5 mM EDTA, 200 mM NaCl and 0.005% Brij35; CatB/L: 50 mM TRIS-HCl pH 6.5, 5 mM EDTA, 200 mM NaCl and 0.005% Brij35) were added to the 96-well plates, afterwards the respective enzyme in enzyme incubation buffer (5 μL; to yield final concentrations for Rhod 0.01 μM, CatB 0.1 μM, CatL 0.2 μM) followed by 10 μL DMSO (control) or inhibitor solution in DMSO and finally substrate (5 μL; final concentrations for Rhod 10 μM, CatB 100 μM and CatL 6.5 μM) were added.^54^

Transpeptidation efficacy of SrtA was performed *in vitro* as described previously.^51^ Briefly, SrtA was diluted in assay buffer (50 mM TRIS-HCl pH 7.50, 150 mM NaCl) to a final concentration of 1 μM. The FRET-pair substrate Abz-LPETG-Dap(Dnp)-OH (Genscript, Piscataway, NJ, Unites States) and the tetraglycine (Sigma Aldrich, St. Louis, MO, Unites States) were added at 25 μM and 0.5 mM, respectively.

α-Chymotrypsin (final concentration 0.4 μM) was dissolved in assay buffer containing 50 mM TRIS-HCl pH 8.0, 100 mM NaCl, 5 mM EDTA. Suc-Leu-Leu-Val-Tyr-AMC (Bachem, Bubendorf BL, Switzerland) was used as a substrate at a final concentration of 52.5 μM.^55^ Proteolytic activity of matriptase-2 was measured in a final concentration of 2.5 nM Enzyme in 180 μL reaction buffer (50 mM TRIS-HCl (pH 8.0), 150 mM NaCl, 5 mM CaCl_2_, 0.01% (v/v) T_x-100_). After addition of the inhibitors, reaction was started without further incubation by adding the substrate Boc-Leu-Arg-Arg-AMC (Bachem. Bubendorf BL, Switzerland, K_M_ = 36.1 ± 5.8 μM) to a final concentration of 100 μM.

The recombinant enzymes falcipain-2 and falcipain-3 were expressed and purified as previously described.^56^ Stock solutions of the compounds, substrate and the positive control E-64 (Sigma Aldrich, St. Louis, MO, Unites States) were prepared at 10 mM in DMSO. The compounds were incubated in 96-well white flat bottom plates with 30 nM of recombinant falcipain-2 or 3 at room temperature in assay buffer (100 mM sodium acetate, pH 5.5 with 5 mM DTT) for 10 min. After incubation, the fluorogenic substrate Z-Leu-Arg-AMC (R&D Systems, Minneapolis, MN, Unites States) was added at a concentration of 25 μM in a final assay volume of 200 μL. Fluorescence was monitored with a Varioskan Flash (Thermo Fisher Scientific Inc., Waltham, MA, Unites States) with excitation 355 nm and emission 460 nm. The IC_50_ was calculated using GraphPad Prism (GraphPad Software, San Diego, CA, Unites States) based on a sigmoidal dose response curve.

### Molecular docking

Molecular docking was performed against falcipain-2 (complex with E-64, PDB-ID 3BPF),^57^ falcipain-3 (complex with Leupeptin, PDB-ID 3BPM)^57^ and chymotrypsin (complex with D-Leucyl-L-phenylalanyl-p-fluorobenzylamide PDB-ID 1AFQ).^36^ Conventional non-covalent template-based docking was performed with HYBRID v3.3.0.3 (OpenEye Scientific Software, Santa Fe, NM, US, http://www.eyesopen.com).^58,59^ The receptor was prepared using the make_receptor tool version 3.3.0.3 under default settings for potential field generation around the reference ligand for 3BPM and 1AFQ. As the complexed ligand E-64 of 3BPF does not reach toward S1 of falcipain-2, the potential field was generated using leupeptin from the aligned structure of 3BPM (falcipain-3) for which a similar binding behaviour for falcipain-2 and falcipain-3 can be expected.^60^ 800 ligand conformers per molecule for docking were generated using omega pose (OMEGA v3.1.0.3, OpenEye Scientific Software, Santa Fe, NM, US, http://www.eyesopen.com).^61^ Covalent docking was performed using MOE 2020.09.^62^ Using the covalent reaction of acetalization from aldehyde and Cys/Ser (for E-64 re-docking from epoxide to beta-hydroxythioether) a rigid docking with GB/VI scoring was applied for initial placement of 50 poses from which the ten best scoring ones were refined using the ASE scoring function. Docking setups were validated by re-docking of crystallographic reference ligands by pose-inspection and RMSD calculation (Table S15).

## ASSOCIATED CONTENT

### Supporting Information

The Supporting Information is available free of charge at LINK.

MS/MS-based assignment of peptide sequences, Marfey’s analysis, detailed description of all syntheses, ^1^H and ^13^C spectra and data of all synthesized compounds and the natural isolated ones, docking results. (PDF)

## AUTHOR INFORMATION

### Authors

**Stephan Brinkmann** – *Fraunhofer Institute for Molecular Biology and Applied Ecology (IME)*, *Branch for Bioresources*, *35392 Giessen*, *Germany*;

**Sandra Semmler** – *Fraunhofer Institute for Molecular Biology and Applied Ecology (IME)*, *Branch for Bioresources*, *35392 Giessen*, *Germany*

**Christian Kersten** – *Institute of Pharmaceutical and Biomedical Sciences*, *Johannes Gutenberg University Mainz*, *55128 Mainz*, *Germany*;

**Maria A. Patras** – *Fraunhofer Institute for Molecular Biology and Applied Ecology (IME)*, *Branch for Bioresources*, *35392 Giessen*, *Germany*;

**Michael Kurz** – *Sanofi-Aventis Deutschland GmbH*, *R&D*, *65926 Frankfurt am Main*, *Germany*

**Natalie Fuchs** – *Institute of Pharmaceutical and Biomedical Sciences*, *Johannes Gutenberg University Mainz*, *55128 Mainz*, *Germany*

**Stefan J. Hammerschmidt** – *Institute of Pharmaceutical and Biomedical Sciences*, *Johannes Gutenberg University Mainz*, *55128 Mainz*, *Germany*;

**Jennifer Legac** – *Department of Medicine*, *University of California*, *San Francisco*, *94143 California*, *United States*

**Peter E. Hammann** – *Evotec International GmbH*, *37079 Göttingen*, *Germany*

**Andreas Vilcinskas** – *Fraunhofer Institute for Molecular Biology and Applied Ecology (IME)*, *Branch for Bioresources*, *35392 Giessen*, *Germany*; *Institute for Insect Biotechnology*, *Justus-Liebig-University of Giessen*, *35392 Giessen*, *Germany*;

**Philip. J. Rosenthal** – *Department of Medicine*, *University of California*, *San Francisco*, *94143 California*, *United States*;

**Tanja Schirmeister** – *Institute of Pharmaceutical and Biomedical Sciences*, *Johannes Gutenberg University Mainz*, *55128 Mainz*, *Germany*;

### Author Contributions

^λ^ S. Brinkmann and S. Semmler contributed equally.

### Notes

The authors declare no competing financial interest.

## ACKNOWLEDGMENT

The authors would like to thank Jens Glaeser, Sören M. M. Schuler, Yolanda Kleiner and Christoph Pöverlein for valuable discussions. We thank Christoph Hartwig, Victoria Öhler and the NMR department of the Justus-Liebig-University Giessen for technical assistance. We further thank OpenEye Scientific for free academic software licenses and Torsten Steinmetzer and coworkers from the Philipps University of Marburg for the kind gift of matriptase-2 coding plasmid for recombinant protein expression.

