## Supporting Information for "Identification, Characterization and Synthesis of Natural Parasitic Cysteine Protease Inhibitors – More Potent Falcitidin Analogs"

**Figure S6.** Comparison of the Marfey derivatization products of a mixed sample containing the DCl hydrolysates of **3** and **4** and commercially available amino acid standards (histidine, isoleucine, valine, proline, phenylalaninol) derivatized with L-FDVA. **A**: Commercially available L-amino acid standards. **B**: Commercially available D-amino acid standards. **C**: hydrolysate of a mixture of **3** and **4**. ........................................................................................................ 10

n.o. - not observed * - data obtained from HMBC or HSQC spectrum

If not noted otherwise, NMR data and spectra are of synthetic molecules. NMR spectra and data of natural isolated compounds are declared as such.

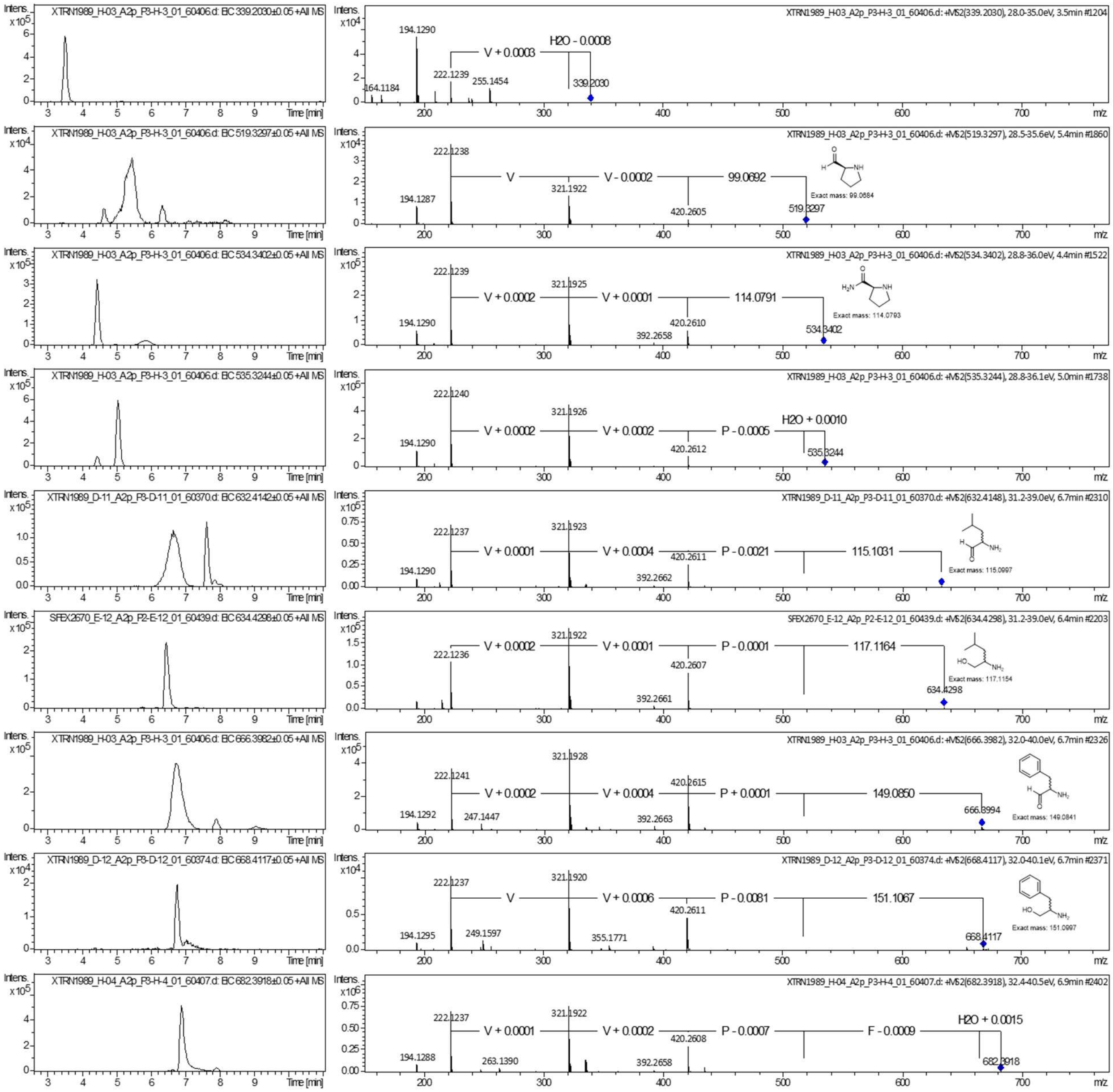

**Figure S1.** $MS^2$-spectra of falcitidin peptide family A.

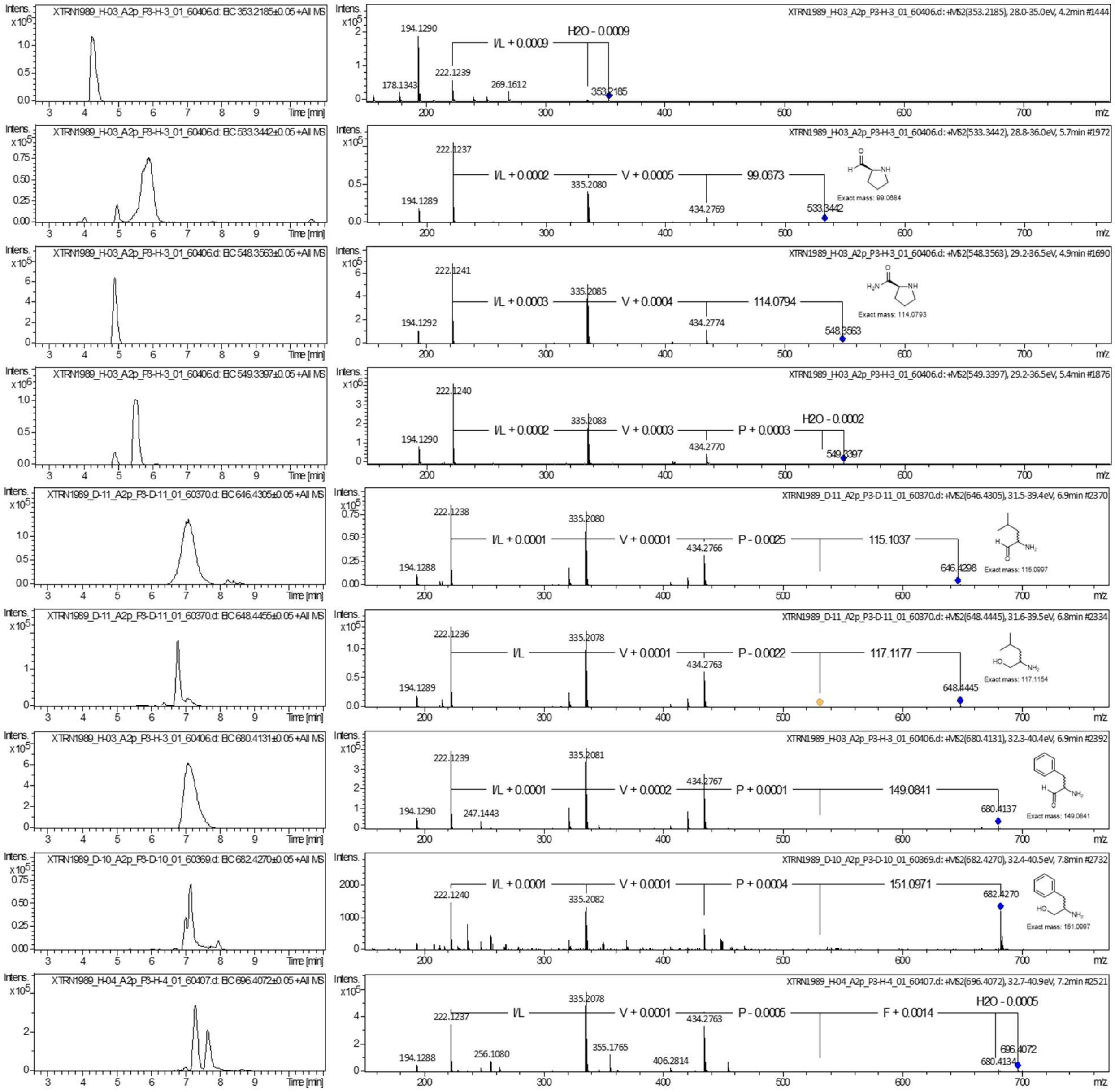

**Figure S2.** MS$^2$-spectra of falcitidin peptide family B.

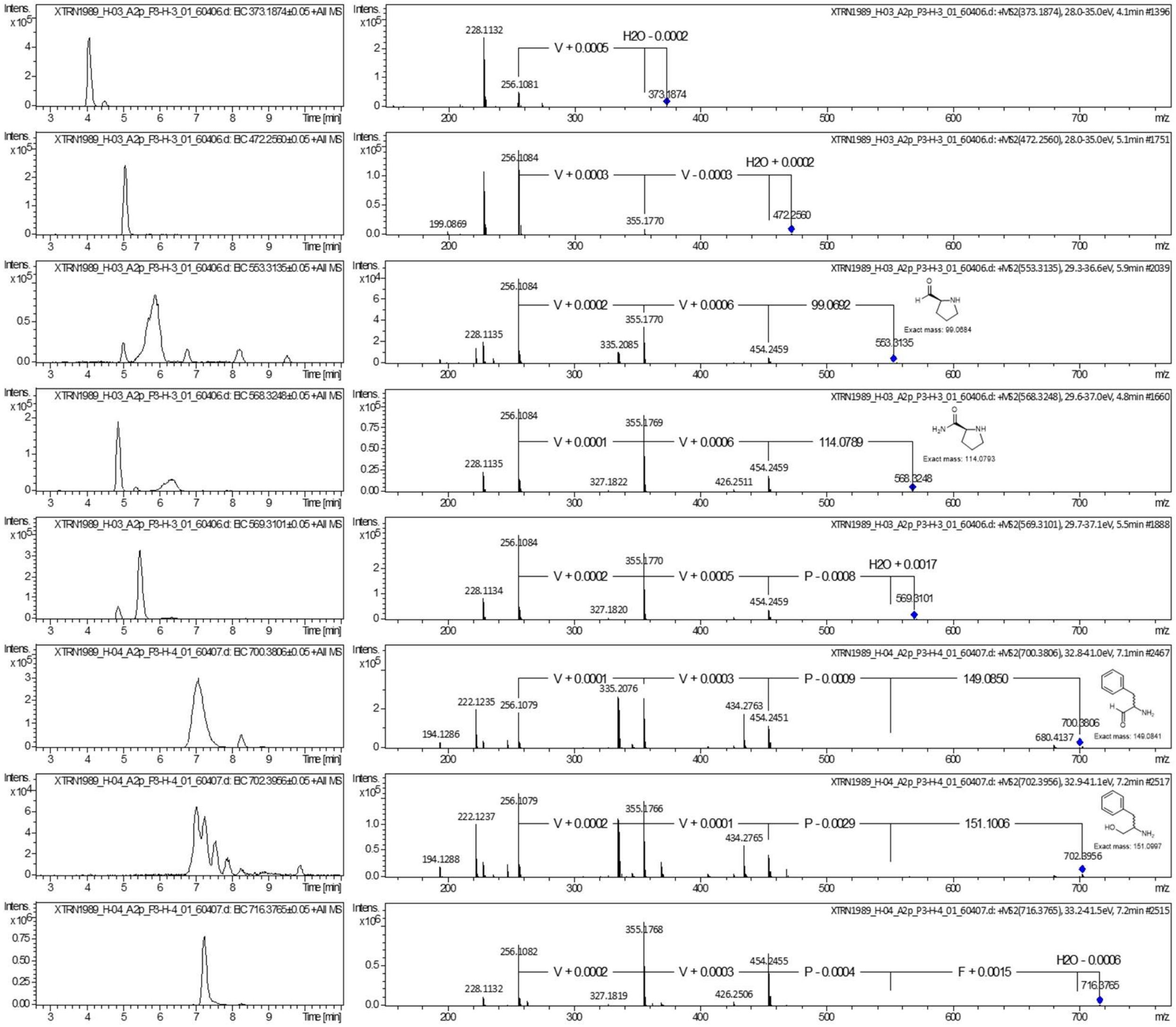

**Figure S3.** MS$^2$-spectra of falcitidin peptide family C.

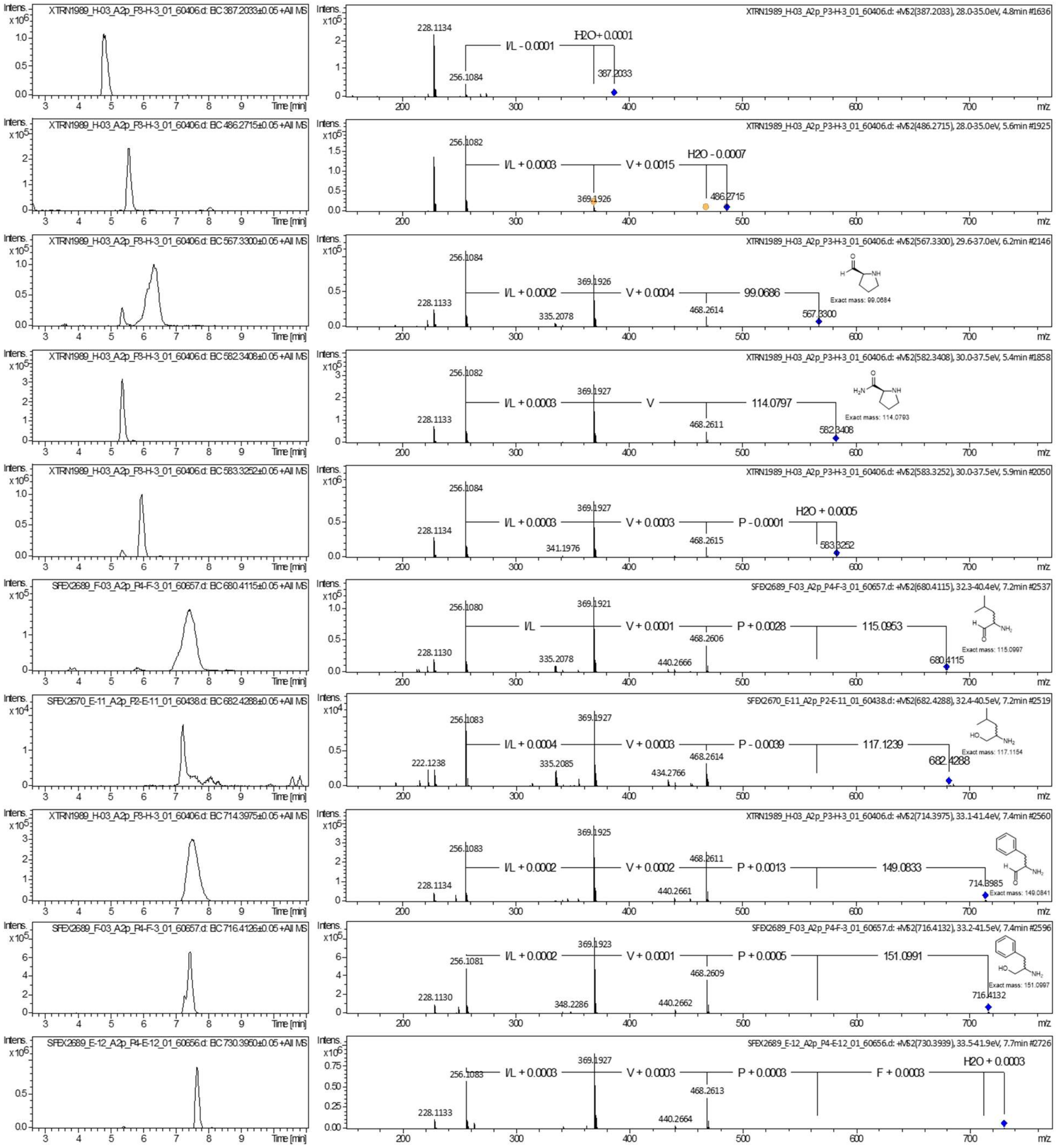

**Figure S4.** MS$^2$-spectra of falcitidin peptide family D.

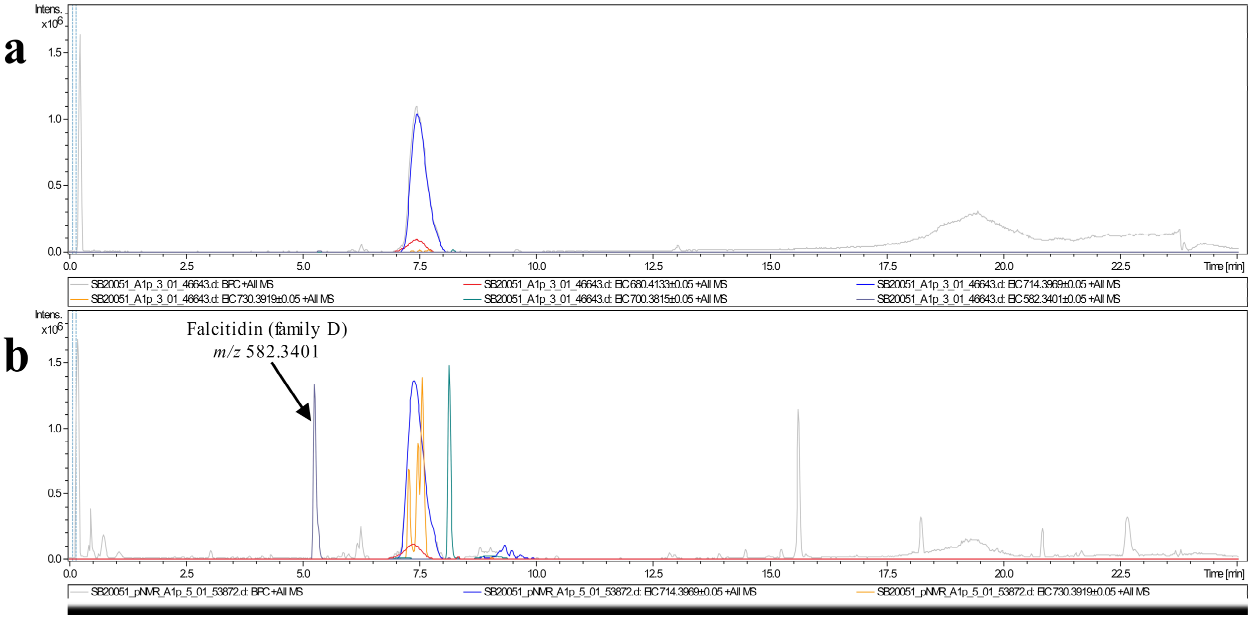

**Figure S5.** LC-MS chromatogram of the NMR sample of compound **3** before (**A**) and after NMR study (**B**).

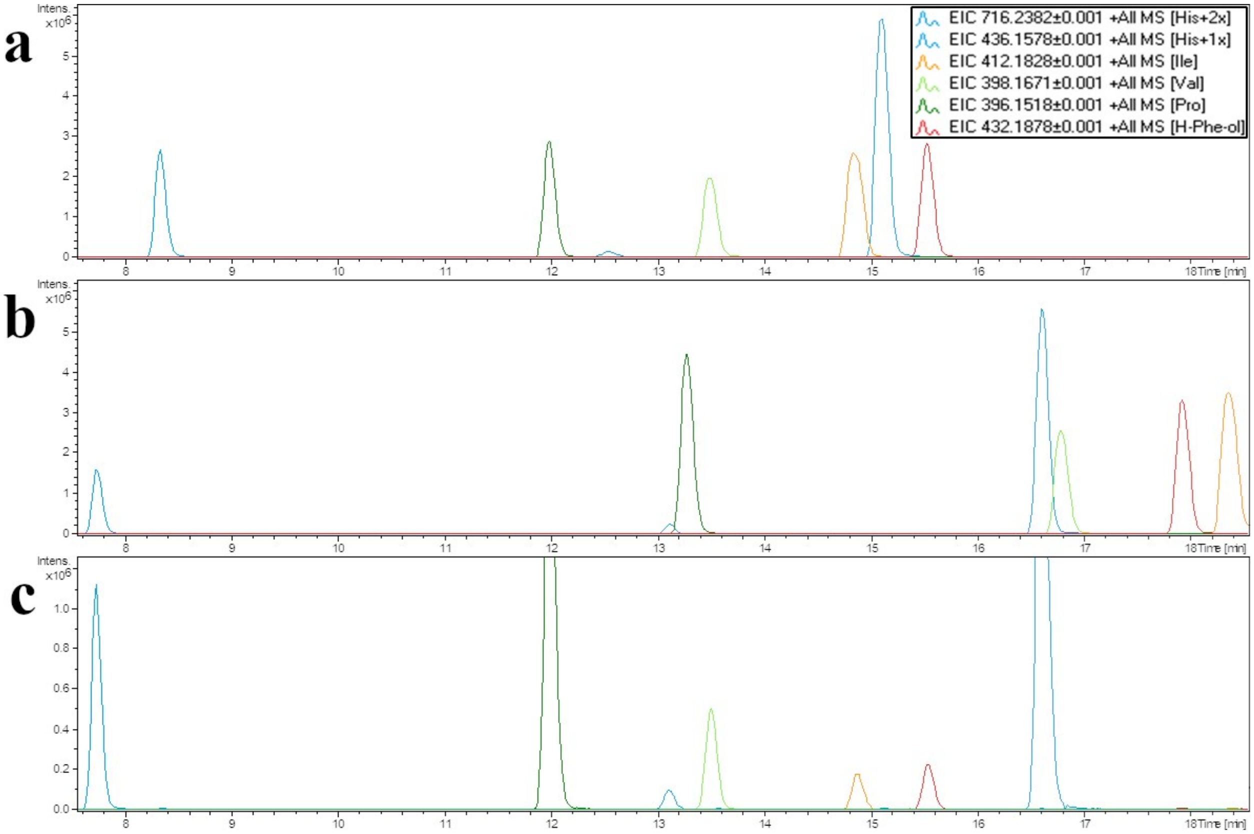

**Figure S6.** Comparison of the Marfey derivatization products of a mixed sample containing the DCl hydrolysates of **3** and **4** and commercially available amino acid standards (histidine, isoleucine, valine, proline, phenylalaninol) derivatized with L-FDVA. **A**: Commercially available L-amino acid standards. **B**: Commercially available D-amino acid standards. **C**: hydrolysate of a mixture of **3** and **4**.

### Synthesis

#### General procedures

##### Coupling of the amino acids and fatty acid[1]

If not noted otherwise, all reactions were carried out in a custom-built solid-phase peptide synthesis vessel with a G2 filter and a diameter of 3, 4 or 5 cm at room temperature. Argon was used for agitation of the resin. 20% Piperidine in DMF and the cleavage cocktail were freshly prepared on the day of use.

The Fmoc-protected amino acid or fatty acid (3 equiv) and HATU (2.9 equiv) dissolved in DMF (5-15 mL) were added to the swelled resin (1 equiv), followed by DIPEA (6 equiv) and the resin was agitated for 1-3 hours.

Each coupling step was monitored as described in the general method part for the LC-MS sample preparation. Fmoc-deprotection was carried out after each coupling step was completed, as indicated by LC-MS result.

##### LC-MS and UHR-MS sample preparation[1]

The reaction progress of each coupling of the Fmoc-protected amino acids was monitored using LC-MS. For that a few beads of the resin were sampled in a 2 mL SPPS syringe, washed once with DMF and then 2-3 times with DCM. The vessel was closed and 20% HFIP in DCM was added (1-1.5 mL), which changed the color of the beads from yellow-orange to a dark red which faded over time. The mixture was shaken for 15-30 min and the filtrate was directly used for LC-MS measurement. For UHR-MS measurement the filtrate was concentrated *in vacuo* and redissolved in AcCN or DMSO.

##### Fmoc-deprotection[1]

The mixture was filtered and the remaining resin was washed 5 times with DMF. 20% Piperidine in DMF (20-30 mL) was added. After agitation for 2-5 min it was filtrated, rinsed with DMF and the process was repeated four more times. Then the resin was washed with DMF three times.

##### Cleavage from the resin[1]

To the washed resin, the cleavage cocktail consisting of TFA:TIS:$H_2O$ (95:2.5:2.5) (40 mL) was added, coloring the mixture a dark red. The resin was agitated for 25-30 min after which the supernatant was drained and the process was repeated once. The combined filtrates were reduced under pressure and then dried further using lyophilization.

##### Synthesis of the methyl ester from the corresponding acid

For the esterification of the peptide, the acid (1 equiv) was dissolved in anhydrous methanol and cooled to 0 °C. . Thionyl chlorid (1.5 equiv) was added dropwise and the mixture was stirred under

---

[1] W. Chan, P. White, Fmoc Solid Phase Peptide Synthesis Practical Approach, Oxford University Press, Oxford, **1999**.

a gentle reflux for 2-4 h. The reaction progress was monitored using LC-MS or UHR-MS. The solvent was evaporated *in vacuo* and the residue was dried further under high vacuum. The crude product was directly used for the synthesis of the alcohol, without further purification.

**Reduction of the methyl ester to the alcohol**

The methyl ester of the peptide (1 equiv) was suspended in anhydrous THF and cooled to 0 °C. $LiBH_4$ (4 equiv) was added and the reaction was stirred at room temperature for 4-8 h. The reaction progress was monitored periodically using LC-MS and if needed more $LiBH_4$ (2 equiv) was added at 0 °C. The reaction was quenched using sat. $NH_4Cl$ solution. Water was added until all formed salts were dissolved. The layers were separated and the aqueous layer was extracted with ethyl acetate three times. The combined organic phase was washed once with brine, dried over $MgSO_4$, filtered and the solvent was evaporated under reduced pressure. Purification by HPLC yielded the product as a white powder or a colorless solid.

#### Dess-Martin oxidation of the alcohol to the aldehyde

The alcohol (1 equiv) was suspended in anhyd. DCM and DMP (15% in DCM, 3 equiv) was added. The mixture was stirred at room temperature for a total of 2 hours. The reaction progress was monitored every 25 min by UHR-MS. DMP (1 equiv, 0.5 equiv) was added as soon as no more significant conversion of the alcohol to the aldehyde could be observed. At the time a conversion of over 95% was observed, the reaction was quenched with methanol and concentrated *in vacuo*. Purification by HPLC yielded the product as a colorless solid. Note that the DCM used, must not be stabilized with methanol!. Amylene as a stabilizing reagent worked fine.

(3-methylbutanoyl)-D-histidyl-L-isoleucyl-L-valyl-L-proline (**17**)

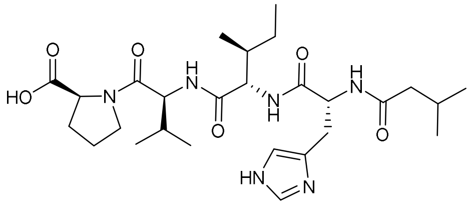

**17**

The H-L-proline-chlorotrityl resin (n = 0.72 mmol/g, 2.70 g, 1.95 mmol) was swelled in DMF for 30 min. After removal of the solvent, Fmoc-(L)-valine-OH (1.987 g, 5.846 mmol) and HATU (2.153 g, 5.662 mmol), dissolved in DMF (5 mL), were added, followed by DIPEA (1986 µL, 11.68 mmol) and more DMF (15 mL). The mixture was agitated for 1.5 h. After Fmoc-deprotection, Fmoc-(L)-isoleucine-OH (2.062 g, 5.834 mmol) and HATU (2.149 g, 5.652 mmol), dissolved in DMF (5 mL) were added, followed by DIPEA (1986 µL, 11.68 mmol) and more DMF (15 mL). The mixture was agitated for 1 h. Following Fmoc-deprotection, Fmoc-D-histidine(Trt)-OH (3.621 g, 5.843 mmol) and HATU (2.147 g, 5.647 mmol), solved in DMF (10 mL) were added, then DIPEA (1986 µL, 11.68 mmol) and more DMF (10 mL). The mixture was agitated for 1.5 h. After Fmoc-deprotection HATU (2.147 g, 5.647 mmol), solved in DMF (3 mL) was added to the resin, then isovaleric acid (642 µL, 5.844 mmol) and DIPEA (1986 µL, 11.68 mmol). More DMF (15 mL) was added and the mixture was agitated for 1.5 h. After draining the supernatant, the resin was washed with DMF (3x), DMC (2x) and MeOH (2x). The cleavage was performed as described in the general method section. The crude product was washed three times with ice cold diethyl ether and dried under reduced pressure. Purification of 40% of the crude (1.935 g) using semi preparative HPLC (5-40-95% AcCN + 0.1% FA, NUCLEODUR® C18 Gravity SB, 3 µm, 250 x 10 mm, flow rate: 2 mL/min) yielded **17** as a white wax like film (328.7 mg, 31%, 0.599 mmol, overall yield calculated to be: 78%).

**$^1$H-NMR** (DMSO-$d_6$, 600 MHz): $\delta_H$ [ppm] = 8.97 (d, 1H, *J* = 1.1 Hz, ε-C*H*$_{arom}$ His), 8.15 (d, 1H, *J* = 8.4 Hz, N*H* His), 8.05 (d, 1H, *J* = 8.3 Hz, N*H* Val), 7.84 (d, 1H, *J* = 8.8 Hz, N*H* Ile), 7.35 (s, 1H, δ-C*H*$_{arom}$ His), 4.74 (ddd, 1H, *J* = 8.2, 8.2, 6.6 Hz, α-C*H* His), 4.27 (dd, 1H, *J* = 9.0, 9.0 Hz, α-C*H* Val), 4.25 (dd, 1H, *J* = 9.0, 7.0 Hz, α-C*H* Ile), 4.17 (dd, 1H, *J* = 8.7, 5.0 Hz, α-C*H* Pro), 3.81 (ddd, 1H, *J* = 9.8, 6.4, 6.4 Hz, δ-C*H*$_2$N a Pro), 3.56 (ddd, 1H, *J* = 10.1, 6.4, 6.4 Hz, δ-C*H*$_2$N b Pro), 3.06 (dd, 1H, *J* = 15.1, 6.1 Hz, β-C*H*$_2$ a His), 2.86 (dd, 1H, *J* = 15.1, 8.7 Hz, β-C*H*$_2$ b His), 2.16-2.09 (m, 1H, β-C*H*$_2$ a Pro), 2.02-1.94 (m, 3H, C*H*$_2$ iVal, β-C*H* Val), 1.94-1.85 (m, 2H, C*H* iVal, γ-CH$_2$ a Pro), 1.85-1.79 (m, 2H, β-C*H*$_2$ b Pro, γ-C*H*$_2$ b Pro), 1.69-1.62 (m, 1H, β-C*H* Ile), 1.28-1.20 (m, 1H, γ-C*H*$_2$ a Ile), 0.97-0.92 (m, 1H, γ-C*H*$_2$ b Ile), 0.91 (d, 1H, *J* = 6.8 Hz, γ-C*H*$_3$ a Val), 0.88 (d, 1H, *J* = 6.8 Hz, γ-C*H*$_3$ b Val), 0.80 (d, 1H, *J* = 6.8 Hz, C*H*$_3$ a iVal), 0.76 (d, 1H, *J* = 6.4 Hz, C*H*$_3$ b iVal), 0.74 (t, 1H, *J* = 7.4 Hz, γ-C*H*$_3$ Ile), 0.71 (d, 1H, *J* = 6.7 Hz, δ-C*H*$_3$ Ile).

**$^{13}$C-NMR** (DMSO-$d_6$, 151 MHz): $\delta_C$ [ppm] = 173.1 (*C*OOH Pro), 171.7 (*C*O iVal), 170.7 (*C*O Ile), 169.8 (*C*O His), 169.6 (*C*O Val), 133.7* (*C*$_{arom}$ His), 129.5 (*C*$_{quart}$ His), 116.8 (*C*$_{arom}$ His), 58.5 (α-*C*H Pro), 56.2 (α-*C*H Ile), 55.7 (α-*C*H Val), 51.3 (α-*C*H His), 46.8 (δ-*C*H$_2$-N Pro), 44.3 (*C*H$_2$ iVal), 36.9 (β-*C*H Ile), 29.8 (β-*C*H Val), 28.7 (β-*C*H$_2$ Pro), 27.3 (β-*C*H$_2$ His), 25.4 (*C*H iVal), 24.6 (γ-*C*H$_2$ Pro), 24.0 (γ-*C*H$_2$ Ile), 22.14 (*C*H$_3$ iVal), 22.06 (*C*H$_3$ iVal), 18.8 (γ-*C*H$_3$ Val), 18.4 (γ-*C*H$_3$ Val), 15.2 (δ-*C*H$_3$ Ile), 10.9 (γ-*C*H$_3$ Ile).

Additional found signals: $\delta_H$ [ppm] = Broad peak between $\delta_H$ 4.08 to 3.60 ($H_2O$), 2.99, 2.54 (DMSO). ε-N*H* His and COO*H* Pro were not observed. $\delta_C$ [ppm] = 158.0, 40.4 (DMSO).

**UHR-MS (ESI-TOF)** *m/z* calcd for $C_{27}H_{45}N_6O_6$: 549.3395 $[M+H]^+$; found: 549.3401 $[M+H]^+$

**Specific rotation** $[\alpha]_D^{21}$ = -24.3° (c = 1.44, $CH_3OH$)

(*S*)-1-((3-methylbutanoyl)-D-histidyl-L-isoleucyl-L-valyl)pyrrolidine-2-carboxamide, falcitidin (**1**)

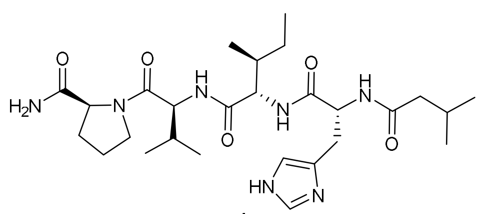

1

Peptide **17** (100.3 mg, 0.1828 mmol) was dissolved in DMF and cooled to 0 °C. HOAt (25.1 mg, 0.184 mmol) and EDC · HCl (36.9 mg, 0.192 mmol) were added and the mixture was stirred for 15 min at 0 °C. Aqueous $NH_4OH$ solution (25%, 220 µL, 1.41 mmol) was added dropwise and the yellow mixture was stirred at room temperature for 6 h. It was then concentrated *in vacuo*. Purification of the crude using semi preparative HPLC (5-35-95% AcCN + 0.1% FA, NUCLEODUR® C18 Gravity SB, 3 µm, 250 x 10 mm, flow rate: 2 mL/min) yielded **1** as a colorless syrup (51.7 mg, 0.0944 mmol, 52%).

**$^1$H-NMR** (DMSO-$d_6$, 600 MHz): $\delta_H$ [ppm] = 8.02 (d, *J* = 8.2 Hz, N*H* His), 7.99 (d, 1H, *J* = 8.3 Hz, N*H* Val), 7.86 (s, 1H, ε-C*H*$_{arom}$ His), 7.70 (d, 1H, *J* =8.8 Hz, N*H* Ile), 7.23 (s, 1H, CON*H*$_2$ Pro), 6.90 (s, 1H, δ-C*H*$_{arom}$ His), 6.81 (s, 1H, CON*H*$_2$ Pro), 4.62 (ddd, *J* =8.3, 8.3, 6.1 Hz, α-C*H* His), 4.27 (dd, 1H, *J* =8.4, 8.4 Hz, α-C*H* Val), 4.23 (dd, 1H, *J* =8.7, 6.9 Hz, α-C*H* Ile), 4.21 (dd, 1H, *J* = 8.5, 4.4 Hz, α-C*H* Pro), 3.78-3.73 (m, 1H, δ-C*H*$_2$N a Pro), 3.58-3.52 (m, 1H, δ-C*H*$_2$N b Pro), 2.92 (dd, *J* =14.9, 5.9 Hz, β-C*H*$_2$ a His), 2.76 (dd, *J* =15.1, 8.8 Hz, β-C*H*$_2$ b His), 2.03-1.97 (m, 2H, β-C*H*$_2$ a Pro, β-C*H* Val), 1.96 (d, *J* =6.7 Hz, C*H*$_2$ iVal), 1.94-1.86 (m, 2H, γ-C*H*$_2$ a Pro, C*H* iVal), 1.82-1.74 (m, 2H, γ-C$H_2$ b Pro, β-C$H_2$ b Pro), 1.69-1.61 (m, 1H, β-C*H* Ile), 1.31-1.22 (m, 1H, γ-C$H_2$ a Ile), 0.98-0.91 (m, 1H, γ-C$H_2$ b Ile), 0.90 (d, 1H, *J* =6.8 Hz, γ-C$H_3$ a Val), 0.87 (d, 1H, *J* =6.7 Hz, γ-C$H_3$ b Val), 0.81 (d, *J* =6.5 Hz, C$H_3$ a iVal), 0.78 (d, *J* =6.5 Hz, C$H_3$ b iVal), 0.74 (t, 1H, *J* =7.4 Hz, δ-C$H_3$ Ile), 0.71 (d, 1H, *J* =7.0 Hz, γ-C$H_3$ Ile).

**$^{13}$C-NMR** (DMSO-$d_6$, 151 MHz): $\delta_C$ [ppm] = 173.4 (*C*ONH$_2$ Pro), 171.5 (*C*O iVal), 170.70 (*C*O Ile), 170.67 (*C*O His), 169.5 (*C*O Val), 134.3 (weak, ε-*C*H$_{arom}$ His), 132.3 (weak, γ-$C_{quart}$ His), 117.0 (weak, δ-*C*H$_{arom}$ His), 59.2 (α-*C*H Pro), 56.3 (α-*C*H Ile), 55.8 (α-*C*H Val), 52.3 (α-*C*H His), 47.0 (δ-*C*H$_2$N Pro), 44.4 (*C*H$_2$ iVal), 36.9 (β-*C*H Ile), 29.7 (β-*C*H Val), 29.3 (β-*C*H$_2$ Pro, β-*C*H$_2$ His), 25.5 (*C*H iVal), 24.5 (γ-*C*H$_2$ Pro), 24.0 (γ-*C*H$_2$ Ile), 22.2 (*C*H$_3$ iVal), 22.1 (*C*H$_3$ iVal), 19.1 (γ-*C*H$_3$ Val), 18.4 (γ-*C*H$_3$ Val), 15.2 (δ-*C*H$_3$ Ile), 11.0 (γ-*C*H$_3$ Ile).

Additional found signals: $\delta_H$ [ppm] = 8.14 (FA), 3.17 (MeOH), 2.54 (DMSO). ε-N*H* His was not observed. $\delta_C$ [ppm] = 163.0 (FA), 48.6 (MeOH), 40.4 (DMSO).

**UHR-MS (ESI-TOF)** *m/z* calcd for $C_{27}H_{46}N_7O_5$: 548.3555 $[M+H]^+$; found: 548.3556 $[M+H]^+$

**Specific rotation** $[\alpha]_D^{21}$ = -70.0° (c = 1.00, $CH_3OH$), the determined specific rotation corresponds to literature.[10]

2-Chlorotrityl-L-Phe-L-Pro-L-NH$_2$ (**5**)

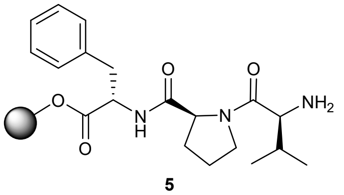

**5**

H-L-phenylalanine-chlorotrityl resin (n = 0.78 mmol/g, 15.01 g, 11.71 mmol) was swelled in DMF for 30 min. After removal of the solvent, Fmoc-(L)-proline-OH (11.84 g, 35.09 mmol) and HATU (12.89 g, 33.90 mmol) were added, dissolved in a small volume of DMF, followed by DIPEA (12.0 mL, 70.6 mmol) and more DMF. The mixture was agitated for 2 h. After Fmoc-deprotection, Fmoc-(L)-valine-OH (11.91 g, 35.09 mmol) and HATU (12.89 g, 33.90 mmol) were added, dissolved in a small volume of DMF, followed by DIPEA (12.0 mL, 70.6 mmol) and more DMF. The mixture was agitated for 2 h. Following Fmoc-deprotection the resin was washed two times with DCM and dried *in vacuo*, yielding **5** (38.48 g, 11.71 mmol).

2-Chlorotrityl-L-Phe-L-Pro-L-Val-L-Val-D-His(Trt)-NH$_2$ (**6**)

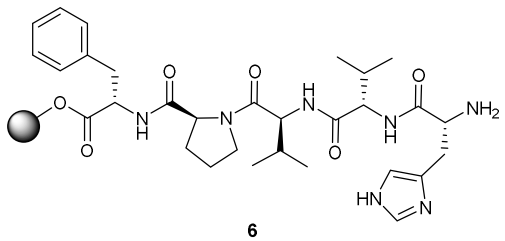

**6**

Resin bound peptide **5** (19.24 g, 5.855 mmol) was swelled in DMF for 10 min. After removal of the solvent, Fmoc-(L)-valine-OH (5.97 g, 17.59 mmol) and HATU (10.89 g, 28.64 mmol)[2] were added, dissolved in a small volume of DMF, followed by DIPEA (6.0 mL, 35 mmol) and more DMF. The mixture was agitated for 2 h. After Fmoc-deprotection, Fmoc-(D)-histidine-(Trt)-OH (10.89 g, 17.57 mmol) and HATU (6.454 g, 16.96 mmol) were added, dissolved in a small amount

[2] Due to miscalculation the equivalents don't adhere to the standard procedure.

of DMF, followed by DIPEA (6.0 mL, 35 mmol) and more DMF. The mixture was agitated for 2 h. Since no complete conversion was observed, the supernatant was drained, the resin was washed three times with DMF and the coupling step was repeated. Fmoc-(D)-histidine-(Trt)-OH (3.618 g, 5.838 mmol) and HATU (2.011 g, 5.289 mmol) were added, dissolved in a small volume of DMF, followed by DIPEA (2000 µL, 11.76 mmol) and more DMF. The mixture was agitated for 1 h, after which complete conversion was observed using UHR-MS. Following Fmoc-deprotection, the resin was washed two times with DCM and dried *in vacuo*, yielding **6** (41.35 g, 5.855 mmol).

2-Chlorotrityl-L-Phe-L-Pro-L-Val-L-Ile-D-His(Trt)-$NH_2$ resin (**7**)

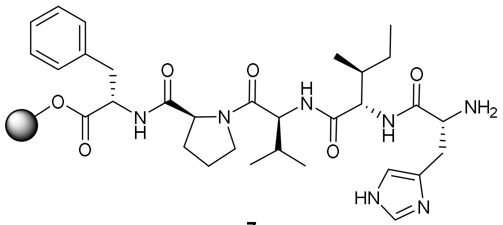

7

**5** (19.24, 5.855 mmol) was swelled in DMF for 20 min. After removal of the solvent, Fmoc-(L)-Ile-OH (6.207 g, 17.56 mmol) and HATU (6.456 g, 16.98 mmol) were added, dissolved in a small volume of DMF, followed by DIPEA (6.0 mL, 35 mmol) and more DMF. The mixture was agitated for 1.5 h. After Fmoc-deprotection, Fmoc-(D)-histidine-(Trt)-OH (10.54g, 17.00 mmol) and HATU (6.440 g, 16.96 mmol) were added, dissolved in a small amount DMF, followed by DIPEA (6.0 mL, 35 mmol) and more DMF. The mixture was agitated for 2 h. Following Fmoc-deprotection the resin was washed two times with DCM and dried *in vacuo*, yielding **7** (21.48 g, 5.855 mmol).

(3-methylbutanoyl)-D-histidyl-L-valyl-L-valyl-L-prolyl-L-phenylalanine (**8**)

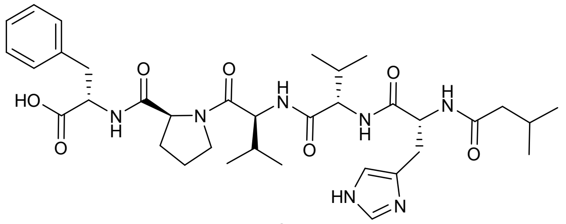

8

**6** (13.00 g, 1.841 mmol) was swelled in DMF for 10 min. After removal of the solvent, HATU (2.150 g, 5.65 mmol) was added, dissolved in a small volume of DMF, followed by isovaleric acid (643 µL, 5.85 mmol), DIPEA (2000 µL, 11.76 mmol) and more DMF. The mixture was agitated for 3 h, after which LC-MS indicated completeness of the reaction. The supernatant was drained, and the resin was washed with DMF (3x) and DMC (2x). The cleavage was done as described in the general method section. The obtained crude was washed four times with ice cold diethyl ether, decanted and dried under reduced pressure. Purification of 48% of the crude using semi preparative HPLC (5-50-95% AcCN + 0.1% FA, NUCLEODUR® C18 Gravity SB, 3 µm, 250 x 10 mm, flow rate: 2 mL/min) yielded **8** as a white waxy film (196 mg, 0.288 mmol, overall yield calculated to be: 33%).

**$^1$H-NMR** (DMSO-$d_6$, 600 MHz): $\delta_H$ [ppm] = 8.03 (d, 1H, $J$ = 8.1 Hz, N$H$ His), 8.03 (d, 1H, $J$ = 8.1 Hz, ε-N$H$ His), 7.99 (d, 1H, $J$ = 7.9 Hz, N$H$ Val 1), 7.98 (d, 1H, $J$ = 7.2 Hz, N$H$ Phe), 7.91 (s, 1H, ε-C$H_{arom}$ His), 7.69 (d, 1H, $J$ = 9.0 Hz, N$H$ Val 2),7.28-7.17 (m, 5H, δ-C$H_{arom}$ Phe), 6.93 (s, 1H, δ-C$H_{arom}$ His), 4.63 (ddd, 1H, $J$ = 8.5, 8.5, 6.0 Hz, α-C$H$ His), 4.38 (ddd, 1H, $J$ = 7.6, 7.6, 5.7 Hz, α-C$H$ Phe), 4.36 (dd, 1H, $J$ = 8.3, 3.7 Hz, α-C$H$ Pro), 4.25 (dd, 1H, $J$ = 8.3, 8.3 Hz, α-C$H$ Val 1), 4.22 (dd, 1H, $J$ = 8.9, 6.1 Hz, α-C$H$ Val), 3.75 (ddd, 1H, $J$ = 9.6, 6.9, 6.9 Hz, δ-C$H_2$N a

Pro), 3.53 (ddd, 1H, *J* = 9.4, 7.1, 5.6 Hz, δ-$CH_2$N b Pro), 3.00 (dd, 1H, *J* = 14.2, 5.8 Hz, β-C$H_2$ a Phe), 2.94 (dd, 1H, *J* = 14.6, 6.5 Hz, β-C$H_2$ a His), 2.92 (dd, 1H, *J* = 13.8, 7.5 Hz, β-C$H_2$ b Phe), 2.77 (dd, 1H, *J* = 13.8, 7.5 Hz, β-C$H_2$ b His), 2.00-1.93 (m, 4H, β-C$H_2$ a Pro, β-C*H* Val, C$H_2$ iVal), 1.93-1.89 (m, 2H, β-C*H* Val, CH iVal), 1.89-1.83 (m, 1H, γ-C$H_2$ a Pro), 1.83-1.75 (m, 2H, β-C$H_2$ b Pro, γ-C$H_2$ b Pro), 0.88 (d, 3H, *J* = 6.8 Hz, γ-C$H_3$ a Val 1), 0.86 (d, 3H, *J* = 6.4 Hz, γ-C$H_3$ b Val 1), 0.81 (d, 3H, *J* = 6.4 Hz, C$H_3$ a iVal), 0.78 (d, 3H, *J* = 6.6 Hz, C$H_3$ b iVal), 0.73 (d, 3H, *J* = 6.6 Hz, γ-C$H_3$ a Val), 0.70 (d, 3H, *J* = 7.0 Hz, γ-C$H_3$ b Val).

**$^{13}$C-NMR** (DMSO-$d_6$, 151 MHz): $\delta_C$ [ppm] = 172.7 (*C*OOH Phe), 171.5 (*C*O iVal), 171.4 (*C*O Pro), 170.7 (*C*O His), 170.6 (*C*O Val 2), 169.7 (*C*O Val 1), 137.3 (γ-$C_{quart}$ Phe), 134.3 (weak, ε-$CH_{arom}$ His), 132.2* (γ-$C_{quart}$ His), 129.1, 128.1, 126.4 (δ-$CH_{arom}$ Phe), 117.0* (δ-$CH_{arom}$ His), 59.0 (α-*C*H Pro), 56.9 (α-*C*H Val 2), 55.8 (α-*C*H Val 1), 53.5 (α-*C*H Phe), 52.3 (α-*C*H His), 47.0 (δ-$CH_2$N Pro), 44.4 ($CH_2$ iVal), 36.7 (β-$CH_2$ a Phe), 30.7 (β-*C*H Val), 29.7 (β-*C*H Val 1), 29.1 (weak, β-$CH_2$ His), 28.9 (β-$CH_2$ Pro), 25.4 (*C*H iVal), 24.3 (γ-$CH_2$ Pro), 22.2 ($CH_3$ a iVal), 22.1 ($CH_3$ b iVal), 19.0 (γ-$CH_3$ a Val 1), 18.4 (γ-$CH_3$ b Val 1), 17.6 (γ-$CH_3$ b Val).

Additional found signals: $\delta_H$ [ppm] = 8.14 (FA), Broad peak between 4.00 to 2.73 ($H_2O$), 3.10, 2.54 (DMSO), 1.18. ε-N*H* His and COO*H* Phe were not observed. $\delta_C$ [ppm] = 45.7, 8.58.

**UHR-MS (ESI-TOF)** *m/z* calcd for $C_{35}H_{52}N_7O_7$: 682.3923 $[M+H]^+$; found: 682.3909 $[M+H]^+$

**Specific rotation** $[\alpha]_D^{21}$ = -11.2° (c = 1.78, $CH_3OH$)

(2-phenylacetyl)-D-histidyl-L-valyl-L-valyl-L-prolyl-L-phenylalanine (**9**)

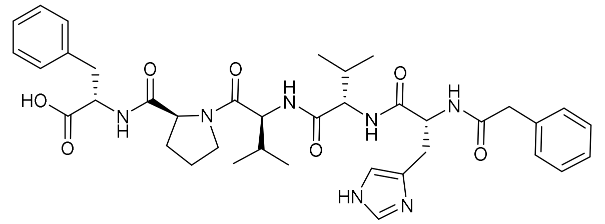

**9**

**6** (13.70 g, 1.940 mmol) was swelled in DMF for 10 min. After removal of the solvent, HATU (2.159 g, 5.68 mmol) was added, dissolved in a small volume of DMF, followed by phenylacetic acid (0.799 g, 5.87 mmol), DIPEA (2000 µL, 11.76 mmol) and more DMF. The mixture was agitated for 3 h, after which LC-MS indicated completeness of the reaction. The supernatant was drained and the resin was washed with DMF (3x), DMC (2x) and MeOH (2x). The cleavage was done as described in the general method section. The obtained crude was washed four times with ice cold diethyl ether, decanted and dried under reduced pressure. Purification of 37% of the crude using semi preparative HPLC (5-50-95% AcCN + 0.1% FA, NUCLEODUR® C18 Gravity SB, 3 µm, 250 x 10 mm, flow rate: 2 mL/min) yielded **9** as a white wax like film (303 mg, 0.423 mmol, overall yield calculated to be: 59%).

**$^1$H-NMR** (DMSO-$d_6$, 600 MHz): $\delta_H$ [ppm] = 8.32 (d, 1H, *J* = 8.3 Hz, N*H* His), 7.98 (d, 1H, *J* = 7.2 Hz, N*H* Phe ), 7.97 (d, 1H, *J* = 7.5 Hz, N*H* Val 1), 7.82 (s, 1H, ε-C$H_{arom}$ His), 7.76 (d, 1H, *J* = 8.9 Hz, N*H* Val 2), 7.28-7.15 (m, 10H, δ-C$H_{arom}$ Phe, C$H_{arom}$ PA), 6.88 (s, 1H, δ-C$H_{arom}$ His), 4.63 (ddd, 1H, *J* = 7.7, 7.7, 6.6 Hz, α-C*H* His), 4.39 (ddd, 1H, J = 7.6, 7.6, 6.0 Hz, α-C*H* Phe), 4.37 (dd, 1H, *J* = 8.5, 3.4 Hz, α-C*H* Pro), 4.26 (dd, 1H, *J* = 8.3, 8.3 Hz, α-C*H* Val 1), 4.22 (dd, 1H, *J* = 9.0, 6.2 Hz, α-C*H* Val 2), 3.76 (ddd, 1H, *J* = 9.8, 6.8, 6.8 Hz, δ-C$H_2$N a Pro), 3.54 (ddd, 1H, *J* = 9.4, 7.2, 5.4 Hz, δ-C$H_2$N b Pro), 3.46 (d 1H, *J* = 14.3 Hz, C$H_2$ a PA), 3.42 (d 1H, *J* = 14.2 Hz, C$H_2$ b PA), 3.00 (dd, 1H, *J* = 14.1, 5.9 Hz, β-C$H_2$ a Phe), 2.95 (dd, 1H, *J* = 14.8, 5.9 Hz, β-C$H_2$ a His),

2.92 (dd, 1H, *J* = 13.9, 7.7 Hz, β-C*H2* b Phe),2.79 (dd, 1H, *J* = 14.8, 8.4 Hz, β-C*H2* b His), 2.00-1.93 (m, 2H, β-C*H2* a Pro, β-C*H* Val 1), 1.93-1.89 (m, 1H, β-C*H* Val 2), 1.89-1.84 (m, 1H, γ-C*H2* a Pro), 1.84-1.76 (m, 2H, β-C*H2* b Pro, γ-C*H2* b Pro), 0.88 (d, 3H, *J* = 6.6 Hz, γ-C*H3* a Val 1), 0.86 (d, 3H, *J* = 6.7 Hz, γ-C*H3* b Val 1), 0.71 (d, 3H, *J* = 6.8 Hz, γ-C*H3* a Val 2), 0.67 (d, 3H, *J* = 6.7 Hz, γ-C*H3* b Val 2).

**$^{13}$C-NMR** (DMSO-*d*6, 151 MHz): $\delta_C$ [ppm] = 172.7 (*C*OOH Phe), 171.5 (*C*O Pro), 170.7 (*C*O His), 170.6 (*C*O Val 2), 170.0 (*C*O PA), 169.7 (*C*O Val 1), 137.3 (γ-$C_{quart}$ Phe), 136.1 ($C_{quart}$ PA), 134.3 (ε-$C$H$_{arom}$ His), 132.3 (weak, γ-$C_{quart}$ His), 129.1, 128.9, 128.1, 128.1, 126.4, 126.2 (δ-$C$H$_{arom}$ Phe, $C$H$_{arom}$ PA), 117.0* (δ-$C$H$_{arom}$ His), 59.0 (α-*C*H Pro), 57.0 (α-*C*H Val 2), 55.9 (α-*C*H Val 1), 53.5 (α-*C*H Phe), 52.6 (α-*C*H His), 47.0 (δ-*C*H$_2$N Pro), 42.0 (*C*H$_2$ PA), 36.7 (β-*C*H$_2$ Phe), 30.6 (β-*C*H Val 2), 29.8 (β-*C*H Val 1), 29.5 (β-*C*H$_2$His), 28.9 (β-*C*H$_2$ Pro), 24.3 (γ-*C*H$_2$ Pro), 19.0 (γ-*C*H$_3$ a Val 2), 19.0 (γ-*C*H$_3$ a Val 1), 18.5 (γ-*C*H$_3$ b Val 1), 17.6 (γ-*C*H$_3$ b Val 2).

Additional found signals: $\delta_H$ [ppm] = 8.14 (FA), 2.54 (DMSO), 3.09, 1.17. ε-N*H* His and COO*H* Phe were not observed. $\delta_C$ [ppm] = 8.62.

**UHR-MS (ESI-TOF)** *m/z* calcd for $C_{38}H_{50}N_7O_7$: 716.3766 [M+H]$^+$; found: 716.3766 [M+H]$^+$

**Specific rotation** $[\alpha]_D^{21}$ = -12.4° (c = 1.61, $CH_3OH$)

(3-methylbutanoyl)-D-histidyl-L-isoleucyl-L-valyl-L-prolyl-L-phenylalanine (**10**)

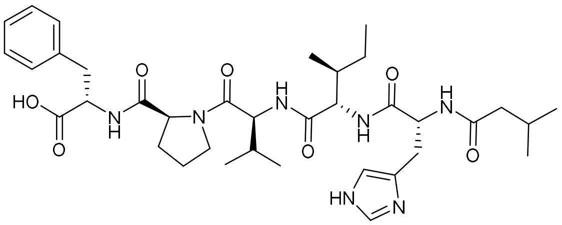

10

**7** (8.090 g, 2.225 mmol) was swelled in DMF for 30 min. After removal of the solvent, HATU (2.425 g, 6.378 mmol) was added, dissolved in a small volume of DMF, followed by isovaleric acid (724 µL, 6.59 mmol), DIPEA (2300 µL, 13.53 mmol) and more DMF. The mixture was agitated for 2 h, after which LC-MS indicated completeness of the reaction. The supernatant was drained, and the resin was washed with DMF (3x), DMC (2x) and MeOH (2x). The cleavage was done as described in the general method section. The obtained crude was washed four times with ice cold diethyl ether, decanted and dried under reduced pressure. Purification of 35% of the crude using semi preparative HPLC (5-50-95% AcCN + 0.1% FA, NUCLEODUR® C18 Gravity SB, 3 µm, 250 x 10 mm, flow rate: 2 mL/min) yielded **10** as a white wax like film (87.8 mg, 0.126 mmol, overall yield calculated to be: 16%).

**$^1$H-NMR** (DMSO-*d*6, 600 MHz): $\delta_H$ [ppm] = 8.23 (s, 1H, ε-C*H*$_{arom}$ His), 8.06 (d, 1H, *J* = 8.1 Hz, N*H* His), 8.00 (d, 1H, *J* = 8.4 Hz, N*H* Val), 7.98 (d, 1H, *J* = 7.7 Hz, N*H* Phe), 7.73 (d, 1H, *J* = 9.2 Hz, N*H* Ile), 7.28-7.18 (m, 5H, δ-C*H*$_{arom}$ Phe), 7.05 (s, 1H, δ-C*H*$_{arom}$ His), 4.66 (ddd, 1H, *J* = 8.1, 8.1, 6.5 Hz, α-C*H* His), 4.39 (ddd, 1H, *J* = 7.3, 7.3, 6.2 Hz, α-C*H* Phe), 4.36 (dd, 1H, *J* = 8.3, 3.9 Hz, α-C*H* Pro), 4.26 (dd, 1H, *J* = 8.3, 8.3 Hz, α-C*H* Val), 4.24 (dd, 1H, *J* = 8.5, 7.5 Hz, α-C*H* Ile), 3.77-3.71 (m, 1H, δ-C*H2*N a Pro), 3.56-3.51 (m, 1H, δ-C*H2*N b Pro), 3.00 (dd, 1H, *J* = 13.9, 5.9 Hz, β-C*H2* a Phe), 2.97 (dd, 1H, *J* = 15.3, 6.3 Hz, β-C*H2* a His), 2.92 (dd, 1H, *J* = 13.9, 7.9 Hz, β-C*H2* b Phe), 2.79 (dd, 1H, *J* = 15.0, 8.6 Hz, β-C*H2* b His), 2.00-1.92 (m, 4H, C*H2* iVal, β-C*H* Val, β-C*H2* a Pro ), 1.92-1.83 (m, 2H, C*H* iVal, γ-C*H2* a Pro), 1.83-1.75 (m, 2H, β-C*H2* b Pro, γ-C*H2* b Pro), 1.69-1.62 (m, 1H, β-C*H* Ile), 1.29-1.22 (m, 1H, γ-C*H2* a Ile), 0.97-0.89 (m, 1H, γ-C*H2* b Ile), 0.87 (d, 3H, *J* = 6.8 Hz, γ-C*H*$_3$ a Val), 0.85 (d, 3H, *J* = 6.4 Hz, γ-C*H*$_3$ b Val), 0.81 (d, 3H, *J* = 6.6 Hz, C*H*$_3$ a iVal), 0.78 (d, 3H, *J* = 6.6 Hz, C*H*$_3$ b iVal), 0.74 (t, 3H, *J* = 7.4 Hz, δ-C*H*$_3$ Ile), 0.71 (d, 3H, *J* = 7.0 Hz, γ-C*H*$_3$ Ile).

**$^{13}$C-NMR** (DMSO-*d*$_6$, 151 MHz): $\delta_C$ [ppm] = 172.7 (COOH Phe), 171.5 (CO iVal), 171.4 (CO Pro), 170.7 (CO Ile), 170.4 (CO His), 169.6 (CO Val), 137.3 (γ-C$_{quart}$ Phe), 134.1 (ε-CH$_{arom}$ His), 131.3 (weak, γ-C$_{quart}$ His), 129.1, 128.1, 126.4 (δ-CH$_{arom}$ Phe), 116.9 (weak, δ-CH$_{arom}$ His), 59.0 (α-CH Pro), 56.2 (α-CH Ile), 55.8 (α-CH Val), 53.5 (α-CH Phe), 52.0 (α-CH His), 47.0 (δ-CH$_2$N Pro), 44.4 (CH$_2$ iVal), 36.9 (β-CH Ile), 36.7 (β-CH$_2$ Phe), 29.8 (β-CH Val), 28.9 (β-CH$_2$ Pro), 28.6 (β-CH$_2$ His), 25.4 (CH iVal), 24.3 (γ-CH$_2$ Pro), 24.0 (γ-CH$_2$ a Ile), 22.2 (CH$_3$ a iVal), 22.1 (CH$_3$ b iVal), 19.0 (γ-CH$_3$ a Val), 18.4 (γ-CH$_3$ b Val), 15.2 (γ-CH$_3$ Ile), 11.0 (δ-CH$_3$ Ile).

Additional found signals: $\delta_H$ [ppm] = 8.14 (FA), Broad peak between 4.00 to 2.60 ($H_2O$), 3.10, 2.54 (DMSO), 1.18. ε-N*H* His and COO*H* Phe were not observed. $\delta_C$ [ppm] = 163.0 (FA), 157.7, 45.7, 8.60.

**UHR-MS (ESI-TOF)** *m/z* calcd for $C_{36}H_{54}N_7O_7$: 696.4079 $[M+H]^+$; found: 696.4082 $[M+H]^+$

**Specific rotation** $[\alpha]_D^{21}$ = -34.0° (c = 2.35, $CH_3OH$)

(2-phenylacetyl)-D-histidyl-L-isoleucyl-L-valyl-L-prolyl-L-phenylalanine (**11**)

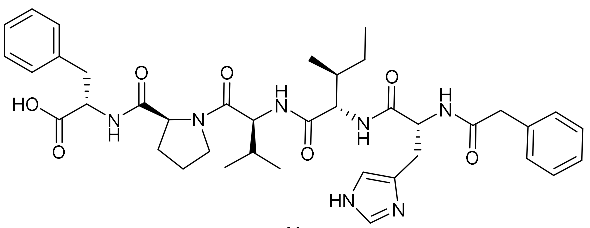

**11**

**7** (7.994 g, 2.166 mmol) were swelled in DMF for 30 min. After removal of the solvent, HATU (2.425 g, 6.378 mmol) was added, dissolved in a small amount of DMF, followed by phenylacetic acid (0.897 g, 6.588 mmol), DIPEA (2300 µL, 13.53 mmol) and additional DMF. The mixture was agitated for 2 h, after which LC-MS indicated completeness of the reaction. The supernatant was drained and the resin was washed with DMF (3x), DMC (2x) and MeOH (2x). The cleavage was done as described in the general method section. The obtained crude was washed four times with ice cold diethyl ether, decanted and dried under reduced pressure. Purification of 40% of the crude using semi preparative HPLC (5-50-95% AcCN + 0.1% FA, NUCLEODUR® C18 Gravity SB, 3 µm, 250 x 10 mm, flow rate: 2 mL/min) yielded **11** as a white wax like film (182 mg, 0.249 mmol, overall yield calculated to be: 29%).

**$^1$H-NMR** (DMSO-*d*$_6$, 600 MHz): $\delta_H$ [ppm] = 8.35 (d, 1H, *J* = 8.1 Hz, N*H* His), 8.20 (s, 1H, ε-C*H*$_{arom}$ His), 7.99 (d, 1H, *J* = 7.5 Hz, N*H* Phe), 7.98 (d, 1H, *J* = 8.0 Hz, N*H* Val), 7.82 (d, 1H, *J* = 8.6 Hz, N*H* Ile), 7.28-7.13 (m, 10H, δ-C*H*$_{arom}$ Phe, C*H*$_{arom}$ PA), 7.02 (s, 1H, δ-C*H*$_{arom}$ His), 4.66 (ddd, 1H, *J* = 8.2, 8.2, 6.4 Hz, α-C*H* His), 4.39 (ddd, 1H, *J* = 7.8, 7.8, 5.9 Hz, α-C*H* Phe), 4.37 (dd, 1H, *J* = 8.7, 4.1 Hz, α-C*H* Pro), 4.27 (dd, 1H, *J* = 8.3, 8.3 Hz, α-C*H* Val), 4.23 (dd, 1H, *J* = 8.8, 7.0 Hz, α-C*H* Ile), 3.78-3.72 (m, 1H, δ-C*H*$_2$N a Pro), 3.57-3.51 (m, 1H, δ-C*H*$_2$N b Pro), 3.45 (d, 1H, *J* = 14.2 Hz, C*H*$_2$ a PA), 3.42 (d, 1H, *J* = 14.2 Hz, C*H*$_2$ b PA), 3.00 (dd, 1H, *J* = 13.8, 5.5 Hz, β-C*H*$_2$ a Phe), 2.98 (dd, 1H, *J* = 14.5, 5.9 Hz, β-C*H*$_2$ a His), 2.92 (dd, 1H, *J* = 13.9, 7.9 Hz, β-C*H*$_2$ b Phe), 2.82 (dd, 1H, *J* = 14.9, 8.3 Hz, β-C*H*$_2$ b His), 2.00-1.92 (m, 2H, β-C*H* Val, β-C*H*$_2$ a Pro), 1.90-1.84 (m, 1H, γ-C*H*$_2$ a Pro), 1.84-1.75 (m, 2H, β-C*H*$_2$ b Pro, γ-C*H*$_2$ b Pro), 1.68-1.61 (m, 1H, β-C$H$ Ile), 1.26-1.18 (m, 1H, γ-C$H_2$ a Ile), 0.94-0.82 (m, 1H, γ-C$H_2$ b Ile), 0.88 (d, 3H, $J$ = 6.4 Hz, γ-C$H_3$ a Val), 0.86 (d, 3H, $J$ = 6.6 Hz, γ-C$H_3$ b Val), 0.73 (t, 3H, $J$ = 7.4 Hz, δ-C$H_3$ Ile), 0.70 (d, 3H, $J$ = 7.0 Hz, γ-C$H_3$ Ile).

**$^{13}$C-NMR** (DMSO-$d_6$, 151 MHz): $\delta_C$ [ppm] = 172.7 (*C*OOH Phe), 171.4 (*C*O Pro), 170.6 (*C*O Ile), 170.2 (*C*O His), 170.1 (*C*O PA), 169.7 (*C*O Val), 137.3 (γ-$C_{quart}$ Phe), 136.1 ($C_{quart}$ PA), 134.1 (ε-$C$H$_{arom}$ His), 131.3* (γ-$C_{quart}$ His), 129.1, 128.9, 128.1, 128.0, 126.4, 126.2 (δ-$C$H$_{arom}$ Phe, $C$H$_{arom}$ PA), 116.8 (δ-$C$H$_{arom}$ His), 59.0 (α-*C*H Pro), 56.3 (α-*C*H Ile), 55.8 (α-*C*H Val), 53.5 (α-*C*H Phe), 52.2 (α-*C*H His), 47.0 (δ-$C$H$_2$N Pro), 42.0 ($C$H$_2$ PA), 36.8 (β-*C*H Ile), 36.7 (β-$C$H$_2$ Phe), 29.7 (β-*C*H Val), 28.9 (β-$C$H$_2$ Pro), 28.8 (β-$C$H$_2$ His), 24.3 (γ-$C$H$_2$ Pro), 23.9 (γ-$C$H$_2$ a Ile), 19.0 (γ-$C$H$_3$ a Val), 18.5 (γ-$C$H$_3$ b Val), 15.2 (γ-$C$H$_3$ Ile), 11.0 (δ-$C$H$_3$ Ile).

Additional found signals: $\delta_H$ [ppm] = 8.14 (FA), Broad peak between 4.00 to 2.32 ($H_2O$), 2.54 (DMSO). ε-N$H$ His and COO$H$ Phe were not observed. $\delta_C$ [ppm] = 163.0 (FA).

**UHR-MS (ESI-TOF)** *m/z* calcd for $C_{39}H_{52}N_7O_7$: 730.3923 [M+H]$^+$; found: 730.3924 [M+H]$^+$

**Specific rotation** $[\alpha]_D^{21}$ = -27.6° (c = 1.45, $CH_3OH$)

(*S*)-*N*-((*S*)-1-hydroxy-3-phenylpropan-2-yl)-1-((3-methylbutanoyl)-D-histidyl-L-valyl-L-valyl)pyrrolidine-2-carboxamide (**12**)

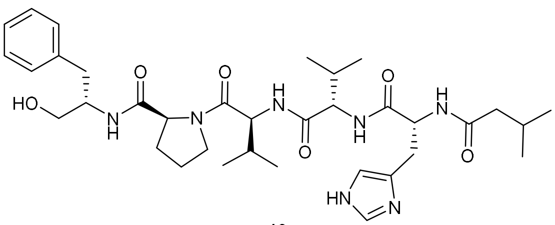

12

Peptide **8** (87.5 mg, 0.128 mmol) was dissolved in anhydrous MeOH (20 mL) and thionyl chloride (14.0 µL, 0.192 mmol) was added at 0 °C. The mixture was stirred under a gentle reflux for 3.5 h and then concentrated *in vacuo*. The crude was suspended in anhydrous THF and cooled to 0 °C. $LiBH_4$ (10.5 mg, 0.482 mmol) was added and the reaction was stirred at room temperature. The reaction progress was monitored using LC-MS and after 2 h more $LiBH_4$ (6.0 mg, 0.28 mmol) was added at 0 °C. After a total of 3 h the reaction was quenched using sat. $NH_4Cl$ solution. Water was added until all formed salts were dissolved. The layers were separated and the aqueous layer was extracted with ethyl acetate three times. The combined organic phases were washed once with brine, dried over $MgSO_4$, filtered and the solvent was evaporated under reduced pressure. Purification of the crude using semi preparative HPLC (5-40-95% AcCN + 0.1% FA, NUCLEODUR® C18 Gravity SB, 3 µm, 250 x 10 mm, flow rate: 2 mL/min) yielded **12** as a colorless solid (63.9 mg, 0.0957 mmol, 75% over 2 steps).

**$^1$H-NMR** (DMSO-$d_6$, 600 MHz): $\delta_H$ [ppm] = 7.98 (d, 1H, $J$ = 7.9 Hz, N$H$ Val 1), 7.98 (d, 1H, $J$ = 7.9 Hz, N$H$ His), 7.66 (d, 1H, $J$ = 9.2 Hz, N$H$ Val 2), 7.51 (s, 1H, ε-C$H_{arom}$ His), 7.51 (d, 1H, $J$ = 6.1 Hz, N$H$ Phe), 7.27-7.14 (m, 5H, δ-C$H_{arom}$ Phe), 6.77 (s, 1H, δ-C$H_{arom}$ His), 4.59 (ddd, 1H, $J$ = 7.9, 7.9, 6.4 Hz, α-C$H$ His), 4.30-4.25(m, 2H, α-C$H$ Val 1, α-C$H$ Pro), 4.23 (dd, 1H, $J$ = 8.2, 6.8 Hz, α-C$H$ Val 2), 3.87-3.80 (m, 1H, α-C$H$ Phe), 3.76-7.70 (m, 1H, δ-C$H_2$N a Pro), 3.57-3.51 (m, 1H, δ-C$H_2$N b Pro), 3.27 (t, 2H, $J$ = 5.2 Hz, C$H_2$OH Phe), 2.90 (dd, 1H, $J$ = 14.9, 5.9 Hz, β-C$H_2$ a His), 2.79 (dd, 1H, $J$ = 13.7, 6.5 Hz, β-C$H_2$ a Phe), 2.73 (dd, 1H, $J$ = 14.8, 8.9 Hz, β-C$H_2$ b His), 2.68 (dd, 1H, $J$ = 13.7, 7.1 Hz, β-C$H_2$ b Phe), 2.03-1.97 (m, 1H, β-C$H$ Val 1), 1.96 (d, 2H, $J$ = 6.6 Hz, $CH_2$ iVal), 1.97-1.84 (m, 4H, β-C$H$ Val 2, β-C$H_2$ a Pro, C$H$ iVal, γ-C$H_2$ a Pro), 1.81-

1.76 (m, 1H, γ-C$H_2$ b Pro), 1.76-1.71 (m, 1H, β-C$H_2$ b Pro), 0.90 (d, 3H, $J$ = 6.8 Hz, γ-C$H_3$ a Val 1), 0.88 (d, 3H, $J$ = 6.6 Hz, γ-C$H_3$ b Val 1), 0.81 (d, 3H, $J$ = 6.4 Hz, C$H_3$ a iVal), 0.78 (d, 3H, $J$ = 6.2 Hz, C$H_3$ b iVal), 0.73 (d, 3H, $J$ = 6.8 Hz, γ-C$H_3$ a Val 2), 0.70 (d, 3H, $J$ = 6.8 Hz, γ-C$H_3$ b Val 2).

**$^{13}$C-NMR** (DMSO-$d_6$, 151 MHz): $\delta_C$ [ppm] = 171.4 (*C*O iVal), 171.1 (*C*O Pro), 171.1 (*C*O His), 170.7 (*C*O Val 2), 169.7 (*C*O Val 1), 138.9 (γ-$C_{quart}$ Phe), 134.5 (weak, ε-*C*$H_{arom}$ His), 129.1, 128.0, 125.8 (δ-*C*$H_{arom}$ Phe), 61.8 (*C*$H_2$OH Phe), 59.5 (α-*C*H Pro), 57.0 (α-*C*H Val 2), 55.9 (α-*C*H Val 1), 52.7 (α-*C*H His), 52.2 (α-*C*H Phe), 47.0 (δ-*C*$H_2$N Pro), 44.5 (*C*$H_2$ iVal), 36.3 (β-*C*$H_2$ Phe), 30.6 (β-*C*H Val 2), 29.7 (β-*C*H Val 1), 29.8* (β-*C*$H_2$ a His), 29.1 (β-*C*$H_2$ Pro), 25.4 (*C*H iVal), 24.4 (γ-*C*$H_2$ Pro), 22.2 (*C*$H_3$ a iVal), 22.1 (*C*$H_3$ b iVal), 19.0 (γ-*C*$H_3$ a Val 2), 19.1 (γ-*C*$H_3$ a Val 1), 18.4 (γ-*C*$H_3$ b Val 1), 17.6 (γ-*C*$H_3$ b Val 2).

Additional found signals: $\delta_H$ [ppm] = 8.15 (FA), Broad peak between 4.00 to 2.58 ($H_2O$), 2.54 (DMSO). ε-N*H* His was not observed. $\delta_C$ [ppm] = δ-*C*$H_{arom}$ His and γ-$C_{quart}$ His were not observed.

**UHR-MS (ESI-TOF)** *m/z* calcd for $C_{35}H_{54}N_7O_6$: 668.4130 $[M+H]^+$; found: 668.4126 $[M+H]^+$

**Specific rotation** $[\alpha]_D^{21}$ = -28.7° (c = 0.87, $CH_3OH$)

(*S*)-*N*-((*S*)-1-hydroxy-3-phenylpropan-2-yl)-1-((2-phenylacetyl)-D-histidyl-L-valyl-L-valyl)pyrrolidine-2-carboxamide (**13**)

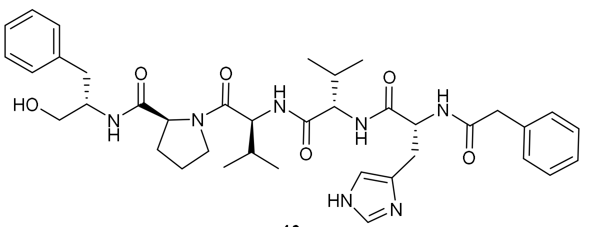

13

To a solution of peptide **9** (118.1 mg, 0.1650 mmol) in anhydrous MeOH (40 mL) was added thionyl chloride (18.0 µL, 0.247 mmol) at 0 °C. The mixture was stirred under a gentle reflux for 3.5 h and then concentrated *in vacuo*. The crude product (107.3 mg, 0.1470 mmol, 89%) was directly used in the next step. For testing the conditions the crude was split into two parts, the syntheses were carried out identical.

The first part of the crude methyl ester (49.5 mg, 0.0678 mmol) was suspended in anhydrous THF and cooled to 0 °C. $LiBH_4$ (5.8 mg, 0.27 mmol) was added and the reaction was stirred at room temperature. The reaction progress was monitored using LC-MS and after 1 h more $LiBH_4$ (3.1 mg, 0.14 mmol) was added at 0 °C. After a total of 3 h the reaction was quenched using sat. $NH_4Cl$ solution. Water was added until all formed salts were dissolved. The layers were separated and the aqueous layer was extracted with ethyl acetate three times. The combined organic phases were washed once with brine, dried over $MgSO_4$, filtered and the solvent was evaporated under reduced pressure.

The second part of the crude methyl ester (68.5 mg, 0.0939 mmol) was suspended in anhydrous THF and cooled to 0 °C. $LiBH_4$ (8.3 mg, 0.38 mmol) was added and the reaction was stirred at room temperature. The reaction progress was monitored using LC-MS and after 1.5 h more $LiBH_4$ (4.1 mg, 0.19 mmol) was added at 0 °C. After a total of 2 h the reaction was quenched using sat. $NH_4Cl$ solution. Water was added until all formed salts were dissolved. The layers were separated and the aqueous layer was extracted with ethyl acetate three times. The combined organic phases were washed once with brine, dried over $MgSO_4$, filtered and the solvent was evaporated under reduced pressure.

Both crudes were combined for purification using semi preparative HPLC (5-40-95% AcCN + 0.1% FA, NUCLEODUR® C18 Gravity SB, 3 µm, 250 x 10 mm, flow rate: 2 mL/min), which yielded **13** as a colorless solid (72.8 mg, 0.104 mmol, 63% over 2 steps).

**$^{1}$H-NMR** (DMSO-*d*$_6$, 600 MHz): $\delta_H$ [ppm] = 8.29 (d, 1H, *J* = 7.9 Hz, N*H* His), 7.96 (d, 1H, *J* = 8.1 Hz, N*H* Val 1), 7.75 (d, 1H, *J* = 9.7 Hz, N*H* Val 2), 7.57 (s, 1H, ε-C*H*$_{arom}$ His), 7.51 (d, 1H, *J* = 8.4 Hz, N*H* Phe), 7.27-7.14 (m, 10H, δ-C*H*$_{arom}$ Phe, C*H*$_{arom}$ PA), 6.79 (s, 1H, δ-C*H*$_{arom}$ His), 4.60 (ddd, 1H, *J* = 7.9, 7.9, 6.6 Hz, α-C*H* His), 4.29-4.25 (m, 2H, α-C*H* Pro, α-C*H* Val 1), 4.22 (dd, 1H, *J* = 9.0, 6.2 Hz, α-C*H* Val 2), 3.87-3.80 (m, 1H, α-C*H* Phe), 3.76-3.70 (m, 1H, δ-C*H*$_2$N a Pro), 3.57-3.52 (m, 1H, δ-C*H*$_2$N b Pro), 3.46 (d 1H, *J* = 14.1 Hz, C*H*$_2$ a PA), 3.42 (d 1H, *J* = 14.3 Hz, C*H*$_2$ b PA), 3.28 (t, 2H, *J* = 5.0 Hz, C*H*$_2$OH Phe), 2.92 (dd, 1H, *J* = 14.7, 6.2 Hz, β-C*H*$_2$ a His), 2.79 (dd, 1H, *J* = 13.4, 6.2 Hz, β-C*H*$_2$ a Phe), 2.77 (dd, 1H, *J* = 14.2, 8.7 Hz, β-C*H*$_2$ b His), 2.68 (dd, 1H, *J* = 13.8, 7.1 Hz, β-C*H*$_2$ b Phe), 2.03-1.95 (m, 1H, β-C*H* Val 1), 1.95-1.84 (m, 3H, γ-C*H*$_2$ a Pro, β-C*H*$_2$ a Pro, β-C*H* Val 2), 1.80-1.71 (m, 2H, γ-C*H*$_2$ b Pro, β-C*H*$_2$ b Pro), 0.90 (d, 3H, *J* = 6.8 Hz, γ-C*H*$_3$ a Val 1), 0.88 (d, 3H, *J* = 6.8 Hz, γ-C*H*$_3$ b Val 1), 0.71 (d, 3H, *J* = 7.0 Hz, γ-C*H*$_3$ a Val 2), 0.67 (d, 3H, *J* = 6.8 Hz, γ-C*H*$_3$ b Val2).

**$^{13}$C-NMR** (DMSO-*d*$_6$, 151 MHz): $\delta_C$ [ppm] = 171.1 (*C*O Pro),170.9 (*C*O His), 170.7 (*C*O Val 2), 170.0 (*C*O PA), 169.8 (*C*O Val 1), 138.9 (γ-*C*$_{quart}$ Phe), 136.2 (*C*$_{quart}$ PA), 134.5 (ε-*C*H$_{arom}$ His), 132.9* (γ-*C*$_{quart}$ His), 129.1, 129.0, 128.0, 126.2, 125.8 (δ-*C*H$_{arom}$ Phe, *C*H$_{arom}$ PA), 117.0* (δ-*C*H$_{arom}$ His), 61.8 (*C*H$_2$OH Phe), 59.5 (α-*C*H Pro), 57.1 (α-*C*H Val 2), 56.0 (α-*C*H Val 1), 52.9 (α-*C*H His), 52.2 (α-*C*H Phe), 47.0 (δ-*C*H$_2$N Pro), 42.0 (*C*H$_2$ PA), 36.3 (β-*C*H$_2$ Phe), 30.5 (β-*C*H Val 2), 29.9 (β-*C*H$_2$ His), 29.7 (β-*C*H Val 1), 29.1 (β-*C*H$_2$ Pro), 24.4 (γ-*C*H$_2$ a Pro), 19.1 (γ-*C*H$_3$ a Val 1), 19.0 (γ-*C*H$_3$ a Val 2), 18.5 (γ-*C*H$_3$ b Val 1), 17.6 (γ-*C*H$_3$ b Val2).

Additional found signals: $\delta_H$ [ppm] = 8.15 (FA), 3.66, 3.17 (MeOH), 2.54 (DMSO). ε-N*H* His was not observed. $\delta_C$ [ppm] = 163.1 (FA), 48.6 (MeOH), 40.4 (DMSO).

**UHR-MS (ESI-TOF)** *m/z* calcd for $C_{38}H_{52}N_7O_6$: 702.3974 $[M+H]^+$; found: 702.3975$[M+H]^+$

**Specific rotation** $[\alpha]_D^{21}$ = -56.1° (c = 1.07, $CH_3OH$)

(*S*)-*N*-((*S*)-1-hydroxy-3-phenylpropan-2-yl)-1-((3-methylbutanoyl)-D-histidyl-L-isoleucyl-L-valyl)pyrrolidine-2-carboxamide (**14**)

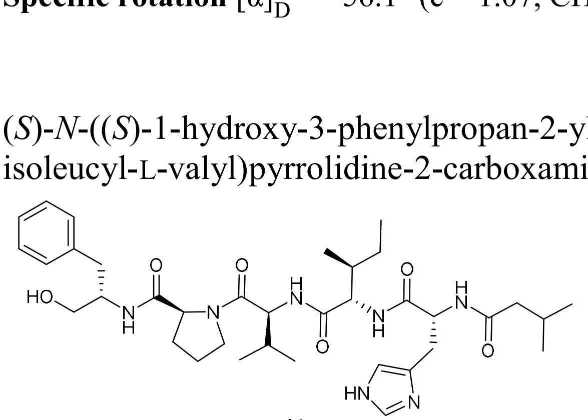

14

The peptide **10** (93.8 mg, 0.135 mmol) was dissolved in anhydrous MeOH (20 mL) and thionyl chloride (14.8 µL, 0.203 mmol) was added at 0 °C. The mixture was stirred under a gentle reflux for 3 h and then concentrated *in vacuo*. The crude product was suspended in anhydrous THF and cooled to 0 °C. $LiBH_4$ (12.6 mg, 0.579 mmol) was added and the reaction was stirred at room temperature. The reaction progress was monitored using LC-MS and after 1.5 h more $LiBH_4$ (6.4 mg, 0.29 mmol) was added at 0 °C. After additional 4 h more $LiBH_4$ (5.0 mg, 0.23 mmol) was added at 0 °C. After a total of 6.5 h the reaction was quenched using sat. $NH_4Cl$ solution. Water was added until all formed salts were dissolved. The layers were separated and the aqueous layer was extracted with ethyl acetate three times. The combined organic phases were washed once with brine, dried over $MgSO_4$, filtered and the solvent was evaporated under reduced pressure. Purification of the crude product using semi preparative HPLC (5-45-95% AcCN + 0.1% FA, NUCLEODUR® C18 Gravity SB, 3 µm, 250 x 10 mm, flow rate: 2 mL/min) yielded **14** as a colorless solid (22.9 mg, 0.0336 mmol, 25% over 2 steps).

**$^1$H-NMR** (DMSO-*d*6, 600 MHz): $\delta_H$ [ppm] = 7.98 (d, 1H, *J* = 8.3 Hz, N*H* Val), 7.98 (d, 1H, *J* = 8.3 Hz, N*H* His), 7.68 (d, 1H, *J* = 9.4 Hz, N*H* Ile), 7.57 (s, 1H, ε-C*H*arom His), 7.51 (d, 1H, *J* = 8.3 Hz, N*H* Phe), 7.28-7.14 (m, 5H, δ-C*H*arom Phe), 6.79 (s, 1H, δ-C*H*arom His), 4.59 (ddd, 1H, *J* = 8.0, 8.0, 6.5 Hz, α-C*H* His), 4.28 (dd, 1H, *J* = 7.6, 7.6 Hz, α-C*H* Val), 4.26 (dd, 1H, *J* = 8.0, 4.7 Hz, α-C*H* Pro), 4.23 (dd, 1H, *J* = 8.8, 7.0 Hz, α-C*H* Ile), 3.87-3.80 (m, 1H, α-C*H* Phe), 3.74-3.69 (m, 1H, δ-C*H2*N a Pro), 3.57-3.51 (m, 1H, δ-C*H2*N b Pro), 3.27 (t, 2H, *J* = 5.4 Hz, C*H2*OH Phe), 2.89 (dd, 1H, *J* = 14.8, 6.0 Hz, β-C*H2* a His), 2.79 (dd, 1H, *J* = 13.7, 6.5 Hz, β-C*H2* a Phe), 2.73 (dd, 1H, *J* = 14.8, 8.5 Hz, β-C*H2* b His), 2.68 (dd, 1H, *J* = 13.9, 7.1 Hz, β-C*H2* b Phe), 2.03-1.97 (m, 1H, β-C*H* Val), 1.95 (d, 2H, *J* = 6.8 Hz, C*H2* iVal), 1.94-1.84 (m, 3H, β-C*H2* a Pro, γ-C*H2* a Pro, C*H* iVal), 1.80-1.71 (m, 2H, β-C*H2* b Pro, γ-C*H2* b Pro), 1.68-1.62 (m, 1H, β-C*H* Ile), 1.31-1.23 (m, 1H, γ-C*H2* a Ile), 0.97-0.91 (m, 1H, γ-C*H2* b Ile), 0.89 (d, 3H, *J* = 6.8 Hz, γ-C*H3* a Val), 0.87 (d, 3H, *J* = 6.8 Hz, γ-C*H3* b Val), 0.81 (d, 3H, *J* = 6.4 Hz, C*H3* a iVal), 0.78 (d, 3H, *J* = 6.6 Hz, C*H3* b iVal), 0.74 (t, 3H, *J* = 7.5 Hz, δ-C*H3* Ile), 0.72 (d, 3H, *J* = 6.8 Hz, γ-C*H3* Ile).

**$^{13}$C-NMR** (DMSO-*d*6, 151 MHz): $\delta_C$ [ppm] = 171.4 (*C*O iVal), 171.0 (*C*O Pro), 170.9 (*C*O His), 170.8 (*C*O Ile), 169.7 (*C*O Val), 138.9 (γ-*C*quart Phe), 134.5 (ε-*C*Harom His), 133.0* (γ-*C*quart His), 129.1, 128.0, 125.8 (δ-*C*Harom Phe), 116.9* (δ-*C*Harom His), 61.8 (*C*H2OH Phe), 59.5 (α-*C*H Pro), 56.3 (α-*C*H Ile), 55.9 (α-*C*H Val), 52.6 (α-*C*H His), 52.2 (α-*C*H Phe), 47.0 (δ-*C*H2N Pro), 44.5 (*C*H2 iVal), 36.8 (β-*C*H Ile), 36.3 (β-*C*H2 a Phe), 29.8 (β-*C*H Val), 29.7 (β-*C*H2 a His), 29.1 (β-*C*H2 Pro), 25.5 (*C*H iVal), 24.4 (γ-*C*H2 Pro), 24.0 (γ-*C*H2 a Ile), 22.2 (*C*H3 a iVal), 22.1 (*C*H3 b iVal), 19.1 (γ-*C*H3 a Val), 18.4 (γ-*C*H3 b Val), 15.3 (γ-*C*H3 Ile), 11.0 (δ-*C*H3 Ile).

Additional found signals: $\delta_H$ [ppm] = 8.15 (FA), 3.17 (MeOH), 2.54 (DMSO). ε-N*H* His was not observed. $\delta_C$ [ppm] = /

**UHR-MS (ESI-TOF)** *m/z* calcd for $C_{36}H_{56}N_7O_6$: 682.4287 $[M+H]^+$; found: 682.4290 $[M+H]^+$

**Specific rotation** $[\alpha]_D^{21}$ = -41.7° (c = 1.20, $CH_3OH$)

(*S*)-*N*-((*S*)-1-hydroxy-3-phenylpropan-2-yl)-1-((2-phenylacetyl)-D-histidyl-L-isoleucyl-L-valyl)pyrrolidine-2-carboxamide (**4**)

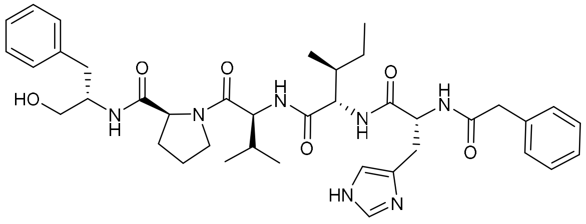

4

To a solution of peptide **11** (160.0 mg, 0.2192 mmol) in anhydrous MeOH (20 mL) was added thionyl chloride (24.0 µL, 0.329 mmol) at 0 °C. The mixture was stirred under a gentle reflux for 3 h and then concentrated *in vacuo*. The crude was suspended in anhydrous THF and cooled to 0 °C. $LiBH_4$ (19.4 mg, 0.891 mmol) was added and the reaction was stirred at room temperature. The reaction progress was monitored using LC-MS and after 2.5 h more $LiBH_4$ (9.3 mg, 0.43 mmol) was added at 0 °C. After a total of 5.5 h the reaction was quenched using sat. $NH_4Cl$ solution. Water was added until all formed salts were dissolved. The layers were separated and the aqueous layer was extracted with ethyl acetate three times. The combined organic phases were washed once with brine, dried over $MgSO_4$, filtered and the solvent was evaporated under reduced pressure. Purification of the crude product using semi preparative HPLC (5-40-95% AcCN + 0.1% FA, NUCLEODUR® C18 Gravity SB, 3 µm, 250 x 10 mm, flow rate: 2 mL/min) yielded **4** as a colorless solid (45.1 mg, 0.0630 mmol, 25% over 2 steps).

**$^{1}$H-NMR** (DMSO-$d_6$, 600 MHz): $\delta_H$ [ppm] = 8.27 (d, 1H, $J$ = 7.9 Hz, N$H$ His), 7.95 (d, 1H, $J$ = 8.3 Hz, N$H$ Val), 7.75 (d, 1H, $J$ = 8.8 Hz, N$H$ Ile), 7.53 (s, 1H, ε-C$H_{arom}$ His), 7.50 (d, 1H, $J$ = 8.4 Hz, N$H$ Phe), 7.27-7.14 (m, 10H, δ-C$H_{arom}$ Phe, C$H_{arom}$ PA), 6.77 (s, 1H, δ-C$H_{arom}$ His), 4.59 (ddd, 1H, $J$ = 7.9, 7.9, 6.6 Hz, α-C$H$ His), 4.27 (dd, 1H, $J$ = 8.1, 8.1 Hz, α-C$H$ Val), 4.27 (dd, 1H, $J$ = 8.1, 3.9 Hz, α-C$H$ Pro), 4.21 (dd, 1H, $J$ = 8.6, 7.2 Hz, α-C$H$ Ile), 3.87-3.80 (m, 1H, α-C$H$ Phe), 3.74-3.69 (m, 1H, δ-C$H_2$N a Pro), 3.57-3.51 (m, 1H, δ-C$H_2$N b Pro), 3.45 (d, 1H, $J$ = 13.9 Hz, C$H_2$ a PA), 3.42 (d, 1H, $J$ = 14.1 Hz, C$H_2$ b PA), 3.27 (t, 2H, $J$ = 5.4 Hz, C$H_2$OH Phe), 2.90 (dd, 1H, $J$ = 14.6, 6.1 Hz, β-C$H_2$ a His), 2.79 (dd, 1H, $J$ = 13.5, 6.3 Hz, β-C$H_2$ a Phe), 2.75 (dd, 1H, $J$ = 14.4, 7.8 Hz, β-C$H_2$ b His), 2.68 (dd, 1H, $J$ = 13.8, 7.2 Hz, β-C$H_2$ b Phe), 2.02-1.95 (m, 1H, β-C$H$ Val), 1.94-1.89 (m, 1H, β-C$H_2$ a Pro), 1.89-1.82 (m, 1H, γ-C$H_2$ a Pro), 1.80-1.75 (m, 1H, γ-C$H_2$ b Pro), 1.75-1.70 (m, 1H, β-C$H_2$ b Pro), 1.68-1.61 (m, 1H, β-C$H$ Ile), 1.27-1.19 (m, 1H, γ-C$H_2$ a Ile), 0.93-0.81 (m, 1H, γ-C$H_2$ b Ile), 0.89 (d, 3H, $J$ = 6.8 Hz, γ-C$H_3$ a Val), 0.87 (d, 3H, $J$ = 6.8 Hz, γ-C$H_3$ b Val), 0.72 (t, 3H, $J$ = 7.4 Hz, δ-C$H_3$ Ile), 0.69 (d, 3H, $J$ = 7.2 Hz, γ-C$H_3$ Ile).

**$^{13}$C-NMR** (DMSO-$d_6$, 151 MHz): $\delta_C$ [ppm] = 171.0 ($C$O Pro), 170.82 ($C$O Ile), 170.75 ($C$O His), 169.9 ($C$O PA), 169.7 ($C$O Val), 138.9 (γ-$C_{quart}$ Phe), 136.2 ($C_{quart}$ PA), 134.5 (ε-$C$H$_{arom}$ His), 129.1, 129.0, 128.0, 126.2, 125.8 (δ-$C$H$_{arom}$ Phe, $C$H$_{arom}$ PA), 61.8 ($C$H$_2$OH Phe), 59.5 (α-$C$H Pro), 56.4 (α-$C$H Ile), 55.9 (α-$C$H Val), 52.9 (α-$C$H His), 52.2 (α-$C$H Phe), 47.0 (δ-$C$H$_2$N Pro), 42.0 ($C$H$_2$ PA), 36.7 (β-$C$H Ile), 36.3 (β-$C$H$_2$ Phe), 30.1 (weak, β-$C$H$_2$ His), 29.7 (β-$C$H Val), 29.1 (β-$C$H$_2$ Pro), 24.4 (γ-$C$H$_2$ Pro), 23.9 (γ-$C$H$_2$ Ile), 19.1 (γ-$C$H$_3$ a Val), 18.5 (γ-$C$H$_3$ b Val), 15.3 (γ-$C$H$_3$ Ile), 11.0 (δ-$C$H$_3$ Ile).

Additional found signals: $\delta_H$ [ppm] = 8.15 (FA), 3.17 (MeOH), 2.54 (DMSO). ε-N$H$ His was not observed. $\delta_C$ [ppm] = 40.4 (DMSO).

**UHR-MS (ESI-TOF)** *m/z* calcd for $C_{39}H_{54}N_7O_6$: 716.4130 $[M+H]^+$; found: 716.4121 $[M+H]^+$

**Specific rotation** $[\alpha]_D^{21}$ = -102.2° (c = 2.03, $CH_3OH$)

(2*R*/*S*)-1-((3-methylbutanoyl)-D-histidyl-L-valyl-L-valyl)-*N*-((*S*)-1-oxo-3-phenylpropan-2-yl)pyrrolidine-2-carboxamide (**15**)

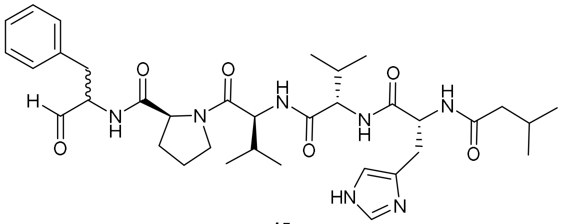

**15**

The alcohol **12** (57.9 mg, 0.0867 mmol) was suspended in anhydrous DCM and DMP (15% in DCM, 540 µL, 0.260 mmol) was added. The mixture was stirred at room temperature and the reaction progress was monitored by UHR-MS. After 2 h more DMP was added (90.0 µL, 0.130 mmol). The reaction was quenched with MeOH after 3 h and concentrated *in vacuo*. Purification of the crude product using semi preparative HPLC (5-35-95% AcCN + 0.1% FA, NUCLEODUR® C18 Gravity SB, 3 µm, 250 x 10 mm, flow rate: 2 mL/min) yielded **15** as a colorless solid (100% pure: 19.3 mg, 0.0289 mmol; 95% pure: 32.8 mg, 0.0469 mmol, calculated total yield: 87%).

**$^1$H-NMR** (DMSO-$d_6$, 600 MHz): $\delta_H$ [ppm] = 9.47 (s, 0.4H, C*H*O Phe), 9.41 (s, 0.2H, C*H*O Phe), 8.36 (d, 0.4H, *J* = 7.5 Hz, N*H* Phe), 8.34 (d, 0.2H, *J* = 7.2 Hz, N*H* Phe), 8.01-7.94 (m, 2H, N*H* Val 1, N*H* His), 7.68-7.61 (m, 1H, N*H* Val 2), 7.51 (s, 1H, ε-C$H_{arom}$ His), 7.30-7.10 (m, 5H, C$H_{arom}$ Phe), 6.77 (s, 1H, δ-C$H_{arom}$ His), 4.62-4.56 (m, 1H, α-C*H* His), 4.33-4.25 (m, 2 H, α-C*H* Phe, α-C*H* Pro, α-C*H* Val 1), 4.25-4.15 (m, 2 H, α-C*H* Phe, α-C*H* Val 2), 3.78-3.72 (m, 1H, δ-C$H_2$N a Pro), 3.57-3.47 (m, 1H, δ-C$H_2$N b Pro), 3.15-3.07 (m, 1H, β-C$H_2$ Phe), 2.93-2.82 (m, 2H, β-C$H_2$ a Phe, β-C$H_2$ a His), 2.78-2.69 (m, 2H, β-C$H_2$ b Phe, β-C$H_2$ b His), 2.04-1.84 (m, 7H, β-C$H_2$ Pro, γ-C$H_2$ Pro, β-C*H* Val 1, β-C*H* Val 2, $CH_2$ iVal, C*H* iVal), 1.84-1.71 (m, 2H, β-C$H_2$ Pro, γ-C$H_2$ Pro), 1.58-1.52 (m, 1H, β-C$H_2$ Pro, γ-C$H_2$ Pro), 0.89 (d, 3H, *J* = 6.8 Hz, γ-C$H_3$ a Val 1), 0.87 (d, 3H, *J* = 6.6 Hz, γ-C$H_3$ b Val 1), 0.82 (d, 3H, *J* = 6.4 Hz, C$H_3$ a iVal), 0.78 (d, 3H, *J* = 6.2 Hz, C$H_3$ b iVal), 0.75-0.68 (m, 6H, γ-C$H_3$ a+b Val 2).

**$^{13}$C-NMR** (DMSO-$d_6$, 151 MHz): $\delta_C$ [ppm] = 200.3 (*C*HO Phe), 172.1, 172.0 (*C*O Pro), 171.4 (*C*O iVal), 171.0 (*C*O His), 170.6 (*C*O Val 2), 169.62, 169.59 (*C*O Val 1), 137.62, 137.60 (γ-$C_{quart}$ Phe), 134.5 (weak, ε-*C*$H_{arom}$ His), 129.3, 129.2, 128.8, 128.2, 128.1, 127.7, 126.23, 126.18 (δ-*C*$H_{arom}$ Phe), 59.7, 59.5 (α-*C*H Phe), 59.1, 59.0 (α-*C*H Pro), 56.9 (α-*C*H Val 2), 55.8 (α-*C*H Val 1), 52.7 (α-*C*H His), 47.0 (δ-*C*$H_2$N Pro), 44.4 (*C*$H_2$ iVal), 33.5, 33.3 (β-*C*$H_2$ Phe), 30.65, 30.60 (β-*C*H Val 2), 29.8 (β-*C*$H_2$ His), 29.7 (β-*C*H Val 1), 29.3 (β-*C*$H_2$ Pro), 25.4 (*C*H iVal), 24.5, 24.4 (γ-*C*$H_2$ Pro), 22.2 (*C*$H_3$ a iVal), 22.1 (*C*$H_3$ b iVal), 19.0 (γ-*C*$H_3$ a Val 1, γ-*C*$H_3$ a Val 2), 18.44, 18.40 (γ-*C*$H_3$ b Val 1), 17.6 (γ-*C*$H_3$ b Val 2).

Additional found signals: $\delta_H$ [ppm] = 8.18 (FA), 3.17 (MeOH). ε-N*H* His was not observed. $\delta_C$ [ppm] = / , γ-$C_{quart}$ His, δ-*C*$H_{arom}$ His were not observed.

Due to racemization of the phenylalanine aldehyde, some chemical shifts appear as a double set.

**UHR-MS (ESI-TOF)** *m/z* calcd for $C_{35}H_{52}N_7O_6$: 666.3974 $[M+H]^+$; found: 666.3976 $[M+H]^+$

**Specific rotation** $[\alpha]_D^{21}$ = --37.5° (c = 0.80, $CH_3OH$)

(2*R*/*S*)-*N*-((*S*)-1-oxo-3-phenylpropan-2-yl)-1-((2-phenylacetyl)-D-histidyl-L-valyl-L-valyl)pyrrolidine-2-carboxamide (**16**)

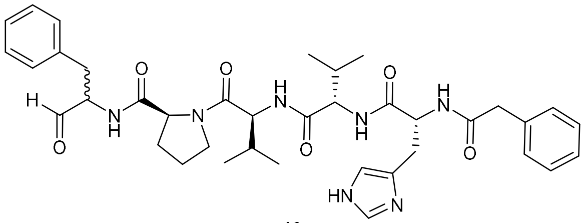

16

The alcohol **13** (14.8 mg, 0.0211 mmol) was suspended in anhydrous DCM and DMP (15% in DCM, 131 µL, 0.0632 mmol) was added. The mixture was stirred at room temperature and the reaction progress was monitored by UHR-MS. The reaction was quenched with MeOH after 2 h and concentrated *in vacuo*. Purification of the crude product using semi preparative HPLC (5-45-95% AcCN + 0.1% FA, NUCLEODUR® C18 Gravity SB, 3 µm, 250 x 10 mm, flow rate: 2 mL/min) yielded **16** as a colorless solid (12.7 mg, 0.0181 mmol, 86%).

**$^1$H-NMR** (DMSO-$d_6$, 600 MHz): $\delta_H$ [ppm] = 9.46 (s, 0.2H, C*H*O Phe), 9.41 (s, 0.3H, C*H*O Phe), 8.36 (d, 0.3H, *J* = 7.5 Hz, N*H* Phe), 8.34 (d, 0.3H, *J* = 6.8 Hz, N*H* Phe), 8.28 (d, 1H, *J* = 8.1 Hz, N*H* His), 7.95 (dd, 1H, *J* = 8.4, 8.4 Hz, N*H* Val 1), 7.76-7.71 (m, 1H, N*H* Val 2), 7.50 (s, 1H, ε-C$H_{arom}$ His), 7.30-7.12 (m, 10H, δ-C$H_{arom}$ Phe, C$H_{arom}$ PA), 6.76 (s, 1H, δ-C$H_{arom}$ His), 4.62-4.56 (m, 1H, α-C*H* His), 4.35-4.23 (m, 2 H, α-C*H* Pro, α-C*H* Val 1, α-C*H* Phe), 4.23-4.15 (m, 2 H, α-C*H* Phe, α-C*H* Val 2), 3.80-3.70 (m, 2H, δ-C$H_2$N a Pro), 3.59-3.47 (m, 4H, δ-C$H_2$N b Pro), 3.46 (d, 2H, *J* = 13.9 Hz, C$H_2$ a PA), 3.42 (d, 2H, *J* = 14.1 Hz, C$H_2$ b PA), 3.15-3.06 (m, 1H, β-C$H_2$ Phe), 2.94-2.82 (m, 2H, β-C$H_2$ His, β-C$H_2$ Phe), 2.79-2.71 (m, 2H, β-C$H_2$ His, β-C$H_2$ Phe), 2.04-1.84 (m, 4H, β-C*H* Val 1, β-C*H* Val 2, γ-C$H_2$ a Pro, β-C$H_2$ a Pro), 1.84-1.69 (m, 2H, γ-C$H_2$ b Pro, β-C$H_2$ b Pro), 1.58-1.50 (m, 1H, γ-C$H_2$ b Pro, β-C$H_2$ b Pro), 0.94-0.80 (m, 6H, γ-C$H_3$ Val 1), 0.75-0.63 (m, 6H, γ-C$H_3$ Val 2).

**$^{13}$C-NMR** (DMSO-$d_6$, 151 MHz): $\delta_C$ [ppm] = 200.3 (*C*HO Phe), 172.1, 172.0, 170.9, 170.7, 169.7, 169.6 (*C*$O_{carbonyl}$), 169.9 (*C*O PA), 137.6 (γ-$C_{quart}$ Phe), 136.2 ($C_{quart}$ PA), 134.5 (ε-*C*$H_{arom}$ His), 129.3, 129.2, 129.0, 128.6, 128.2, 128.1, 128.0, 126.25, 126.20, 126.17 (*C*$H_{arom}$ Phe, *C*$H_{arom}$ PA), 59.7, 59.5 (α-*C*H Phe), 59.1, 59.0 (α-*C*H Pro), 57.1 (α-*C*H Val 2), 55.8 (α-*C*H Val 1), 52.9 (α-*C*H His), 47.0 (δ-*C*$H_2$N Pro), 42.0 (*C*$H_2$ PA), 33.5, 33.3 (β-*C*$H_2$ Phe), 30.0* (β-*C*$H_2$ His), 30.6, 30.5 (β-*C*H Val 2), 29.7 (β-*C*H Val 1), 29.3 (β-*C*$H_2$ Pro), 24.5, 24.4 (γ-*C*$H_2$ Pro),19.0 (γ-*C*$H_3$ a Val 2), 18.49 (γ-*C*$H_3$ a Val 1), 18.46 (γ-*C*$H_3$ b Val 1), 17.6 (γ-*C*$H_3$ b Val 2).

Additional found signals: $\delta_H$ [ppm] = 8.19 (FA), Broad peak between 4.00 to 2.59 ($H_2O$), 3.17 (MeOH), ε-N*H* His was not observed. $\delta_C$ [ppm] = 163.4 (FA), γ-$C_{quart}$ His, δ-*C*$H_{arom}$ His were not observed.

Due to racemization of the phenylalanine aldehyde, some chemical shifts appear as a double set and some integrals aren't accurate due to the broad peak of $\delta_H$ = 4.00 to 2.59 ppm.

The carbonyl signals 172.1, 172.0, 170.9, 170.7, 169.7, 169.6 were not distinguishable using HMBC data.

**UHR-MS (ESI-TOF)** *m/z* calcd for $C_{38}H_{50}N_7O_6$: 700.3817 $[M+H]^+$; found: 700.3820 $[M+H]^+$

**Specific rotation** $[\alpha]_D^{21}$ = -66.7° (c = 0.75, $CH_3OH$)

(2*R*/S)-1-((3-methylbutanoyl)-D-histidyl-L-isoleucyl-L-valyl)-*N*-((*S*)-1-oxo-3-phenylpropan-2-yl)pyrrolidine-2-carboxamide (**2**)

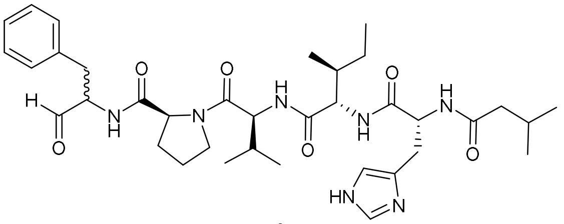

2

Alcohol **14** (19.7 mg, 0.0289 mmol) was suspended in anhydrous DCM and DMP (15% in DCM, 180 µL, 0.0867 mmol) was added. The mixture was stirred at room temperature and the reaction progress was monitored by UHR-MS. The reaction was quenched with MeOH after 2 h and concentrated *in vacuo*. Purification of the crude product using semi preparative HPLC (5-45-95% AcCN + 0.1% FA, NUCLEODUR® C18 Gravity SB, 3 µm, 250 x 10 mm, flow rate: 2 mL/min) yielded **2** as a colorless solid (12.1 mg, 0.0178 mmol, 62%).

**$^1$H-NMR** (DMSO-$d_6$, 600 MHz): $\delta_H$ [ppm] = 9.47 (s, 0.2H, C*H*O Phe), 9.41 (s, 0.2H, C*H*O Phe), 8.36 (d, 0.2H, *J* = 7.7 Hz, N*H* Phe), 8.34 (d, 0.2H, *J* = 7.0 Hz, N*H* Phe), 8.00-7.94 (m, 2H, N*H* Val, N*H* His), 7.69-7.63 (m, 1H, N*H* Ile), 7.50 (s, 1H, ε-C$H_{arom}$ His), 7.29-7.12 (m, 5H, δ-C$H_{arom}$ Phe), 6.76 (s, 1H, δ-C$H_{arom}$ His), 4.61-4.55 (m, 1H, α-C*H* His), 4.35-4.24 (m, 2H, α-C*H* Phe, α-C*H* Val, α-C*H* Pro), 4.24-4.16 (m, 2H, α-C*H* Phe, α-C*H* Ile), 3.78-3.68 (m, 1H, δ-C$H_2$N a Pro), 3.57-3.47 (m, 1H, δ-C$H_2$N b Pro), 3.15-3.07 (m, 1H, β-C$H_2$ Phe), 2.91-2.83 (m, 2H, β-C$H_2$ Phe, β-C$H_2$ a His), 2.78-2.69 (m, 2H, β-C$H_2$ Phe, β-C$H_2$ b His), 2.02-1.83 (m, 5H, β-C*H* Val, C$H_2$ iVal, C*H* iVal, β-C$H_2$ a Pro, γ-C$H_2$ a Pro), 1.83-1.70 (m, 2H, β-C$H_2$ b Pro, γ-C$H_2$ b Pro), 1.70-1.60 (m, 2H, β-C*H* Ile), 1.60-1.51 (m, 1H, β-C$H_2$ Pro), 1.30-1.22 (m, 1H, γ-C$H_2$ a Ile), 0.98-0.90 (m, 1H, γ-C$H_2$ b Ile), 0.90-0.84 (m, 6H, γ-C$H_3$ Val), 0.81 (d, 3H, *J* = 6.4 Hz, C$H_3$ a iVal), 0.78 (d, 3H, *J* = 6.4 Hz, C$H_3$ b iVal), 0.76-0.67 (m, 6H, δ-C$H_3$ Ile, γ-C$H_3$ Ile).

**$^{13}$C-NMR** (DMSO-$d_6$, 151 MHz): $\delta_C$ [ppm] = 200.3 (*C*HO Phe), 172.1, 172.0 (*C*O Pro), 171.4 (*C*O iVal), 170.9 (*C*O His), 170.7 (*C*O Ile), 169.58, 169.55 (*C*O Val), 137.6 (γ-$C_{quart}$ Phe), 134.5 (ε-$C$H$_{arom}$ His), 134.2 (γ-$C_{quart}$ His), 129.3, 129.2, 129.1, 128.9, 128.2, 128.1, 126.24, 126.19 (δ-$C$H$_{arom}$ Phe), 59.7, 59.5 (α-*C*H Phe), 59.1, 59.0 (α-*C*H Pro), 56.32, 56.29 (α-*C*H Ile), 55.7 (α-*C*H Val), 52.7 (α-*C*H His), 47.00, 46.96 (δ-*C*H$_2$N Pro), 44.5, (*C*H$_2$ iVal), 36.9, 36.8 (β-*C*H Ile), 33.6, 33.3 (β-*C*H$_2$ Phe), 29.9* (β-*C*H$_2$ His), 29.7 (β-*C*H Val),29.31, 29.28 (β-*C*H$_2$ Pro), 25.5 (*C*H iVal), 24.44, 24.38 (γ-*C*H$_2$ Pro), 24.0 (γ-*C*H$_2$ Ile), 22.2 (*C*H$_3$ a iVal), 22.1 (*C*H$_3$ b iVal), 19.03, 19.00 (γ-*C*H$_3$ a Val), 18.45, 18.41 (γ-*C*H$_3$ b Val), 15.2 (γ-*C*H$_3$ Ile), 11.0 (δ-*C*H$_3$ Ile).

Additional found signals: $\delta_H$ [ppm] = 8.19 (FA), 3.17 (MeOH), ε-N*H* His was not observed. $\delta_C$ [ppm] = δ-$C$H$_{arom}$ His was not observed.

Due to racemization of the phenylalanine aldehyde, some chemical shifts appear as a double set.

**UHR-MS (ESI-TOF)** *m/z* calcd for $C_{36}H_{54}N_7O_6$: 680.4130 [M+H]$^+$; found: 680.4128 [M+H]$^+$

**Specific rotation** $[\alpha]_D^{21}$ = -90.4° (c = 0.83, $CH_3OH$)

(2*R*/*S*)-*N*-((*S*)-1-oxo-3-phenylpropan-2-yl)-1-((2-phenylacetyl)-D-histidyl-L-isoleucyl-L-valyl)pyrrolidine-2-carboxamide (**3**)

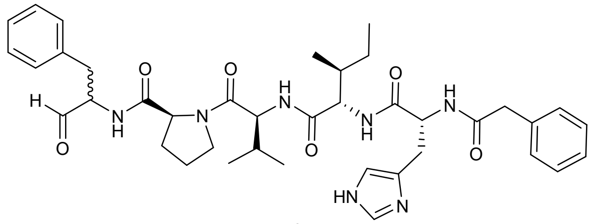

**3**

To a solution of alcohol **4** (36.9 mg, 0.0515 mmol) in anhydrous DCM DMP (15% in DCM, 321 µL, 0.155 mmol) was added. The mixture was stirred at room temperature and the reaction progress was monitored by UHR-MS. The reaction was quenched with MeOH after 2 h and concentrated *in vacuo*. Purification of the crude product using semi preparative HPLC (5-45-95% AcCN + 0.1% FA, NUCLEODUR® C18 Gravity SB, 3 µm, 250 x 10 mm, flow rate: 2 mL/min) yielded **3** as a colorless solid (24.5 mg, 0.0343 mmol, 67%).

**$^{1}$H-NMR** (DMSO-$d_6$, 600 MHz): $\delta_H$ [ppm] = 9.47 (s, 0.2H, C*H*O Phe), 9.41 (s, 0.2H, C*H*O Phe), 8.36 (d, 0.25H, *J* = 7.7 Hz, N*H* Phe), 8.34 (d, 0.25H, *J* = 7.7 Hz, N*H* Phe), 8.26 (d, 1H, *J* = 8.1 Hz, N*H* His), 7.95 (t, 1H, *J* = 9.0 Hz, N*H* Val), 7.77-7.71 (m, 1H, N*H* Ile), 7.51 (s, 1H, ε-C$H_{arom}$ His), 7.29-7.13 (m, 10H, δ-C$H_{arom}$ Phe, C$H_{arom}$ PA), 6.76 (s, 1H, δ-C$H_{arom}$ His), 4.62-4.55 (m, 1H, α-C*H* His), 4.33-4.24 (m, 2H, α-C*H* Val, α-C*H* Pro), 4.24-4.16 (m, 2H, α-C*H* Ile, α-C*H* Phe), 3.78-3.68 (m, 1H, δ-C$H_2$N a Pro), 3.57-3.47 (m, 1H, δ-C$H_2$N b Pro), 3.45 (d, 1H, *J* = 14.1 Hz, C$H_2$ a PA), 3.42 (d, 1H, *J* = 14.1 Hz, C$H_2$ b PA), 3.15-3.06 (m, 1H, β-C$H_2$ Phe), 2.93-2.83 (m, 2H, β-C$H_2$ His, β-C$H_2$ Phe), 2.78-2.71 (m, 1H, β-C$H_2$ His, β-C$H_2$ Phe), 2.02-1.84 (m, 2H, β-C*H* Val, β-C$H_2$ Pro, γ-C$H_2$ Pro), 1.84-1.69 (m, 2H, β-C$H_2$ Pro, γ-C$H_2$ Pro), 1.69-1.59 (m, 1H, β-C*H* Ile), 1.59-1.51 (m, 1H, β-C$H_2$ Pro), 1.27-1.17 (m, 1H, γ-C$H_2$ a Ile), 0.94-0.82 (m, 7H, γ-C$H_2$ b Ile, γ-C$H_3$ Val), 0.75-0.66 (m, 6H, δ-C$H_3$ Ile, γ-C$H_3$ Ile).

**$^{13}$C-NMR** (DMSO-$d_6$, 151 MHz): $\delta_C$ [ppm] = 200.3 (*C*HO Phe), 172.1, 172.0 (*C*O Pro), 170.8 (*C*O His), 170.7 (*C*O Ile), 169.9 (*C*O PA), 169.61, 169.58 (*C*O Val), 137.6 (γ-$C_{quart}$ Phe), 136.2 ($C_{quart}$ PA), 134.5 (ε-*C*$H_{arom}$ His), 129.3, 129.2, 129.14, 129.09, 129.0, 128.2, 128.1, 128.0, 126.25, 126.20, 126.16 (δ-*C*$H_{arom}$ Phe, *C*$H_{arom}$ PA), 59.5 (α-*C*H Pro), 59.1, 59.0 (α-*C*H Phe), 56.40, 56.37 (α-*C*H Ile), 55.8 (α-*C*H Val), 52.9 (α-*C*H His), 47.01, 46.97 (δ-*C*$H_2$N Pro), 42.0 (*C*$H_2$ PA), 36.8, 36.7 (β-*C*H Ile), 33.5, 33.3 (β-*C*$H_2$ Phe), 30.1* (β-*C*$H_2$ His), 29.7 (β-*C*H Val), 29.32, 29.29 (β-*C*$H_2$ Pro), 24.5, 24.4 (γ-*C*$H_2$ Pro), 23.9 (γ-*C*$H_2$ Ile), 19.03, 19.00 (γ-*C*$H_3$ a Val), 18.5, 18.4 (γ-*C*$H_3$ b Val), 15.3 (γ-*C*$H_3$ Ile),, 11.0 (δ-*C*$H_3$ Ile).

Additional found signals: $\delta_H$ [ppm] = 8.16 (FA), 3.17 (MeOH), ε-N*H* His was not observed. $\delta_C$ [ppm] = 59.7, 48.6 (MeOH), δ-*C*$H_{arom}$ His and γ-$C_{quart}$ His were not observed.

Due to racemization of the phenylalanine aldehyde, some chemical shifts appear as a double set.

**UHR-MS (ESI-TOF)** *m/z* calcd for $C_{39}H_{52}N_7O_6$: 714.3974 $[M+H]^+$; found: 714.3974 $[M+H]^+$

**Specific rotation** $[\alpha]_D^{21}$ = -74.6° (c = 0.67, $CH_3OH$)

### NMR spectra

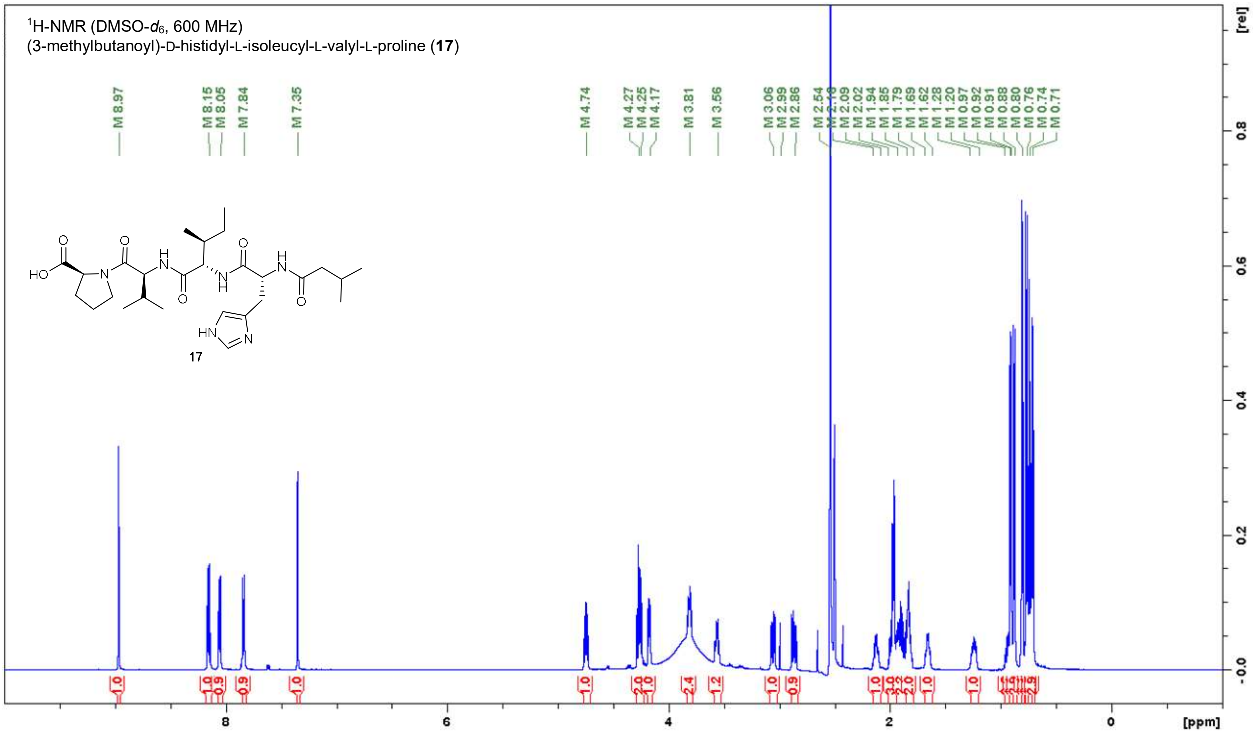

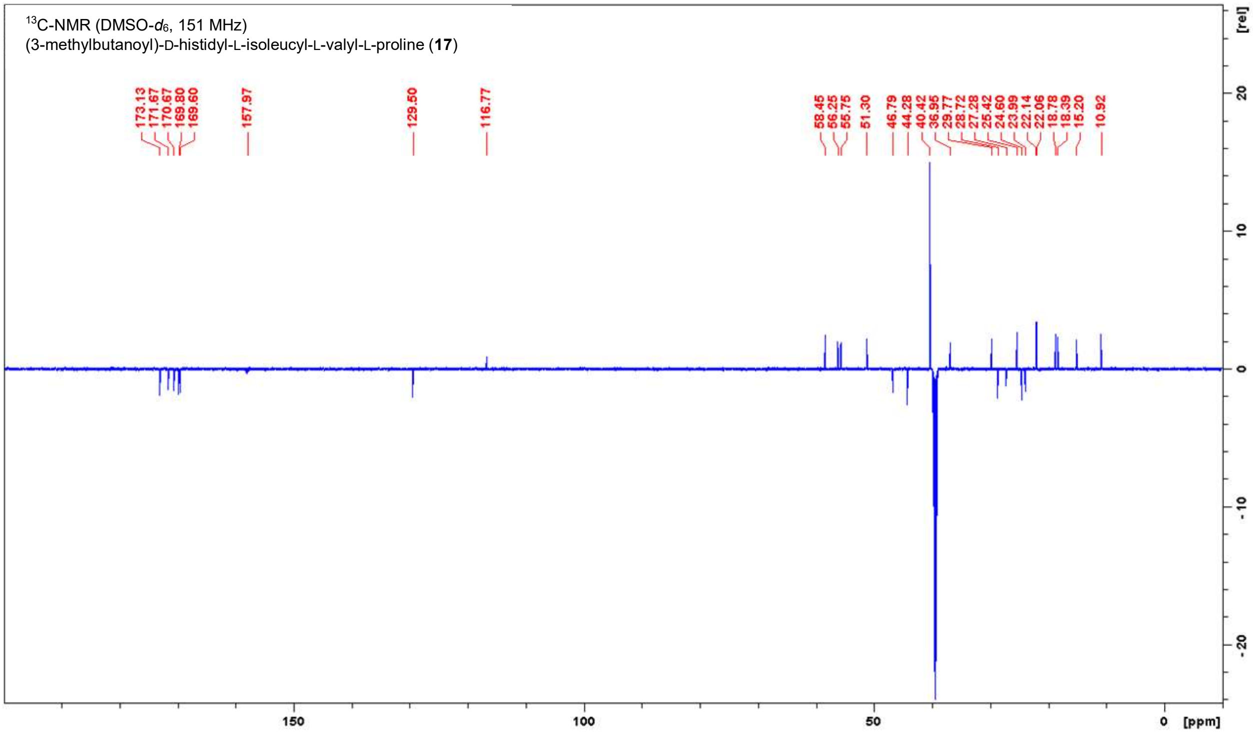

**Figure S7.** NMR spectra of **17** in DMSO-$d_6$.

**Table S1.** NMR data for **17** in DMSO-$d_6$ (600 MHz, 151 MHz).

| **Position** | **δ $^{13}$C [ppm]** | **δ $^{1}$H [ppm]; (m, ∫, *J*)** |
|---|---|---|
| L-Proline | | |
| **COOH** | 173.1 | n.o. |
| **α-CH** | 58.5 | 4.17 (dd, 1H, *J* = 8.7, 5.0 Hz) |
| **β-CH₂** | 28.7 | 2.16-2.09 (m, 1H)<br>1.85-1.79 (m, 2H, overlay) |
| **γ-CH₂** | 24.6 | 1.94-1.85 (m, 2H, overlay)<br>1.85-1.79 (m, 2H, overlay) |
| **δ-CH₂N** | 46.8 | 3.81 (ddd, 1H, *J* = 9.8, 6.4, 6.4 Hz)<br>3.56 (ddd, 1H, *J* = 10.1, 6.4, 6.4 Hz) |
| L-Valine | | |
| **CO** | 169.6 | - |
| **NH** | - | 8.05 (d, 1H, *J* = 8.3 Hz) |
| **α-CH** | 55.7 | 4.27 (dd, 1H, *J* = 9.0, 9.0 Hz) |
| **β-CH** | 29.8 | 2.02-1.94 (m, 3H, overlay) |
| **γ-CH₃** | 18.8 | 0.91 (d, 3H, *J* = 6.8 Hz) |
| **γ-CH₃** | 18.4 | 0.88 (d, 3H, *J* = 6.8 Hz) |
| L-Isoleucine | | |
| **CO** | 170.7 | - |
| **NH** | - | 7.84 (d, 1H, *J* = 8.8 Hz) |
| **α-CH** | 56.2 | 4.25 (dd, 1H, *J* =9.0, 7.0 Hz) |
| **β-CH** | 36.9 | 1.69-1.62 (m, 1H) |
| **γ-CH₂** | 24.0 | 1.28-1.20 (m, 1H)<br>0.97-0.92 (m, 1H) |
| **γ-CH₃** | 10.9 | 0.74 (t, 3H, *J* = 7.4 Hz) |
| **δ-CH₃** | 15.2 | 0.71 (d, 3H, *J* = 6.7 Hz) |
| D-Histidine | | |
| **CO** | 169.8 | - |
| **NH** | - | 8.15 (d, 1H, *J* = 8.4 Hz) |
| **α-CH** | 51.3 | 4.74 (ddd, 1H, *J* = 8.2, 8.2, 6.6 Hz) |
| **β-CH₂** | 27.3 | 3.06 (dd, 1H, *J* = 15.1, 6.1 Hz)<br>2.86 (dd, 1H, *J* = 15.1, 8.7 Hz) |
| **γ-C$_{quart}$** | 129.5 | - |
| **δ-CH$_{arom}$** | 116.8 | 7.35 (s, 1H) |
| **ε-CH$_{arom}$** | 133.7* | 8.97 (d, 1H, *J* = 1.1 Hz) |
| **ε-NH** | - | n.o. |
| Isovaleroyl | | |
| **CO** | 171.7 | - |
| **CH₂** | 44.3 | 2.02-1.94 (m, 3H, overlay) |
| **CH** | 25.4 | 1.94-1.85 (m, 2H, overlay) |
| **CH₃** | 22.14 | 0.80 (d, 3H, *J* = 6.8 Hz) |
| **CH₃** | 22.06 | 0.76 (d, 3H, *J* = 6.4 Hz) |

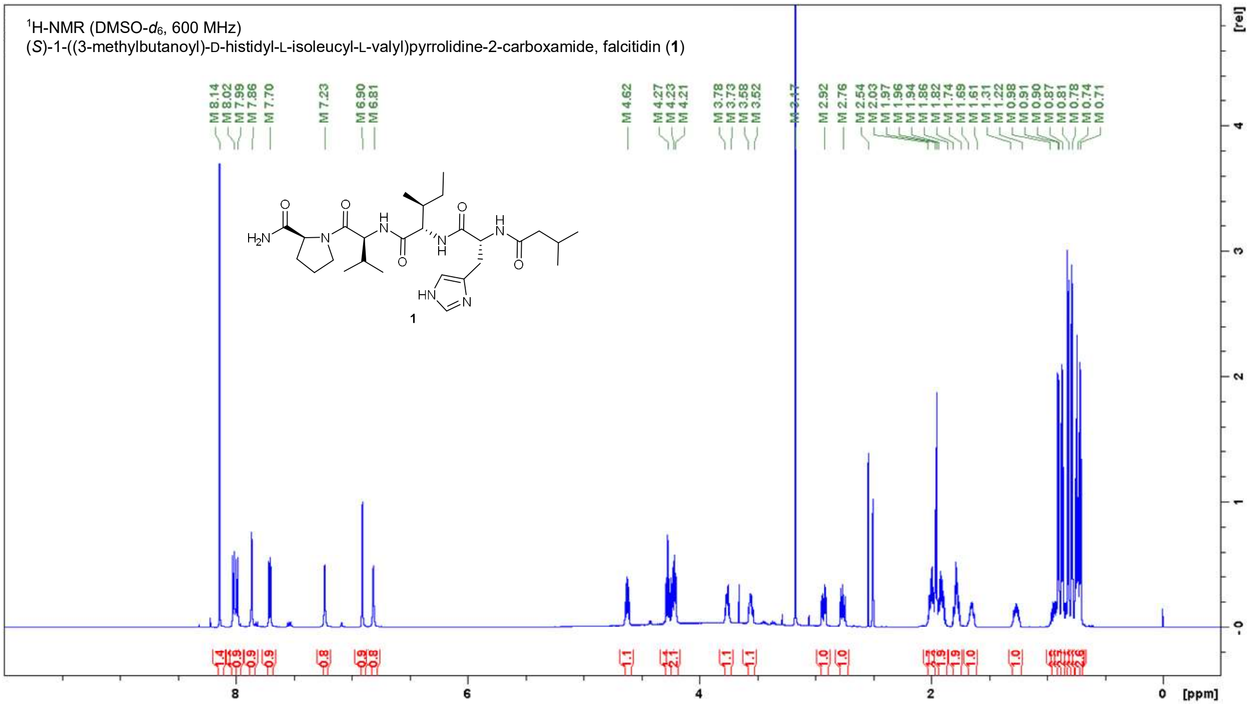

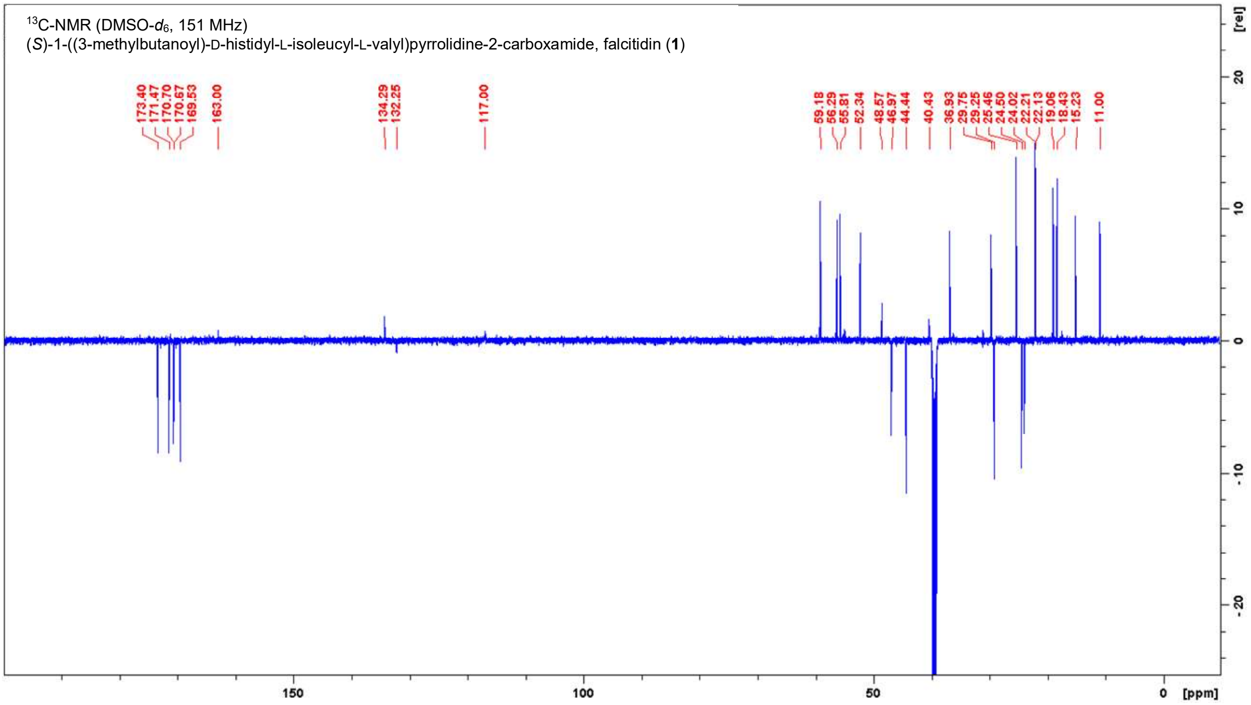

**Figure S8.** NMR spectra of **1** in DMSO-$d_6$.

**Table S2.** NMR data for **1** in DMSO-$d_6$ (600 MHz, 151 MHz).

| **Position** | **δ $^{13}$C [ppm]** | **δ $^{1}$H [ppm]; (m, ∫, *J*)** |
|---|---|---|
| L-Proline | | |
| **$CONH_2$** | 173.4 | 7.23 (s, 1H), 6.81 (s, 1H) |
| **α-CH** | 59.2 | 4.21 (dd, 1H, *J* = 8.5, 4.4 Hz) |
| **β-$CH_2$** | 29.3 | 2.03-1.97 (m, 2H, overlay)<br>1.82-1.74 (m, 2H, overlay) |
| **γ-$CH_2$** | 24.5 | 1.94-1.86 (m, 2H, overlay)<br>1.82-1.74 (m, 2H, overlay) |
| **δ-$CH_2$N** | 47.0 | 3.78-3.73 (m, 1H)<br>3.58-3.52 (m, 1H) |
| L-Valine | | |
| **CO** | 169.5 | - |
| **NH** | - | 7.99 (d, 1H, *J* = 8.3 Hz) |
| **α-CH** | 55.8 | 4.27 (dd, 1H, *J* =8.4, 8.4 Hz) |
| **β-CH** | 29.7 | 2.03-1.97 (m, 2H, overlay) |
| **γ-$CH_3$** | 19.1 | 0.90 (d, 1H, *J* =6.8 Hz) |
| **γ-$CH_3$** | 18.4 | 0.87 (d, 1H, *J* =6.7 Hz) |
| L-Isoleucine | | |
| **CO** | 170.70 | - |
| **NH** | - | 7.70 (d, 1H, *J* =8.8 Hz) |
| **α-CH** | 56.3 | 4.23 (dd, 1H, *J* =8.7, 6.9 Hz) |
| **β-CH** | 36.9 | 1.69-1.61 (m, 1H) |
| **γ-$CH_2$** | 24.0 | 1.31-1.22 (m, 1H)<br>0.98-0.91 (m, 1H) |
| **γ-$CH_3$** | 11.0 | 0.71 (d, 1H, *J* =7.0 Hz) |
| **δ-$CH_3$** | 15.2 | 0.74 (t, 1H, *J* =7.4 Hz) |
| D-Histidine | | |
| **CO** | 170.67 | - |
| **NH** | - | 8.02 (d, *J* = 8.2 Hz) |
| **α-CH** | 52.3 | 4.62 (ddd, *J* =8.3, 8.3, 6.1 Hz) |
| **β-$CH_2$** | 29.3 | 2.92 (dd, *J* =14.9, 5.9 Hz)<br>2.76 (dd, *J* =15.1, 8.8 Hz) |
| **γ-$C_{quart}$** | 132.3, weak | - |
| **δ-$CH_{arom}$** | 117.0, weak | 6.90 (s, 1H) |
| **ε-$CH_{arom}$** | 134.3, weak | 7.86 (s, 1H) |
| **ε-NH** | - | n.o. |
| Isovaleroyl | | |
| **CO** | 171.5 | - |
| **$CH_2$** | 44.4 | 1.96 (d, *J* =6.7 Hz) |
| **CH** | 25.5 | 1.94-1.86 (m, 2H, overlay) |
| **$CH_3$** | 22.2 | 0.81 (d, *J* =6.5 Hz) |
| **$CH_3$** | 22.1 | 0.78 (d, *J* =6.5 Hz) |

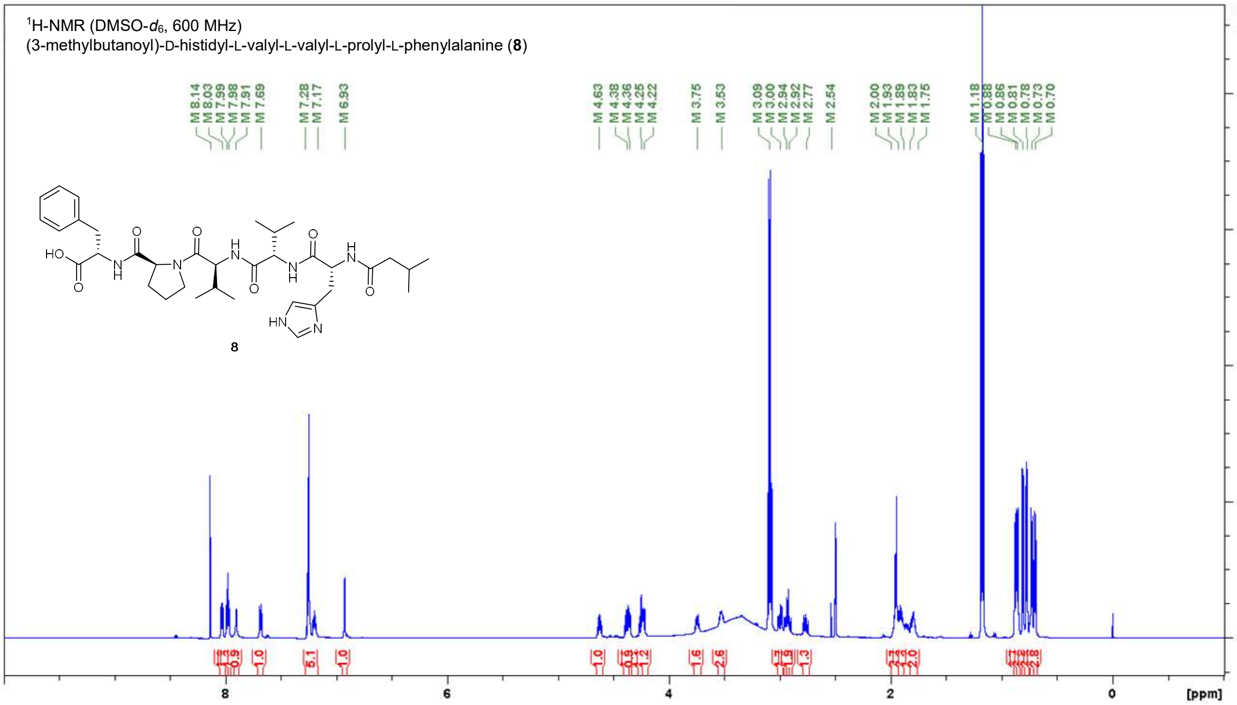

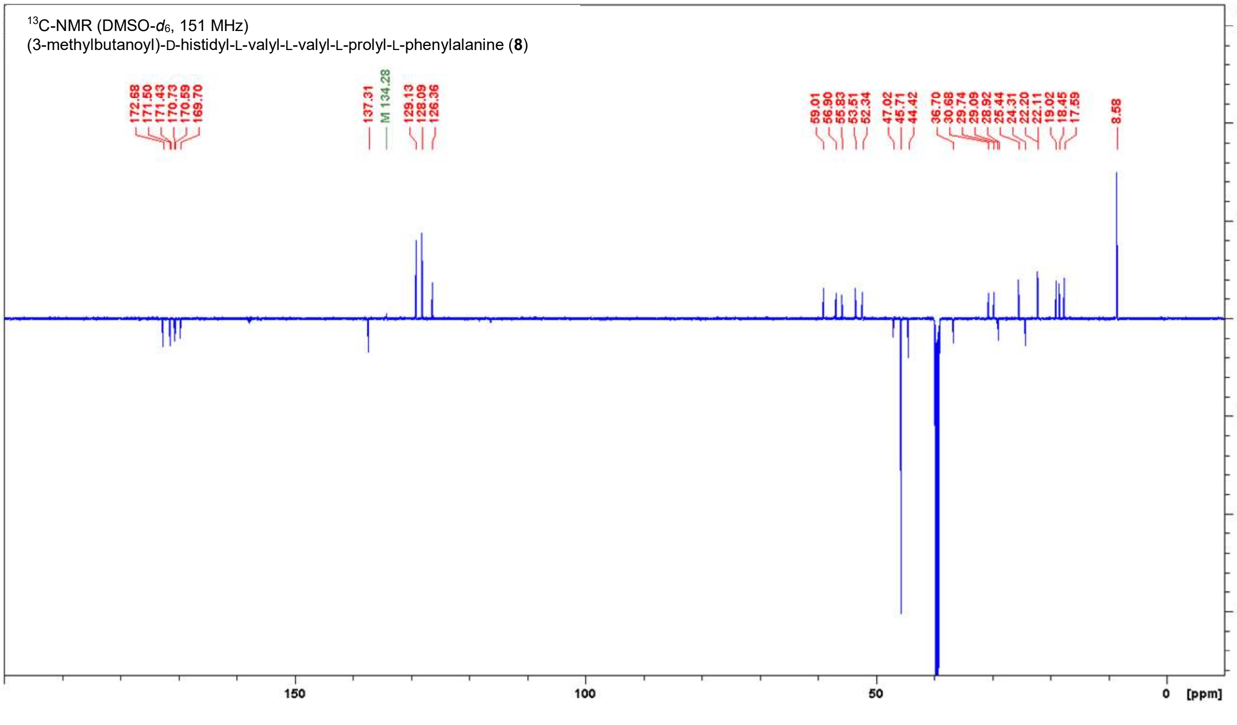

**Figure S9.** NMR spectra of **8** in DMSO-$d_6$.

**Table S3.** NMR data for **8** in DMSO-$d_6$ (600 MHz, 151 MHz).

| **Position** | **δ $^{13}$C [ppm]** | **δ $^{1}$H [ppm]; (m, ∫, *J*)** |
|---|---|---|
| L-Phenylalanine | | |
| **COOH** | 172.7 | n.o. |
| **NH** | - | 7.98 (d, 1H, *J* = 7.2 Hz) |
| **α-CH** | 53.5 | 4.38 (ddd, 1H, *J* = 7.6, 7.6, 5.7 Hz) |
| **β-CH$_2$ a** | 36.7 | 3.00 (dd, 1H, *J* = 14.2, 5.8 Hz)<br>2.92 (dd, 1H, *J* = 13.8, 7.5 Hz) |
| **γ-C$_{quart}$** | 137.3 | - |
| **CH$_{arom}$** | 129.1, 128.1, 126.4 | 7.28-7.17 (m, 5H) |
| L-Proline | | |
| **CO** | 171.4 | - |
| **α-CH** | 59.0 | 4.36 (dd, 1H, *J* = 8.3, 3.7 Hz) |
| **β-CH$_2$** | 28.9 | 2.00-1.93 (m, 4H, overlay)<br>1.83-1.75 (m, 2H, overlay) |
| **γ-CH$_2$** | 24.3 | 1.89-1.83 (m, 1H)<br>1.83-1.75 (m, 2H, overlay) |
| **δ-CH$_2$N** | 47.0 | 3.75 (ddd, 1H, *J* = 9.6, 6.9, 6.9 Hz)<br>3.53 (ddd, 1H, *J* = 9.4, 7.1, 5.6 Hz) |
| L-Valine 1 | | |
| **CO** | 169.7 | - |
| **NH** | - | 7.99 (d, 1H, *J* = 7.9 Hz) |
| **α-CH** | 55.8 | 4.25 (dd, 1H, *J* = 8.3, 8.3 Hz) |
| **β-CH** | 29.7 | 2.00-1.93 (m, 4H, overlay) |
| **γ-CH$_3$** | 19.0 | 0.88 (d, 3H, *J* = 6.8 Hz) |
| **γ-CH$_3$** | 18.4 | 0.86 (d, 3H, *J* = 6.4 Hz) |
| L-Valine 2 | | |
| **CO** | 170.6 | - |
| **NH** | - | 7.69 (d, 1H, *J* = 9.0 Hz) |
| **α-CH** | 56.9 | 4.22 (dd, 1H, *J* = 8.9, 6.1 Hz |
| **β-CH** | 30.7 | 1.93-1.89 (m, 2H, overlay) |
| **γ-CH$_3$** | 19.0 | 0.73 (d, 3H, *J* = 6.6 Hz) |
| **γ-CH$_3$** | 17.6 | 0.70 (d, 3H, *J* = 7.0 Hz) |
| D-Histidine | | |
| **CO** | 170.7 | - |
| **NH** | - | 8.03 (d, 1H, *J* = 8.1 Hz) |
| **α-CH** | 52.3 | 4.63 (ddd, 1H, *J* = 8.5, 8.5, 6.0 Hz) |
| **β-CH$_2$ a** | 29.1, weak | 2.94 (dd, 1H, *J* = 14.6, 6.5 Hz)<br>2.77 (dd, 1H, *J* = 13.8, 7.5 Hz) |
| **γ-C$_{quart}$** | 132.2* | - |
| **δ-CH$_{arom}$** | 117.0* | 6.93 (s, 1H) |
| **ε-CH$_{arom}$** | 134.3, weak | 7.91 (bs, 1H) |
| **ε-NH** | - | n.o. |
| Isovaleroyl | | |
| **CO** | 171.5 | - |
| **CH$_2$** | 44.4 | 2.00-1.93 (m, 4H, overlay) |
| **CH** | 25.4 | 1.93-1.89 (m, 2H, overlay) |
| **CH$_3$** | 22.2 | 0.81 (d, 3H, *J* = 6.4 Hz) |
| **CH$_3$** | 22.1 | 0.78 (d, 3H, *J* = 6.6 Hz) |

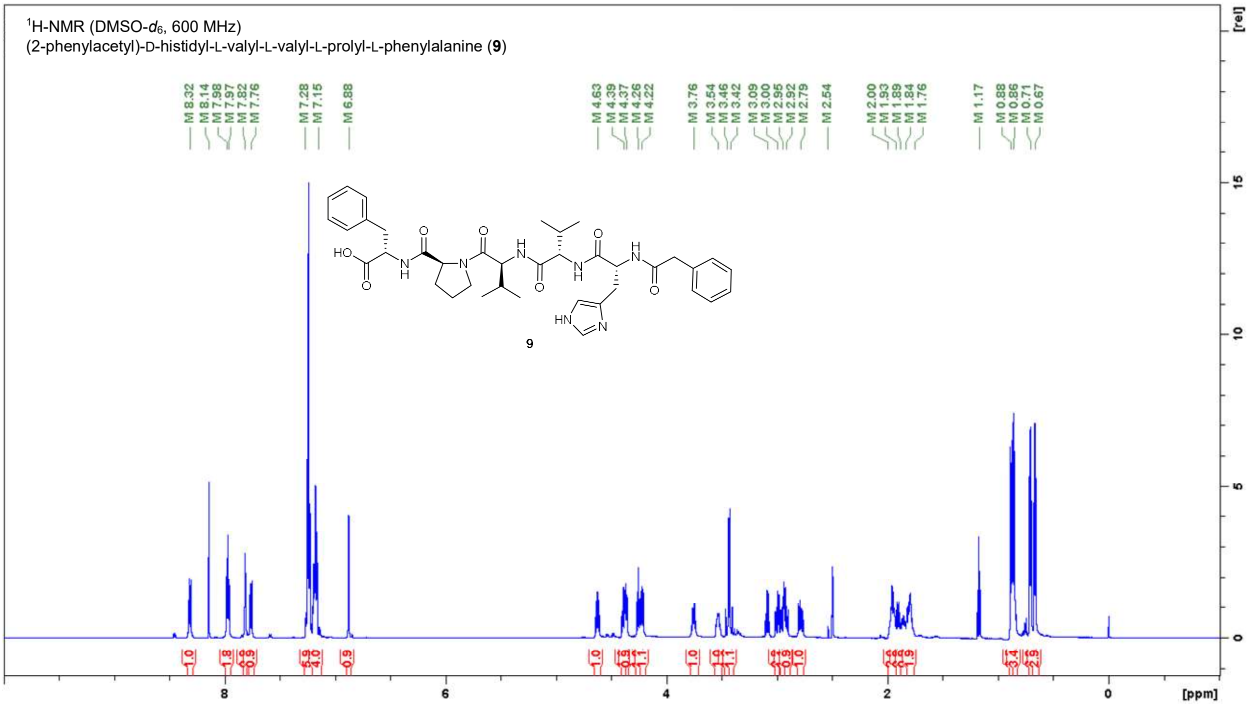

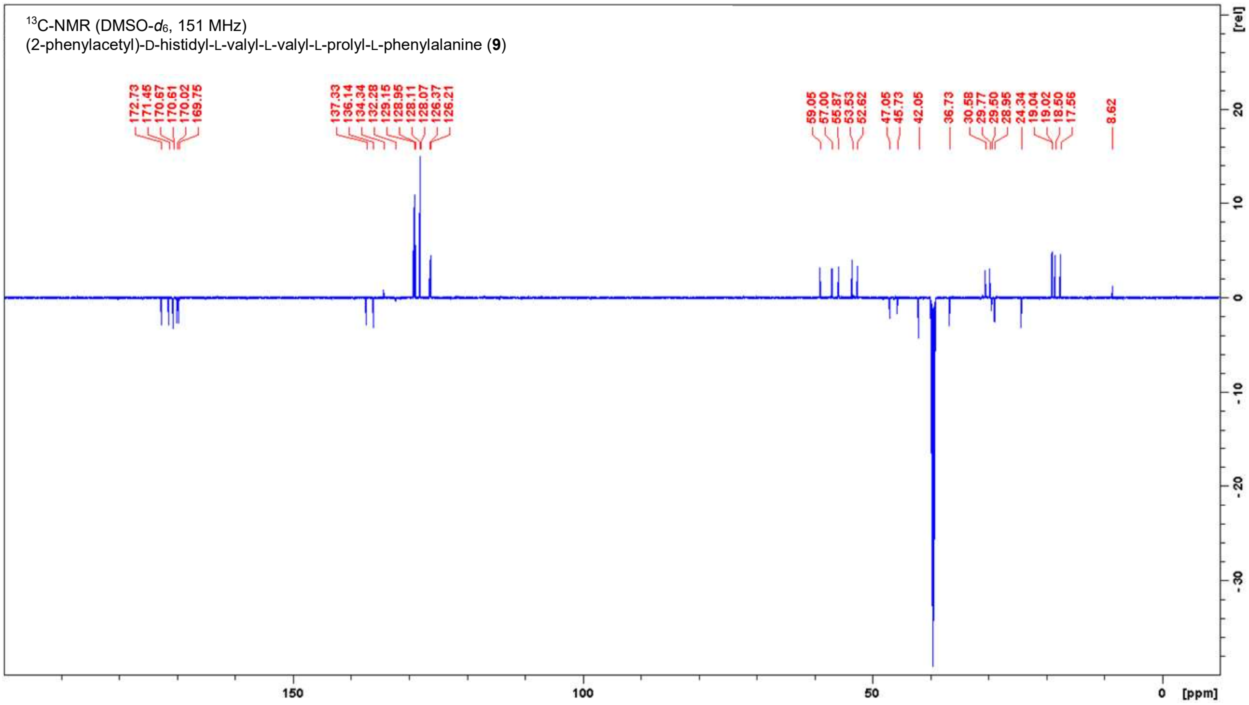

**Figure S10.** NMR spectra of **9** in DMSO-$d_6$.

**Table S4.** NMR data for **9** in DMSO-$d_6$ (600 MHz, 151 MHz).

| Position | δ $^{13}$C [ppm] | δ $^{1}$H [ppm]; (m, ∫, $J$) |
|---|---|---|
| L-Phenylalanine | | |
| **COOH** | 172.7 | n.o. |
| **NH** | - | 7.98 (d, 1H, $J$ = 7.2 Hz) |
| **α-CH** | 53.5 | 4.39 (ddd, 1H, $J$ = 7.6, 7.6, 6.0 Hz |
| **β-$CH_2$ a** | 36.7 | 3.00 (dd, 1H, $J$ = 14.1, 5.9 Hz)<br>2.92 (dd, 1H, $J$ = 13.9, 7.7 Hz) |
| **γ-$C_{quart}$** | 137.3 | - |
| **$CH_{arom}$** | 129.1, 128.9, 128.1, 128.1, 126.4, 126.2 | 7.28-7.15 (m, 10H, overlay) |
| L-Proline | | |
| **CO** | 171.5 | - |
| **α-CH** | 59.0 | 4.37 (dd, 1H, $J$ = 8.5, 3.4 Hz) |
| **β-$CH_2$** | 28.9 | 2.00-1.93 (m, 2H, overlay)<br>1.84-1.76 (m, 2H, overlay) |
| **γ-$CH_2$** | 24.3 | 1.89-1.84 (m, 1H)<br>1.84-1.76 (m, 2H, overlay) |
| **δ-$CH_2$N** | 47.0 | 3.76 (ddd, 1H, $J$ = 9.8, 6.8, 6.8 Hz)<br>3.54 (ddd, 1H, $J$ = 9.4, 7.2, 5.4 Hz) |
| L-Valine 1 | | |
| **CO** | 169.7 | - |
| **NH** | - | 7.97 (d, 1H, $J$ = 7.5 Hz) |
| **α-CH** | 55.9 | 4.26 (dd, 1H, $J$ = 8.3, 8.3 Hz) |
| **β-CH** | 29.8 | 2.00-1.93 (m, 2H, overlay) |
| **γ-$CH_3$** | 19.0 | 0.88 (d, 3H, $J$ = 6.6 Hz) |
| **γ-$CH_3$** | 18.5 | 0.86 (d, 3H, $J$ = 6.7 Hz) |
| L-Valine 2 | | |
| **CO** | 170.6 | - |
| **NH** | - | 7.76 (d, 1H, $J$ = 8.9 Hz) |
| **α-CH** | 57.0 | 4.22 (dd, 1H, $J$ = 9.0, 6.2 Hz) |
| **β-CH** | 30.6 | 1.93-1.89 (m, 1H) |
| **γ-$CH_3$** | 19.0 | 0.71 (d, 3H, $J$ = 6.8 Hz) |
| **γ-$CH_3$** | 17.6 | 0.67 (d, 3H, $J$ = 6.7 Hz) |
| D-Histidine | | |
| **CO** | 170.7 | - |
| **NH** | - | 8.32 (d, 1H, $J$ = 8.3 Hz) |
| **α-CH** | 52.6 | 4.63 (ddd, 1H, $J$ = 7.7, 7.7, 6.6 Hz) |
| **β-$CH_2$** | 29.5 | 2.79 (dd, 1H, $J$ = 14.8, 8.4 Hz) |
| **γ-$C_{quart}$** | 132.3, weak | - |
| **δ-$CH_{arom}$** | 117.0* | 6.88 (s, 1H) |
| **ε-$CH_{arom}$** | 134.3 | 7.82 (s, 1H) |
| **ε-NH** | - | n.o. |
| Phenylacetyl | | |
| **CO** | 170.0 | - |
| **$CH_2$** | 42.0 | 3.46 (d 1H, $J$ = 14.3 Hz)<br>3.42 (d 1H, $J$ = 14.2 Hz) |
| **$C_{quart}$** | 136.1 | |
| **$CH_{arom}$** | 129.1, 128.9, 128.1, 128.1, 126.4, 126.2 | 7.28-7.15 (m, 10H, overlay) |

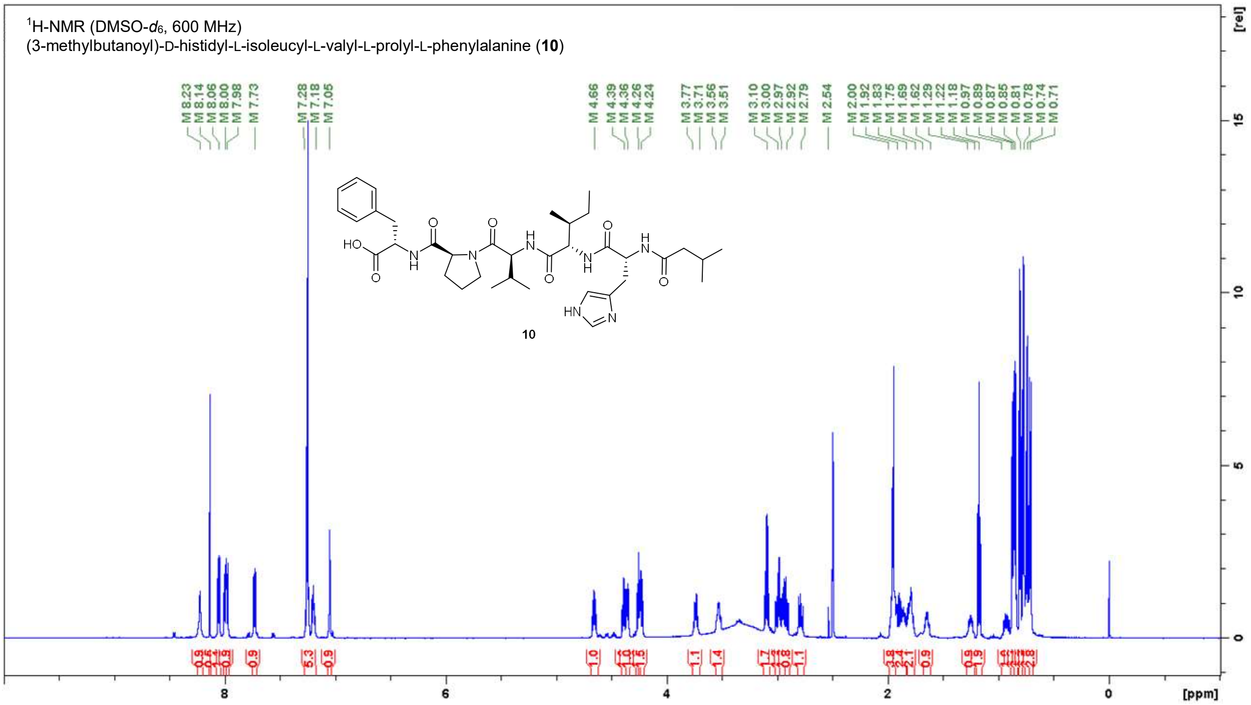

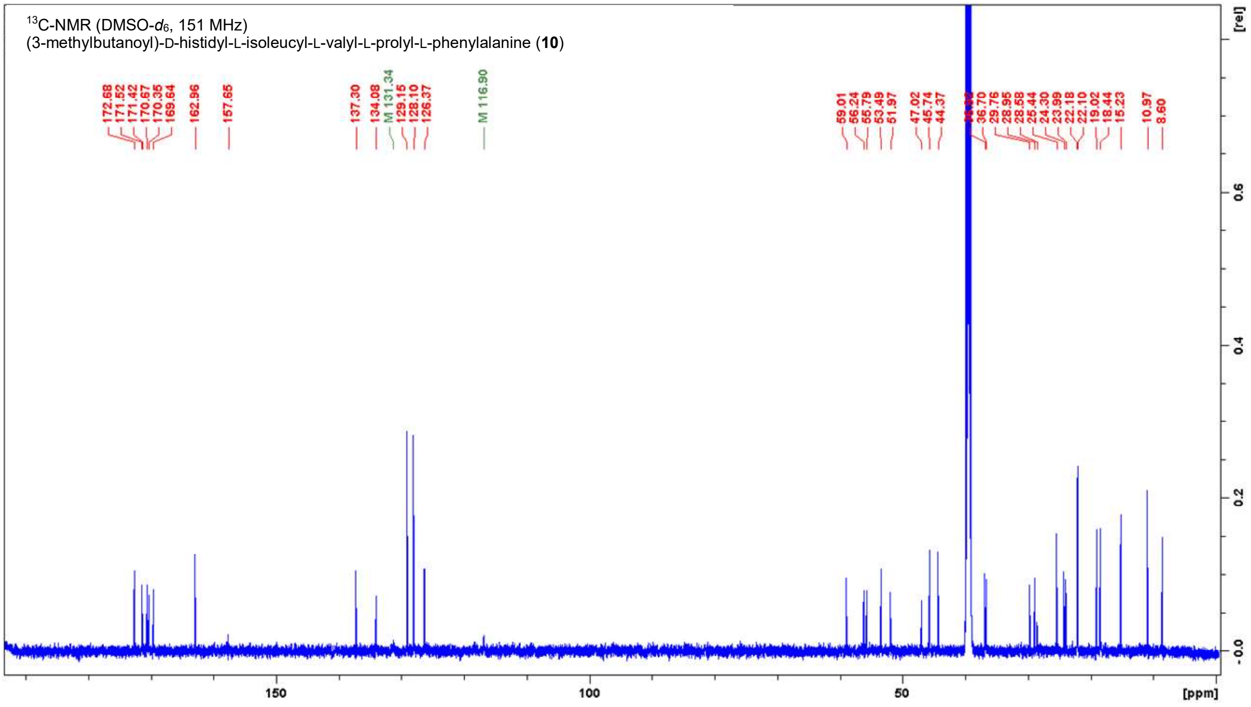

**Figure S11.** NMR spectra of **10** in DMSO-$d_6$.

**Table S5.** NMR data for **10** in DMSO-$d_6$ (600 MHz, 151 MHz).

| Position | δ $^{13}$C [ppm] | δ $^{1}$H [ppm]; (m, ∫, *J*) |
|---|---|---|
| L-Phenylalanine | | |
| **COOH** | 172.7 | n.o. |
| **NH** | - | 7.98 (d, 1H, *J* = 7.7 Hz) |
| **α-CH** | 53.5 | 4.39 (ddd, 1H, *J* = 7.3, 7.3, 6.2 Hz) |
| **β-CH$_2$** | 36.7 | 3.00 (dd, 1H, *J* = 13.9, 5.9 Hz)<br>2.92 (dd, 1H, *J* = 13.9, 7.9 Hz) |
| **γ-C$_{quart}$** | 137.3 | - |
| **CH$_{arom}$** | 129.1, 128.1, 126.4 | 7.28-7.18 (m, 5H) |
| L-Proline | | |
| **CO** | 171.4 | - |
| **α-CH** | 59.0 | 4.36 (dd, 1H, *J* = 8.3, 3.9 Hz) |
| **β-CH$_2$** | 28.9 | 2.00-1.92 (m, 4H, overlay)<br>1.83-1.75 (m, 2H, overlay) |
| **γ-CH$_2$** | 24.3 | 1.92-1.83 (m, 2H, overlay)<br>1.83-1.75 (m, 2H, overlay) |
| **δ-CH$_2$N** | 47.0 | 3.77-3.71 (m, 1H)<br>3.56-3.51 (m, 1H) |
| L-Valine | | |
| **CO** | 169.6 | - |
| **NH** | - | 8.00 (d, 1H, *J* = 8.4 Hz) |
| **α-CH** | 55.8 | 4.26 (dd, 1H, *J* = 8.3, 8.3 Hz) |
| **β-CH** | 29.8 | 2.00-1.92 (m, 4H, overlay) |
| **γ-CH$_3$** | 19.0 | 0.87 (d, 3H, *J* = 6.8 Hz) |
| **γ-CH$_3$** | 18.4 | 0.85 (d, 3H, *J* = 6.4 Hz) |
| L-Isoleucine | | |
| **CO** | 170.7 | - |
| **NH** | - | 7.73 (d, 1H, *J* = 9.2 Hz) |
| **α-CH** | 56.2 | 4.24 (dd, 1H, *J* = 8.5, 7.5 Hz) |
| **β-CH** | 36.9 | 1.69-1.62 (m, 1H) |
| **γ-CH$_2$** | 24.0 | 1.29-1.22 (m, 1H)<br>0.97-0.89 (m, 1H) |
| **γ-CH$_3$** | 15.2 | 0.71 (d, 3H, *J* = 7.0 Hz) |
| **δ-CH$_3$** | 11.0 | 0.74 (t, 3H, *J* = 7.4 Hz) |
| D-Histidine | | |
| **CO** | 170.4 | - |
| **NH** | - | 8.06 (d, 1H, *J* = 8.1 Hz) |
| **α-CH** | 52.0 | 4.66 (ddd, 1H, *J* = 8.1, 8.1, 6.5 Hz) |
| **β-CH$_2$** | 28.6 | 2.97 (dd, 1H, *J* = 15.3, 6.3 Hz)<br>2.79 (dd, 1H, *J* = 15.0, 8.6 Hz) |
| **γ-C$_{quart}$** | 131.3 | - |
| **δ-CH$_{arom}$** | 116.9 | 7.05 (s, 1H) |
| **ε-CH$_{arom}$** | 134.1 | 8.23 (s, 1H) |
| **ε-NH His** | - | n.o. |
| Isovaleroyl | | |
| **CO** | 171.5 | - |
| **CH$_2$** | 44.4 | 2.00-1.92 (m, 4H, overlay) |
| **CH** | 25.4 | 1.92-1.83 (m, 2H, overlay) |
| **CH$_3$** | 22.2 | 0.81 (d, 3H, *J* = 6.6 Hz) |
| **CH$_3$** | 22.1 | 0.78 (d, 3H, *J* = 6.6 Hz) |

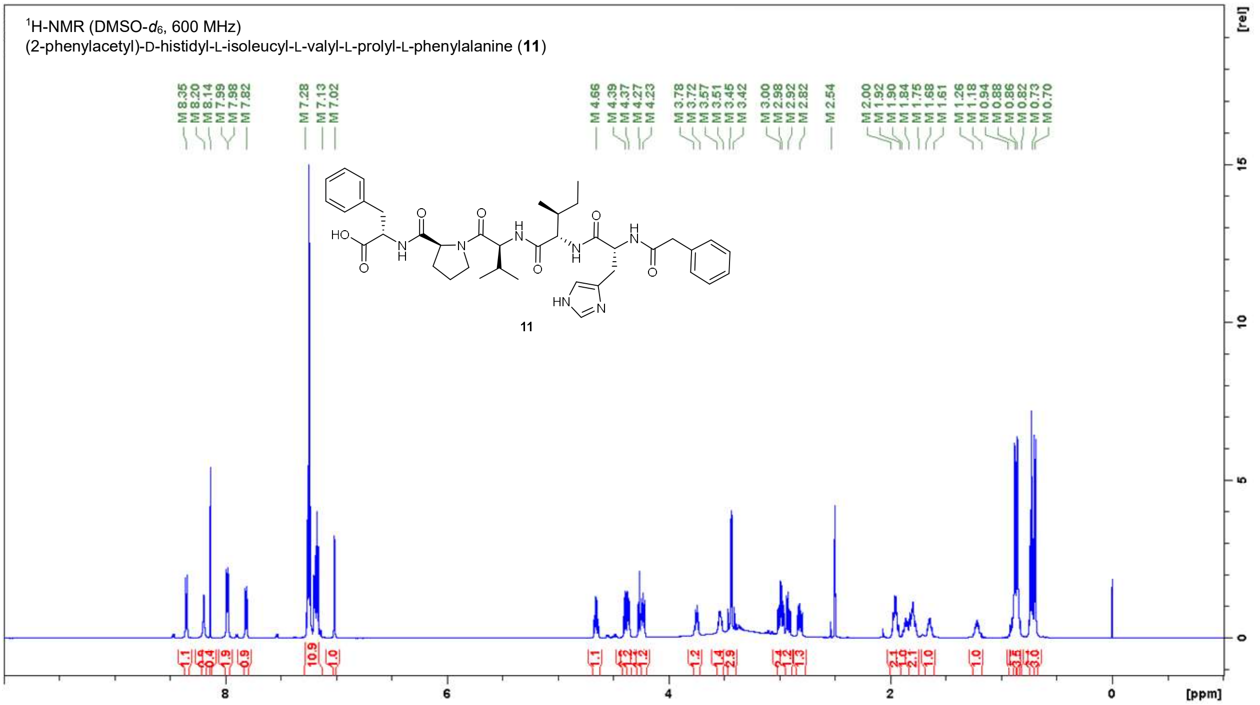

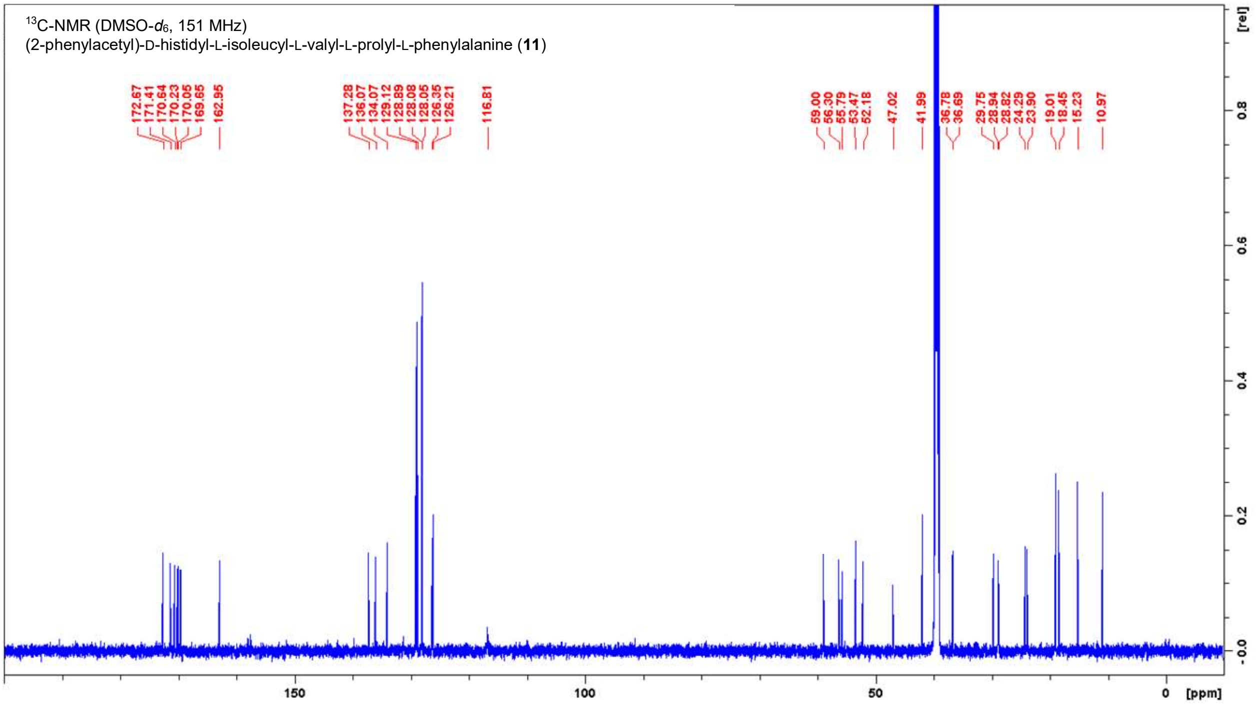

**Figure S12.** NMR spectra of **11** in DMSO-$d_6$.

**Table S6.** NMR data for **11** in DMSO-$d_6$ (600 MHz, 151 MHz).

| **Position** | **δ $^{13}$C [ppm]** | **δ $^{1}$H [ppm]; (m, ∫, *J*)** |
|---|---|---|
| L-Phenylalanine | | |
| **COOH** | 172.7 | n.o. |
| **NH** | | 7.99 (d, 1H, *J* = 7.5 Hz) |
| **α-CH** | 53.5 | 4.39 (ddd, 1H, *J* = 7.8, 7.8, 5.9 Hz) |
| **β-$CH_2$** | 36.7 | 2.92 (dd, 1H, *J* = 13.9, 7.9 Hz) |
| **γ-$C_{quart}$** | 137.3 | - |
| **$CH_{arom}$** | 129.1, 128.9, 128.1,<br>128.0, 126.4, 126.2 | 7.28-7.13 (m, 10H, overlay) |
| L-Proline | | |
| **CO** | 171.4 | - |
| **α-CH** | 59.0 | 4.37 (dd, 1H, *J* = 8.7, 4.1 Hz) |
| **β-$CH_2$** | 28.9 | 2.00-1.92 (m, 2H, overlay)<br>1.84-1.75 (m, 2H, overlay) |
| **γ-$CH_2$** | 24.3 | 1.90-1.84 (m, 1H)<br>1.84-1.75 (m, 2H, overlay) |
| **δ-$CH_2$N** | 47.0 | 3.78-3.72 (m, 1H)<br>3.57-3.51 (m, 1H) |
| L-Valine | | |
| **CO** | 169.7 | - |
| **NH** | - | 7.98 (d, 1H, *J* = 8.0 Hz) |
| **α-CH** | 55.8 | 4.27 (dd, 1H, *J* = 8.3, 8.3 Hz) |
| **β-CH** | 29.7 | 2.00-1.92 (m, 2H, overlay) |
| **γ-$CH_3$** | 19.0 | 0.88 (d, 3H, *J* = 6.4 Hz) |
| **γ-$CH_3$** | 18.5 | 0.86 (d, 3H, *J* = 6.6 Hz) |
| L-Isoleucine | | |
| **CO** | 170.6 | - |
| **NH** | - | 7.82 (d, 1H, *J* = 8.6 Hz) |
| **α-CH** | 56.3 | 4.23 (dd, 1H, *J* = 8.8, 7.0 Hz) |
| **β-CH** | 36.8 | 1.68-1.61 (m, 1H) |
| **γ-$CH_2$** | 23.9 | 1.26-1.18 (m, 1H)<br>0.94-0.82 (m, 1H) |
| **γ-$CH_3$** | 15.2 | 0.70 (d, 3H, *J* = 7.0 Hz) |
| **δ-$CH_3$** | 11.0 | 0.73 (t, 3H, *J* = 7.4 Hz) |
| D-Histidine | | |
| **CO** | 170.2 | - |
| **NH** | - | 8.35 (d, 1H, *J* = 8.1 Hz) |
| **α-CH** | 52.2 | 4.66 (ddd, 1H, *J* = 8.2, 8.2, 6.4 Hz) |
| **β-$CH_2$** | 28.8 | 2.98 (dd, 1H, *J* = 14.5, 5.9 Hz)<br>2.82 (dd, 1H, *J* = 14.9, 8.3 Hz) |
| **γ-$C_{quart}$** | 131.3* | - |
| **δ-$CH_{arom}$** | 116.8 | 7.02 (s, 1H) |
| **ε-$CH_{arom}$** | 134.1 | 8.20 (s, 1H) |
| **ε-NH** | | n.o. |
| Phenylacetyl | | |
| **CO** | 170.1 | |
| **$CH_2$ a** | 42.0 | 3.45 (d, 1H, *J* = 14.2 Hz)<br>3.42 (d, 1H, *J* = 14.2 Hz) |
| **$C_{quart}$** | 136.1 | |
| **$CH_{arom}$** | 129.1, 128.9, 128.1,<br>128.0, 126.4, 126.2 | 7.28-7.13 (m, 10H, overlay) |

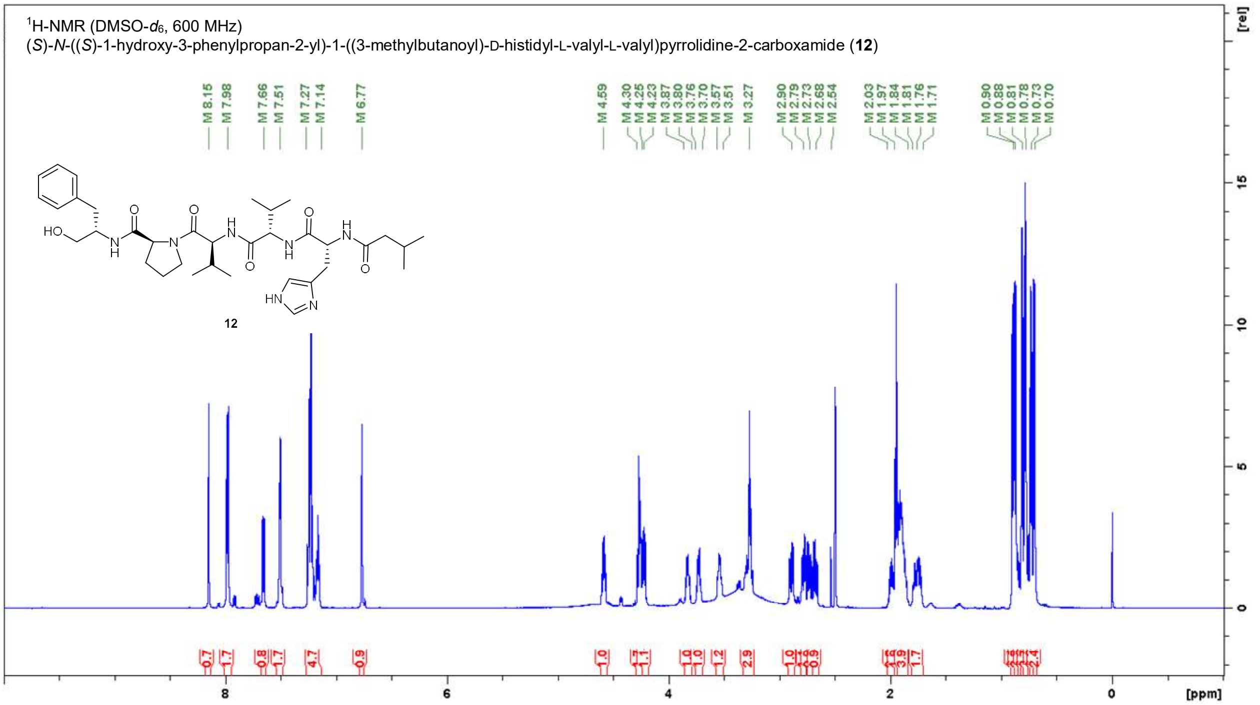

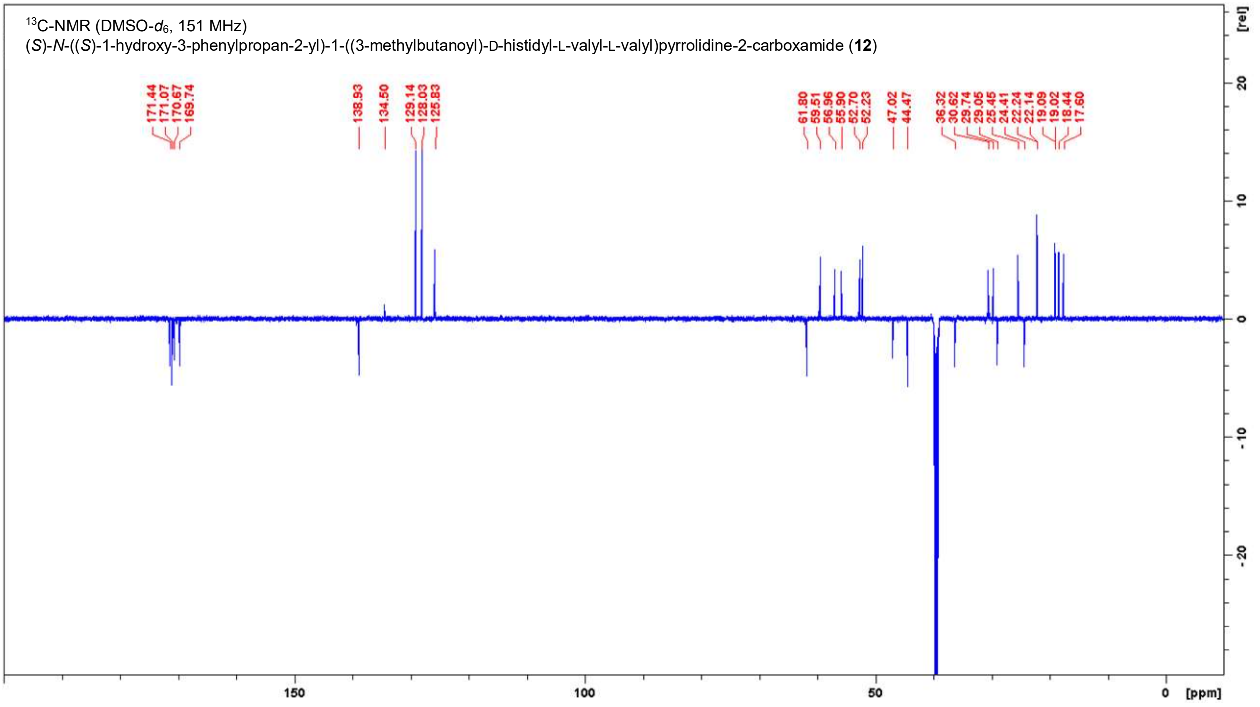

**Figure S13.** NMR spectra of **12** in DMSO-$d_6$.

**Table S7.** NMR data for **12** in DMSO-$d_6$ (600 MHz, 151 MHz).

| **Position** | **δ $^{13}$C [ppm]** | **δ $^{1}$H [ppm]; (m, ∫, *J*)** |
|---|---|---|
| L-Phenylalaninol | | |
| **$CH_2OH$** | 61.8 | 3.27 (t, 2H, *J* = 5.2 Hz) |
| **NH** | - | 7.51 (d, 1H, *J* = 6.1 Hz) |
| **α-CH** | 52.2 | 3.87-3.80 (m, 1H) |
| **β-$CH_2$** | 36.3 | 2.79 (dd, 1H, *J* = 13.7, 6.5 Hz)<br>2.68 (dd, 1H, *J* = 13.7, 7.1 Hz) |
| **γ-$C_{quart}$** | 138.9 | - |
| **$CH_{arom}$** | 129.1, 128.0, 125.8 | 7.27-7.14 (m, 5H) |
| L-Proline | | |
| **CO** | 171.1 | - |
| **α-CH** | 59.5 | 4.30-4.25(m, 2H, overlay) |
| **β-$CH_2$** | 29.1 | 1.97-1.84 (m, 4H, overlay)<br>1.76-1.71 (m, 1H) |
| **γ-$CH_2$** | 24.4 | 1.97-1.84 (m, 4H, overlay)<br>1.81-1.76 (m, 1H) |
| **δ-$CH_2N$** | 47.0 | 3.76-7.70 (m, 1H)<br>3.57-3.51 (m, 1H) |
| L-Valine 1 | | |
| **CO** | 169.7 | - |
| **NH** | - | 7.98 (d, 1H, *J* = 7.9 Hz) |
| **α-CH** | 55.9 | 4.30-4.25(m, 2H, overlay) |
| **β-CH** | 29.7 | 2.03-1.97 (m, 1H) |
| **γ-$CH_3$** | 19.1 | 0.90 (d, 3H, *J* = 6.8 Hz) |
| **γ-$CH_3$** | 18.4 | 0.88 (d, 3H, *J* = 6.6 Hz) |
| L-Valine 2 | | |
| **CO** | 170.7 | - |
| **NH** | - | 7.66 (d, 1H, *J* = 9.2 Hz) |
| **α-CH** | 57.0 | 4.23 (dd, 1H, *J* = 8.2, 6.8 Hz) |
| **β-CH** | 30.6 | 1.97-1.84 (m, 4H, overlay) |
| **γ-$CH_3$** | 19.0 | 0.73 (d, 3H, *J* = 6.8 Hz) |
| **γ-$CH_3$** | 17.6 | 0.70 (d, 3H, *J* = 6.8 Hz) |
| D-Histidine | | |
| **CO** | 171.1 | - |
| **NH** | - | 7.98 (d, 1H, *J* = 7.9 Hz) |
| **α-CH** | 52.7 | 4.59 (ddd, 1H, *J* = 7.9, 7.9, 6.4 Hz) |
| **β-$CH_2$** | 29.8* | 2.90 (dd, 1H, *J* = 14.9, 5.9 Hz)<br>2.73 (dd, 1H, *J* = 14.8, 8.9 Hz) |
| **γ-$C_{quart}$** | n.o. | - |
| **δ-$CH_{arom}$** | n.o. | 6.77 (s, 1H) |
| **ε-$CH_{arom}$** | 134.5, weak | 7.51 (s, 1H) |
| **ε-NH** | - | n.o. |
| Isovaleroyl | | |
| **CO** | 171.4 | - |
| **$CH_2$** | 44.5 | 1.96 (d, 2H, *J* = 6.6 Hz) |
| **CH** | 25.4 | 1.97-1.84 (m, 4H, overlay) |
| **$CH_3$** | 22.2 | 0.81 (d, 3H, *J* = 6.4 Hz) |
| **$CH_3$** | 22.1 | 0.78 (d, 3H, *J* = 6.2 Hz) |

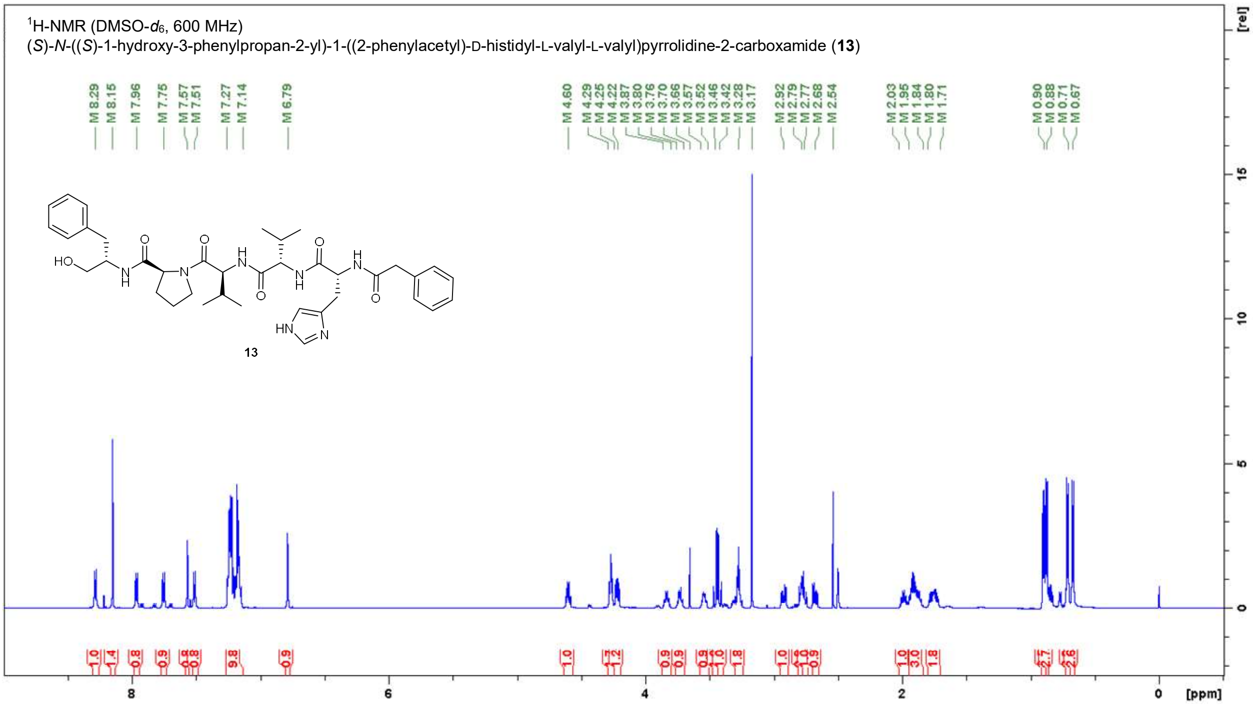

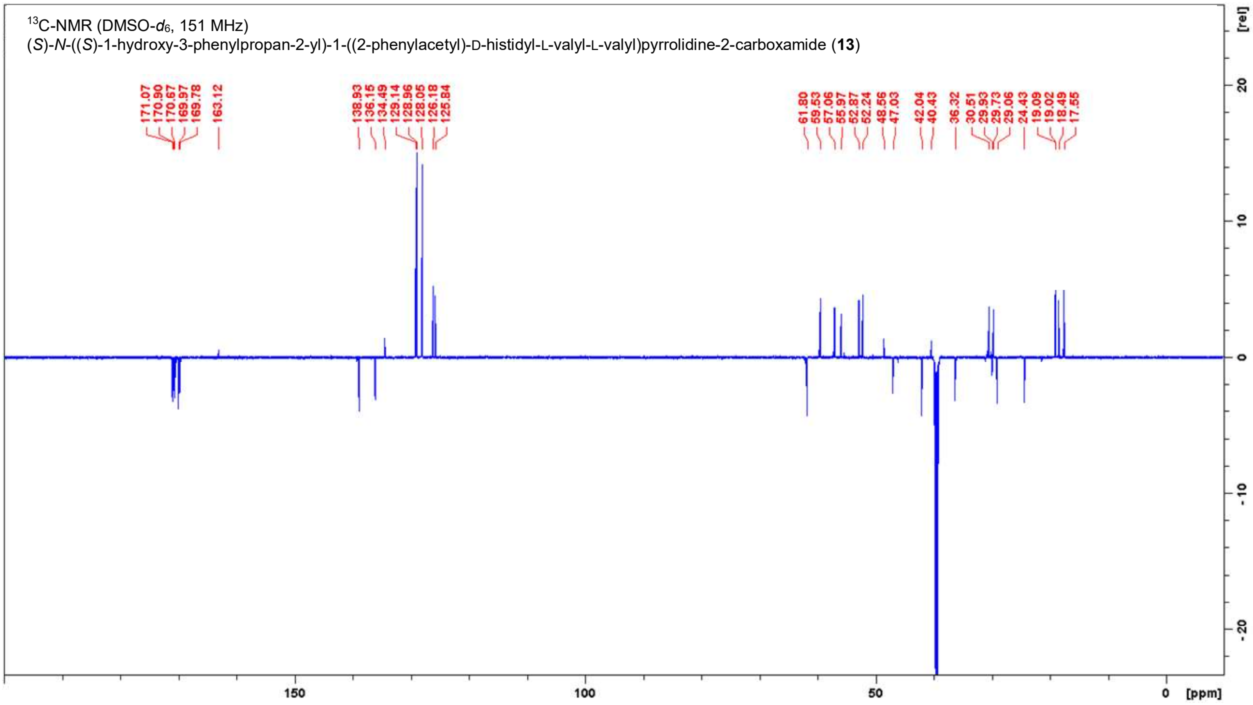

**Figure S14.** NMR spectra of **13** in DMSO-$d_6$.

**Table S8.** NMR data for **13** in DMSO-$d_6$ (600 MHz, 151 MHz).

| Position | δ $^{13}$C [ppm] | δ $^{1}$H [ppm]; (m, ∫, *J*) |
|---|---|---|
| L-Phenylalaninol | | |
| **$CH_2OH$** | 61.8 | 3.28 (t, 2H, *J* = 5.0 Hz) |
| **NH** | - | 7.51 (d, 1H, *J* = 8.4 Hz) |
| **α-CH** | 52.2 | 3.87-3.80 (m, 1H) |
| **β-$CH_2$** | 36.3 | 2.79 (dd, 1H, *J* = 13.4, 6.2 Hz)<br>2.68 (dd, 1H, *J* = 13.8, 7.1 Hz) |
| **γ-$C_{quart}$** | 138.9 | - |
| **$CH_{arom}$** | 129.1, 129.0, 128.0, 126.2, 125.8 | 7.27-7.14 (m, 10H, overlay) |
| L-Proline | | |
| **CO** | 171.1 | - |
| **α-CH** | 59.5 | 4.29-4.25 (m, 2H, overlay) |
| **β-$CH_2$** | 29.1 | 1.95-1.84 (m, 3H, overlay)<br>1.80-1.71 (m, 2H, overlay) |
| **γ-$CH_2$** | 24.4 | 1.95-1.84 (m, 3H, overlay)<br>1.80-1.71 (m, 2H, overlay) |
| **δ-$CH_2N$** | 47.0 | 3.76-3.70 (m, 1H)<br>3.57-3.52 (m, 1H) |
| L-Valine 1 | | |
| **CO** | 169.8 | - |
| **NH** | - | 7.96 (d, 1H, *J* = 8.1 Hz) |
| **α-CH** | 56.0 | 4.29-4.25 (m, 2H, overlay) |
| **β-CH** | 29.7 | 2.03-1.95 (m, 1H) |
| **γ-$CH_3$** | 19.1 | 0.90 (d, 3H, *J* = 6.8 Hz) |
| **γ-$CH_3$** | 18.5 | 0.88 (d, 3H, *J* = 6.8 Hz) |
| L-Valine 2 | | |
| **CO** | 170.7 | - |
| **NH** | - | 7.75 (d, 1H, *J* = 9.7 Hz) |
| **α-CH** | 57.1 | 4.22 (dd, 1H, *J* = 9.0, 6.2 Hz) |
| **β-CH** | 30.5 | 1.95-1.84 (m, 3H, overlay) |
| **γ-$CH_3$** | 19.0 | 0.71 (d, 3H, *J* = 7.0 Hz) |
| **γ-$CH_3$** | 17.6 | 0.67 (d, 3H, *J* = 6.8 Hz) |
| D-Histidine | | |
| **CO** | 170.9 | - |
| **NH** | - | 8.29 (d, 1H, *J* = 7.9 Hz) |
| **α-CH** | 52.9 | 4.60 (ddd, 1H, *J* = 7.9, 7.9, 6.6 Hz) |
| **β-$CH_2$** | 29.9 | 2.92 (dd, 1H, *J* = 14.7, 6.2 Hz)<br>2.77 (dd, 1H, *J* = 14.2, 8.7 Hz) |
| **γ-$C_{quart}$** | 132.9* | - |
| **δ-$CH_{arom}$** | 117.0* | 6.79 (s, 1H) |
| **ε-$CH_{arom}$** | 134.5 | 7.57 (s, 1H) |
| **ε-NH** | - | n.o. |
| Phenylacetyl | | |
| **CO** | 170.0 | - |
| **$CH_2$** | 42.0 | 3.46 (d 1H, *J* = 14.1 Hz)<br>3.42 (d 1H, *J* = 14.3 Hz) |
| **$C_{quart}$** | 136.2 | - |
| **$CH_{arom}$** | 129.1, 129.0, 128.0, 126.2, 125.8 | 7.27-7.14 (m, 10H, overlay) |

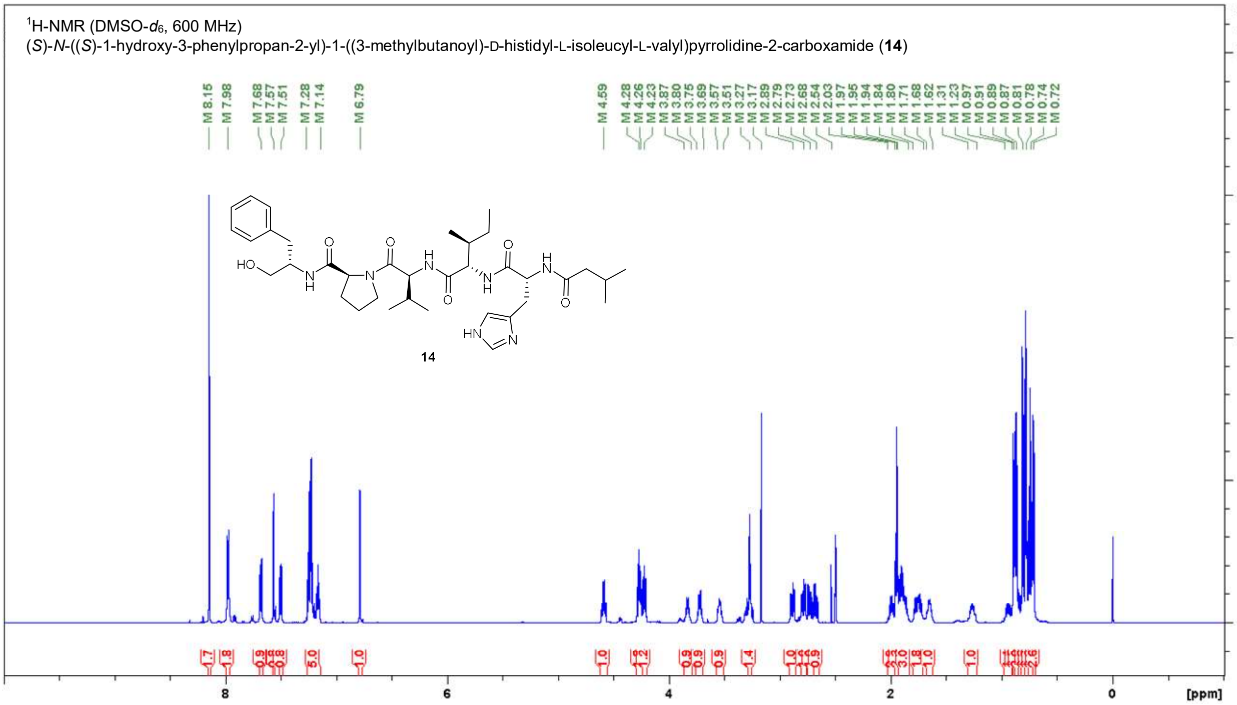

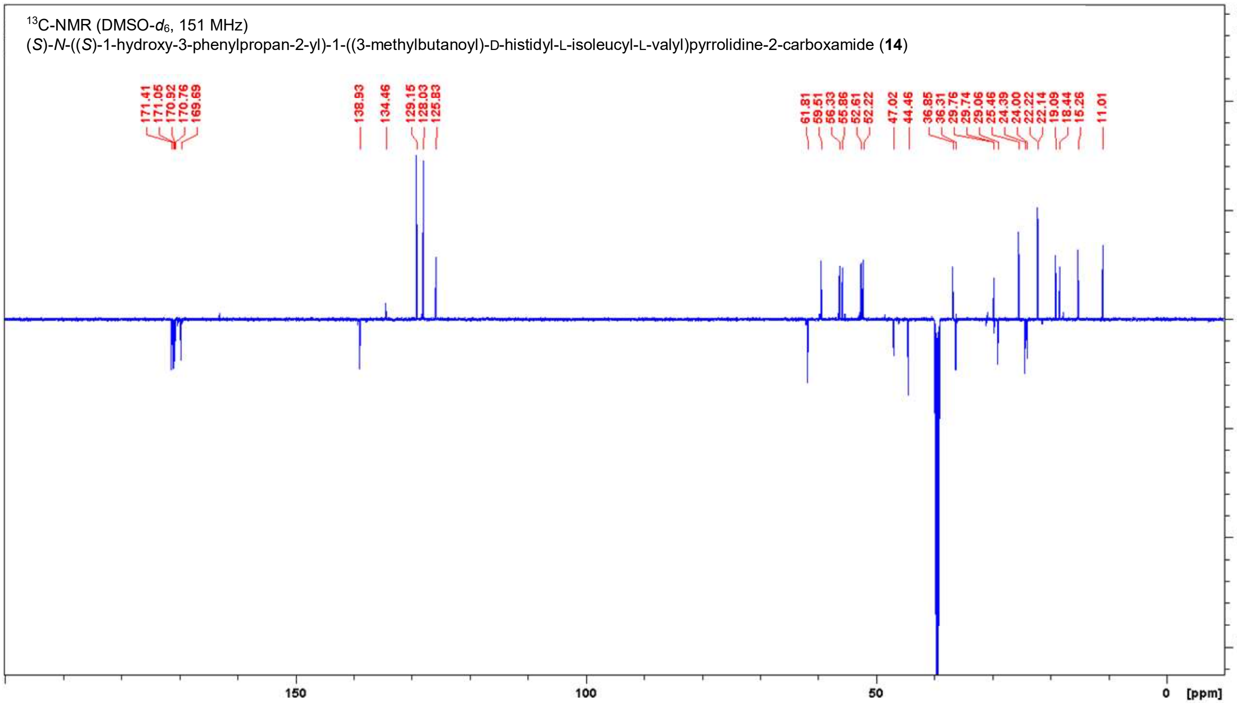

**Figure S15.** NMR spectra of **14** in DMSO-$d_6$.

**Table S9.** NMR data for **14** in DMSO-$d_6$ (600 MHz, 151 MHz).

| Position | δ $^{13}$C [ppm] | δ $^{1}$H [ppm]; (m, ∫, *J*) |
|---|---|---|
| L-Phenylalaninol | | |
| **$CH_2OH$** | 61.8 | 3.27 (t, 2H, *J* = 5.4 Hz) |
| **NH** | - | 7.51 (d, 1H, *J* = 8.3 Hz) |
| **α-CH** | 52.2 | 3.87-3.80 (m, 1H) |
| **β-$CH_2$** | 36.3 | 2.79 (dd, 1H, *J* = 13.7, 6.5 Hz)<br>2.68 (dd, 1H, *J* = 13.9, 7.1 Hz) |
| **γ-$C_{quart}$** | 138.9 | - |
| **$CH_{arom}$** | 129.1, 128.0, 125.8 | 7.28-7.14 (m, 5H) |
| L-Proline | | |
| **CO** | 171.0 | - |
| **α-CH** | 59.5 | 4.26 (dd, 1H, *J* = 8.0, 4.7 Hz) |
| **β-$CH_2$** | 29.1 | 1.94-1.84 (m, 3H, overlay)<br>1.80-1.71 (m, 2H, overlay) |
| **γ-$CH_2$** | 24.4 | 1.94-1.84 (m, 3H, overlay)<br>1.80-1.71 (m, 2H, overlay) |
| **δ-$CH_2N$** | 47.0 | 3.74-3.69 (m, 1H)<br>3.57-3.51 (m, 1H) |
| L-Valine | | |
| **CO** | 169.7 | - |
| **NH** | - | 7.98 (d, 1H, *J* = 8.3 Hz) |
| **α-CH** | 55.9 | 4.28 (dd, 1H, *J* = 7.6, 7.6 Hz) |
| **β-CH** | 29.8 | 2.03-1.97 (m, 1H) |
| **γ-$CH_3$** | 19.1 | 0.89 (d, 3H, *J* = 6.8 Hz) |
| **γ-$CH_3$** | 18.4 | 0.87 (d, 3H, *J* = 6.8 Hz) |
| L-Isoleucine | | |
| **CO** | 170.8 | - |
| **NH** | - | 7.68 (d, 1H, *J* = 9.4 Hz) |
| **α-CH** | 56.3 | 4.23 (dd, 1H, *J* = 8.8, 7.0 Hz) |
| **β-CH** | 36.8 | 1.68-1.62 (m, 1H) |
| **γ-$CH_2$** | 24.0 | 1.31-1.23 (m, 1H)<br>0.97-0.91 (m, 1H) |
| **γ-$CH_3$** | 15.3 | 0.72 (d, 3H, *J* = 6.8 Hz) |
| **δ-$CH_3$** | 11.0 | 0.74 (t, 3H, *J* = 7.5 Hz) |
| D-Histidine | | |
| **CO** | 170.9 | - |
| **NH** | - | 7.98 (d, 1H, *J* = 8.3 Hz) |
| **α-CH** | 52.6 | 4.59 (ddd, 1H, *J* = 8.0, 8.0, 6.5 Hz) |
| **β-$CH_2$** | 29.7 | 2.89 (dd, 1H, *J* = 14.8, 6.0 Hz)<br>2.73 (dd, 1H, *J* = 14.8, 8.5 Hz) |
| **γ-$C_{quart}$** | 133.0* | - |
| **δ-$CH_{arom}$** | 116.9* | 6.79 (s, 1H) |
| **ε-$CH_{arom}$** | 134.5 | 7.57 (s, 1H) |
| **ε-NH** | - | n.o. |
| Isovaleroyl | | |
| **CO** | 171.4 | - |
| **$CH_2$** | 44.5 | 1.95 (d, 2H, *J* = 6.8 Hz) |
| **CH** | 25.5 | 1.94-1.84 (m, 3H, overlay) |
| **$CH_3$** | 22.2 | 0.81 (d, 3H, *J* = 6.4 Hz) |
| **$CH_3$** | 22.1 | 0.78 (d, 3H, *J* = 6.6 Hz) |

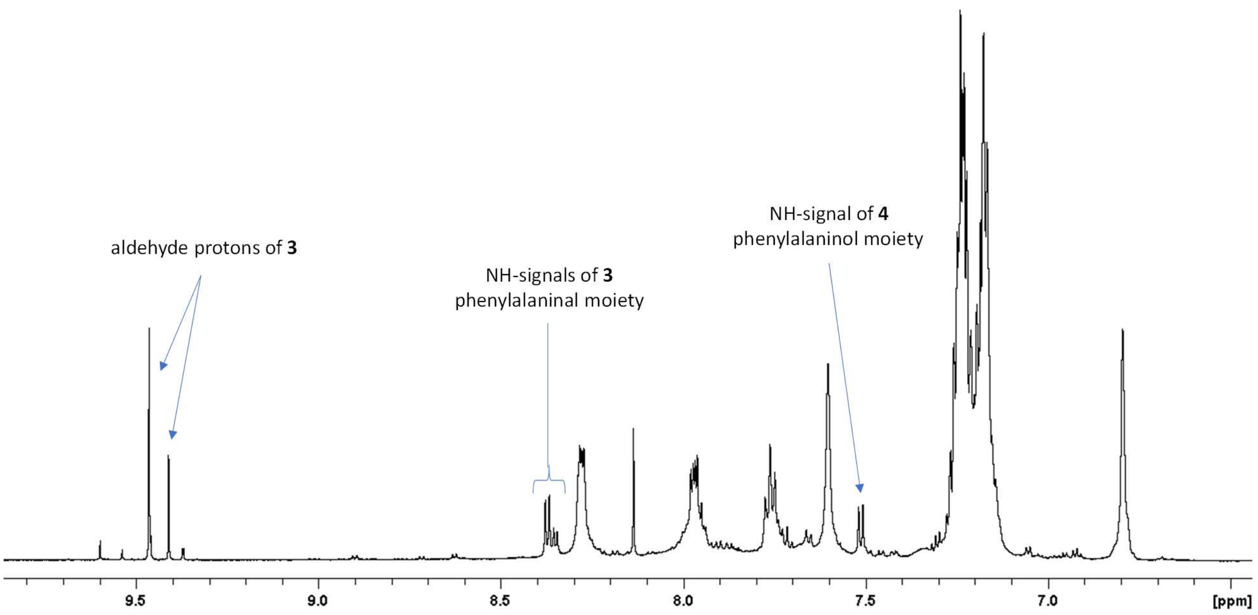

**Figure S16.** 1H NMR section of natural isolated **3** and **4** in DMSO-$d_6$.

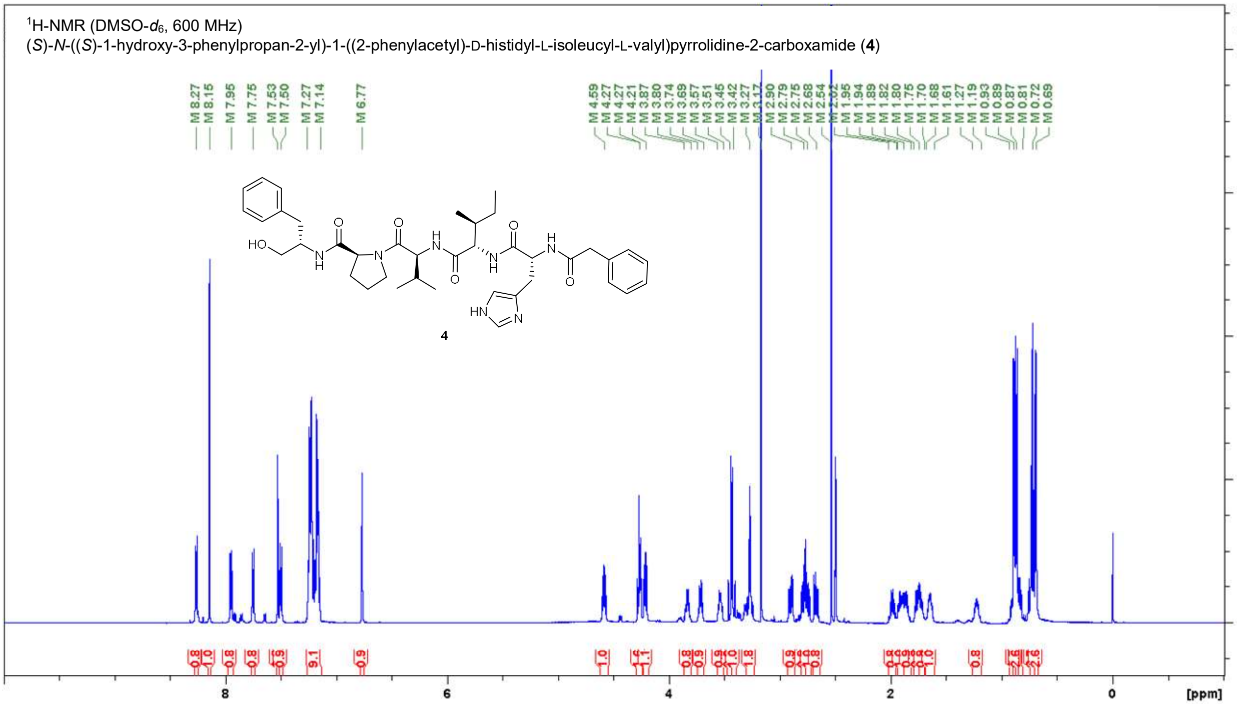

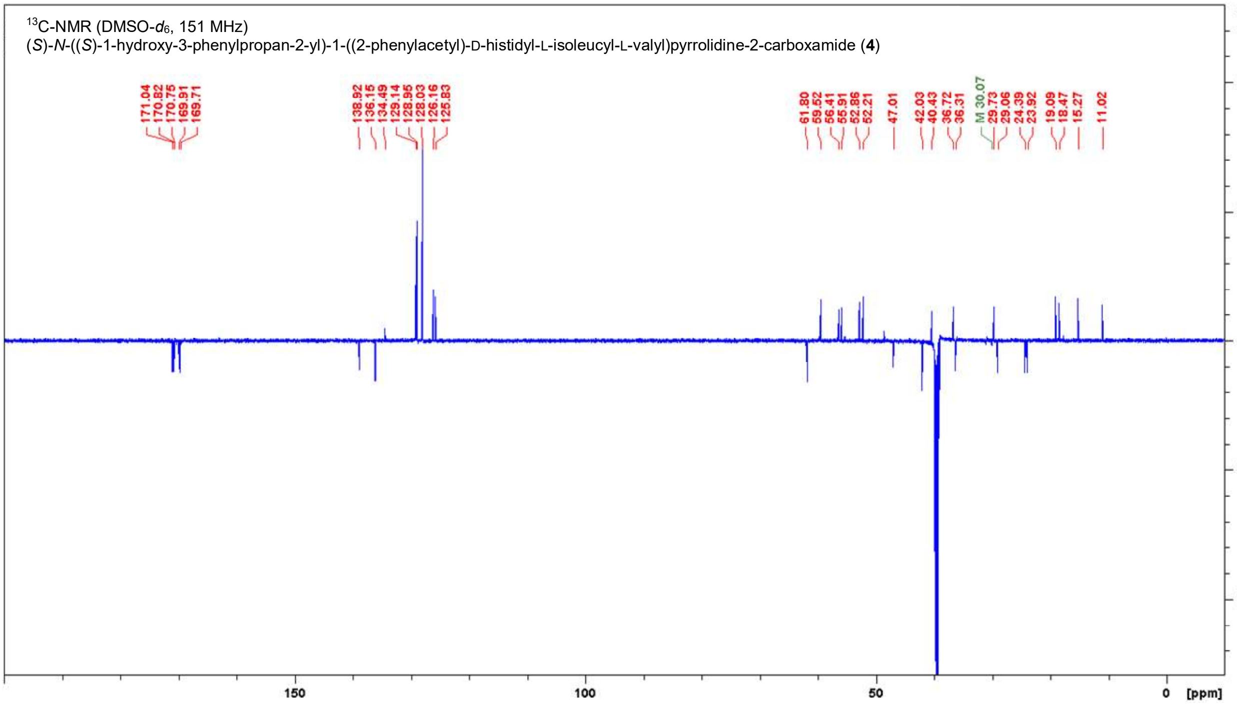

**Figure S17.** NMR spectra of **4** in DMSO-$d_6$.

**Table S10.** NMR data for **4** in DMSO-$d_6$ (600 MHz, 151 MHz).

| Position | δ $^{13}$C [ppm] | δ $^{1}$H [ppm]; (m, ∫, *J*) |
|---|---|---|
| L-Phenylalaninol | | |
| **$CH_2OH$** | 61.8 | 3.27 (t, 2H, *J* = 5.4 Hz) |
| **NH** | - | 7.50 (d, 1H, *J* = 8.4 Hz) |
| **α-CH** | 52.2 | 3.87-3.80 (m, 1H) |
| **β-$CH_2$** | 36.3 | 2.79 (dd, 1H, *J* = 13.5, 6.3 Hz)<br>2.68 (dd, 1H, *J* = 13.8, 7.2 Hz) |
| **γ-$C_{quart}$** | 138.9 | - |
| **$CH_{arom}$** | 129.1, 129.0, 128.0, 126.2, 125.8 | 7.27-7.14 (m, 10H, overlay) |
| L-Proline | | |
| **CO** | 171.0 | - |
| **α-CH** | 59.5 | 4.27 (dd, 1H, *J* = 8.1, 3.9 Hz) |
| **β-$CH_2$** | 29.1 | 1.94-1.89 (m, 1H)<br>1.75-1.70 (m, 1H) |
| **γ-$CH_2$** | 24.4 | 1.89-1.82 (m, 1H)<br>1.80-1.75 (m, 1H) |
| **δ-$CH_2N$** | 47.0 | 3.74-3.69 (m, 1H)<br>3.57-3.51 (m, 1H) |
| L-Valine | | |
| **CO** | 169.7 | - |
| **NH** | - | 7.95 (d, 1H, *J* = 8.3 Hz) |
| **α-CH** | 55.9 | 4.27 (dd, 1H, *J* = 8.1, 8.1 Hz) |
| **β-CH** | 29.7 | 2.02-1.95 (m, 1H) |
| **γ-$CH_3$** | 19.1 | 0.89 (d, 3H, *J* = 6.8 Hz) |
| **γ-$CH_3$** | 18.5 | 0.87 (d, 3H, *J* = 6.8 Hz) |
| L-Isoleucine | | |
| **CO** | 170.82 | - |
| **NH** | - | 7.75 (d, 1H, *J* = 8.8 Hz) |
| **α-CH** | 56.4 | 4.21 (dd, 1H, *J* = 8.6, 7.2 Hz) |
| **β-CH** | 36.7 | 1.68-1.61 (m, 1H) |
| **γ-$CH_2$** | 23.9 | 1.27-1.19 (m, 1H)<br>0.93-0.81 (m, 1H) |
| **γ-$CH_3$** | 15.3 | 0.69 (d, 3H, *J* = 7.2 Hz) |
| **δ-$CH_3$** | 11.0 | 0.72 (t, 3H, *J* = 7.4 Hz) |
| D-Histidine | | |
| **CO** | 170.75 | - |
| **NH** | - | 8.27 (d, 1H, *J* = 7.9 Hz) |
| **α-CH** | 52.9 | 4.59 (ddd, 1H, *J* = 7.9, 7.9, 6.6 Hz) |
| **β-$CH_2$** | 30.1, weak | 2.90 (dd, 1H, *J* = 14.6, 6.1 Hz)<br>2.75 (dd, 1H, *J* = 14.4, 7.8 Hz) |
| **γ-$C_{quart}$** | n.o. | - |
| **δ-$CH_{arom}$** | n.o. | 6.77 (s, 1H) |
| **ε-$CH_{arom}$** | 134.5 | 7.53 (s, 1H) |
| **ε-NH** | - | n.o. |
| Phenylacetyl | | |
| **CO** | 169.9 | - |
| **$CH_2$ a** | 42.0 | 3.45 (d, 1H, *J* = 13.9 Hz)<br>3.42 (d, 1H, *J* = 14.1 Hz) |
| **$C_{quart}$** | 136.2 | - |
| **$CH_{arom}$** | 129.1, 129.0, 128.0, 126.2, 125.8 | 7.27-7.14 (m, 10H, overlay) |

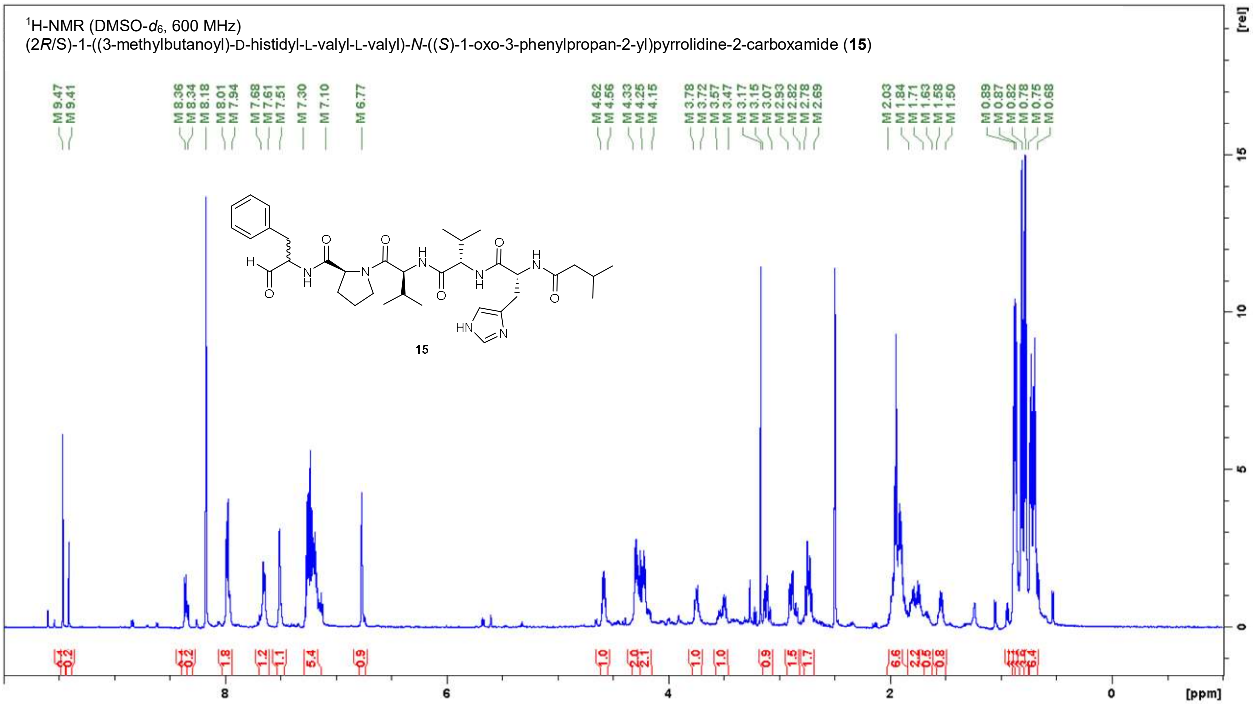

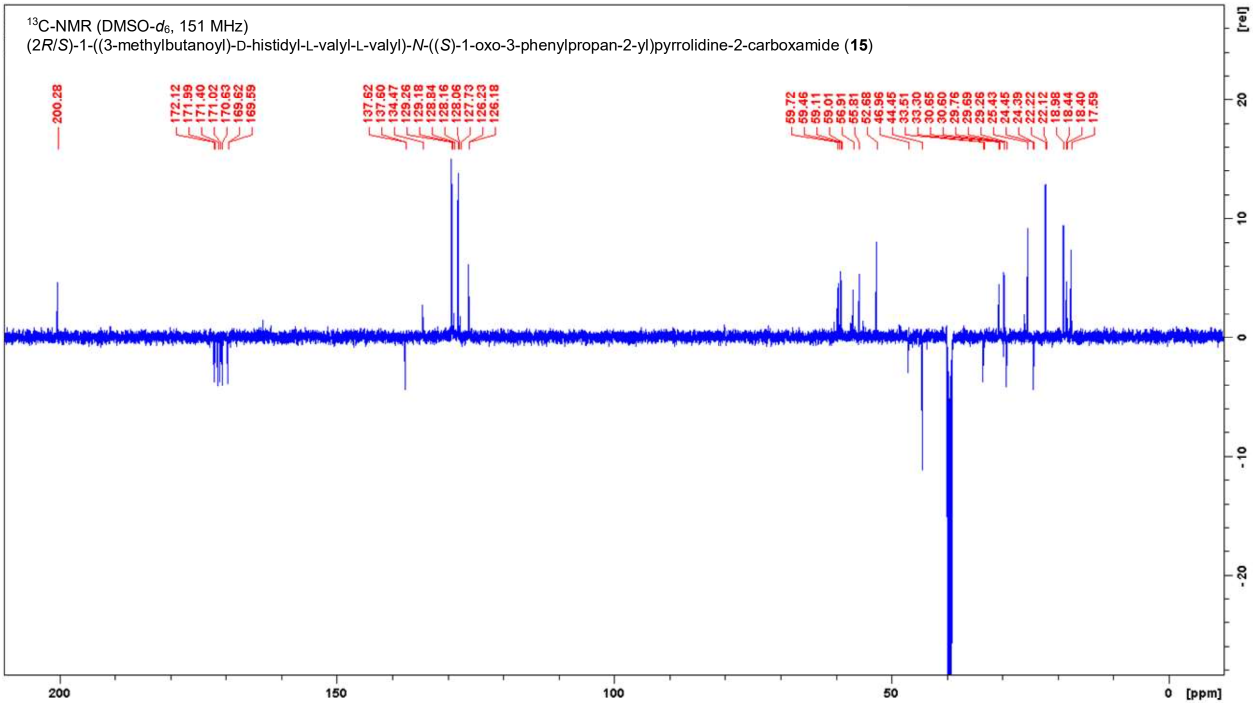

**Figure S18.** NMR spectra of **15** in DMSO-$d_6$.

**Table S11.** NMR data for **15** in DMSO-$d_6$ (600 MHz, 151 MHz).

| Position | δ $^{13}$C [ppm] | | δ $^{1}$H [ppm]; (m, ∫, *J*) |
|---|---|---|---|
| Phenylalaninal | | | |
| **CHO** | 200.3 | | 9.47 (s, 0.4H)<br>9.41 (s, 0.2H) |
| **NH** | - | | 8.36 (d, 0.4H, *J* = 7.5 Hz)<br>8.34 (d, 0.2H, *J* = 7.2 Hz) |
| **α-CH** | 59.7 | 59.5 | 4.33-4.25 (m, 2H, overlay)<br>4.25-4.15 (m, 2H, overlay) |
| **β-CH₂** | 33.5 | 33.3 | 3.15-3.07 (m, 1H)<br>2.93-2.82 (m, 2H, overlay)<br>2.78-2.69 (m, 2H, overlay) |
| **γ-C$_{quart}$** | 137.62 | 137.60 | - |
| **CH$_{arom}$** | 129.3, 129.2, 128.8, 128.2, 128.1, 127.7, 126.23, 126.18 | | 7.30-7.10 (m, 5H) |
| L-Proline | | | |
| **CO** | 172.1 | 172.0 | - |
| **α-CH** | 59.1 | 59.0 | 4.33-4.25(m, 2H, overlay) |
| **β-CH$_2$** | 29.3 | | 2.04-1.84 (m, 7H, overlay)<br>1.84-1.71 (m, 2H)<br>1.58-1.52 (m, 1H) |
| **γ-CH$_2$** | 24.5 | 24.4 | 2.04-1.84 (m, 7H, overlay)<br>1.84-1.71 (m, 2H)<br>1.58-1.52 (m, 1H) |
| **δ-CH$_2$N** | 47.0 | | 3.78-3.72 (m, 1H)<br>3.57-3.47 (m, 1H) |
| L-Valine 1 | | | |
| **CO** | 169.62 | 169.59 | - |
| **NH** | - | | 8.01-7.94 (m, 2H, overlay) |
| **α-CH** | 55.8 | | 4.33-4.25(m, 2H, overlay) |
| **β-CH** | 29.7 | | 2.04-1.84 (m, 7H, overlay) |
| **γ-CH$_3$** | 19.0 | | 0.89 (d, 3H, *J* = 6.8 Hz) |
| **γ-CH$_3$** | 18.44 | 18.40 | 0.87 (d, 3H, *J* = 6.6 Hz) |
| L-Valine 2 | | | |
| **CO** | 170.6 | | - |
| **NH** | - | | 7.68-7.61 (m, 1H) |
| **α-CH** | 56.9 | | 4.25-4.15 (m, 2H, overlay) |
| **β-CH** | 30.65 | 30.60 | 2.04-1.84 (m, 7H, overlay) |
| **γ-CH$_3$** | 19.0<br>17.6 | | 0.75-0.68 (m, 6H) |
| D-Histidine | | | |
| **CO** | 171.0 | | - |
| **NH** | - | | 8.01-7.94 (m, 2H, overlay) |
| **α-CH** | 52.7 | | 4.62-4.56 (m, 1H) |
| **β-CH$_2$** | 29.8 | | 2.93-2.82 (m, 2H, overlay)<br>2.78-2.69 (m, 2H, overlay) |
| **γ-C$_{quart}$** | n.o. | | - |
| **δ-CH$_{arom}$** | n.o. | | 6.77 (s, 1H) |
| **ε-CH$_{arom}$** | 134.5 | | 7.51 (s, 1H) |
| **ε-NH** | - | | n.o. |
| Isovaleroyl | | | |
| **CO** | 171.4 | | - |

| | | |
|---|---|---|
| **$CH_2$** | 44.4 | 2.04-1.84 (m, 7H, overlay) |
| **CH** | 25.4 | 2.04-1.84 (m, 7H, overlay) |
| **$CH_3$** | 22.2 | 0.82 (d, 3H, *J* = 6.4 Hz) |
| **$CH_3$** | 22.1 | 0.78 (d, 3H, *J* = 6.2 Hz) |

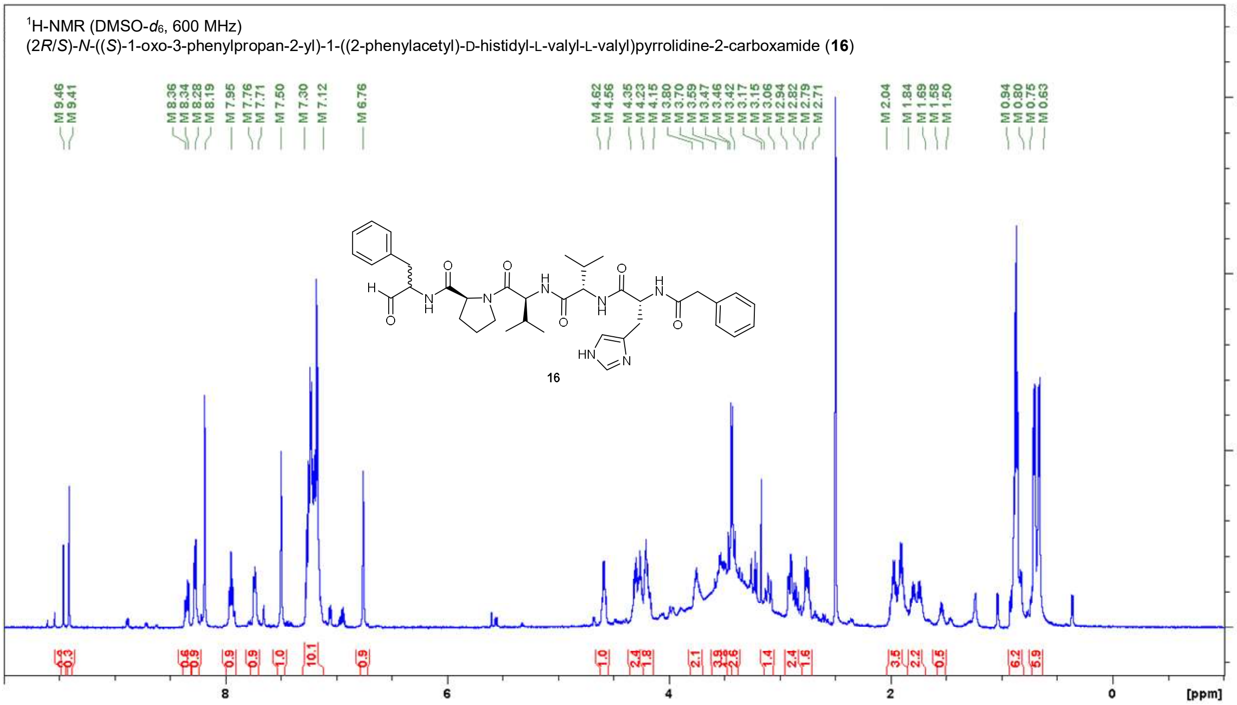

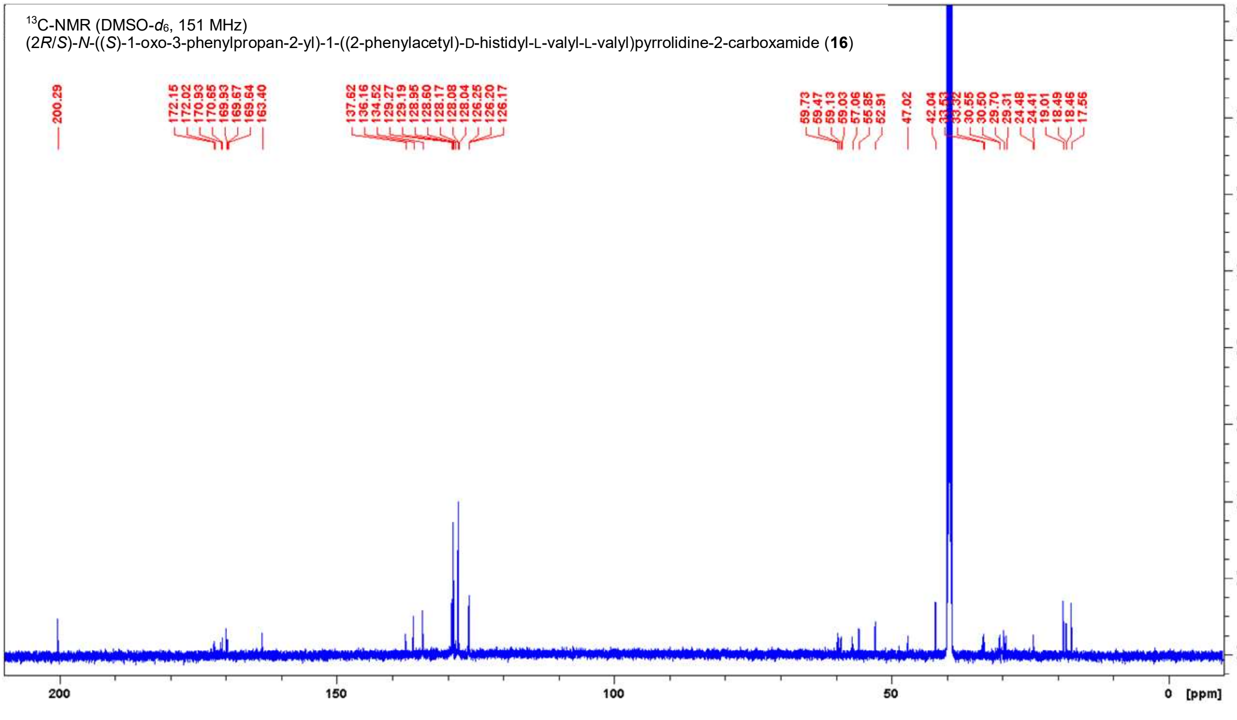

**Figure S19.** NMR spectra of **16** in DMSO-$d_6$.

**Table S12.** NMR data for **16** in DMSO-$d_6$ (600 MHz, 151 MHz).

| Position | δ $^{13}$C [ppm] | | δ $^{1}$H [ppm]; (m, ∫, *J*) |
|---|---|---|---|
| Phenylalaninal | | | |
| **CHO** | 200.3 | | 9.46 (s, 0.2H)<br>9.41 (s, 0.3H) |
| **NH** | - | | 8.36 (d, 0.3H, *J* = 7.5 Hz)<br>8.34 (d, 0.3H, *J* = 6.8 Hz) |
| **α-CH** | 59.7 | 59.5 | 4.35-4.23 (m, 2H, overlay)<br>4.23-4.15 (m, 2 H, overlay) |
| **β-CH₂** | 33.5 | 33.3 | 3.15-3.06 (m, 1H)<br>2.94-2.82 (m, 2H, overlay)<br>2.79-2.71 (m, 2H, overlay) |
| **γ-C$_{quart}$** | 137.6 | | - |
| **CH$_{arom}$** | 129.3, 129.2, 129.0, 128.6, 128.2, 128.1, 128.0, 126.25, 126.20, 126.17 | | 7.30-7.12 (m, 10H, overlay) |
| L-Proline | | | |
| **CO** | not distinguishable | | - |
| **α-CH** | 59.1 | 59.0 | 4.35-4.23 (m, 2H, overlay) |
| **β-CH₂** | 29.3 | | 2.04-1.84 (m, 4H, overlay)<br>1.84-1.69 (m, 2H, overlay) |
| **γ-CH₂** | 24.5 | 24.4 | 2.04-1.84 (m, 4H, overlay)<br>1.58-1.50 (m, 1H, overlay) |
| **δ-CH₂N** | 47.0 | | 3.80-3.70 (m, 2H)<br>3.59-3.47 (m, 4H) |
| L-Valine 1 | | | |
| **CO** | | | - |
| **NH** | - | | 7.95 (dd, 1H, *J* = 8.4, 8.4 Hz) |
| **α-CH** | 55.8 | | 4.35-4.23 (m, 2H, overlay) |
| **β-CH** | 29.7 | | 2.04-1.84 (m, 4H, overlay) |
| **γ-CH₃** | 18.49, 18.46 | | 0.94-0.80 (m, 6H) |
| L-Valine 2 | | | |
| **CO** | not distinguishable | | - |
| **NH** | - | | 7.76-7.71 (m, 1H) |
| **α-CH** | 57.1 | | 4.23-4.15 (m, 2 H, overlay) |
| **β-CH** | 30.6 | 30.5 | 2.04-1.84 (m, 4H, overlay) |
| **γ-CH₃** | 19.0, 17.6 | | 0.75-0.63 (m, 6H) |
| D-Histidine | | | |
| **CO** | not distinguishable | | - |
| **NH** | - | | 8.28 (d, 1H, *J* = 8.1 Hz) |
| **α-CH** | 52.9 | | 4.62-4.56 (m, 1H) |
| **β-CH₂** | 30.0* | | 2.94-2.82 (m, 2H, overlay)<br>2.79-2.71 (m, 2H, overlay) |
| **γ-C$_{quart}$** | n.o. | | - |
| **δ-CH$_{arom}$** | n.o. | | 6.76 (s, 1H) |
| **ε-CH$_{arom}$** | 134.5 | | 7.50 (s, 1H) |
| **ε-NH** | - | | n.o. |
| Phenylacetyl | | | |
| **CO** | 169.9 | | - |
| **CH₂** | 42.0 | | 3.46 (d, 2H, *J* = 13.9 Hz)<br>3.42 (d, 2H, *J* = 14.1 Hz) |

| | | |
|---|---|---|
| **$C_{quart}$** | 136.2 | - |
| **$CH_{arom}$** | 129.3, 129.2, 129.0, 128.6, 128.2, 128.1, 128.0, 126.25, 126.20, 126.17 | 7.30-7.12 (m, 10H, overlay) |

Some integrals aren’t accurate due to the broad peak of $\delta_H$ = 4.00 to 2.59 ppm.

The carbonyl signals 172.1, 172.0, 170.9, 170.7, 169.7, 169.6 were not distinguishable with HMBC data.

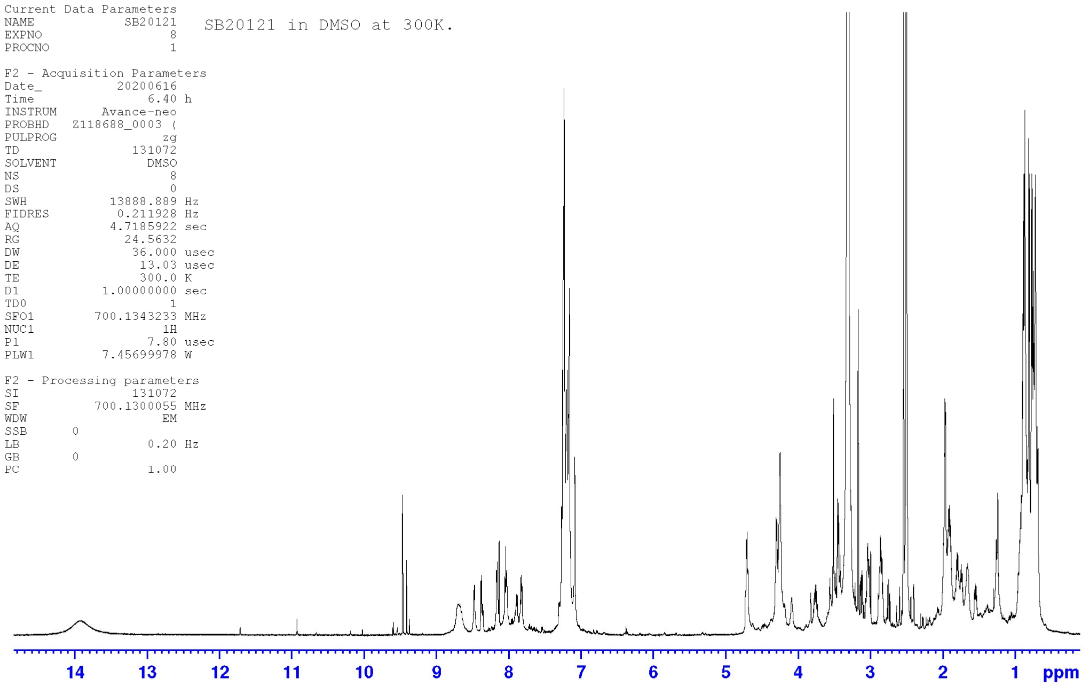

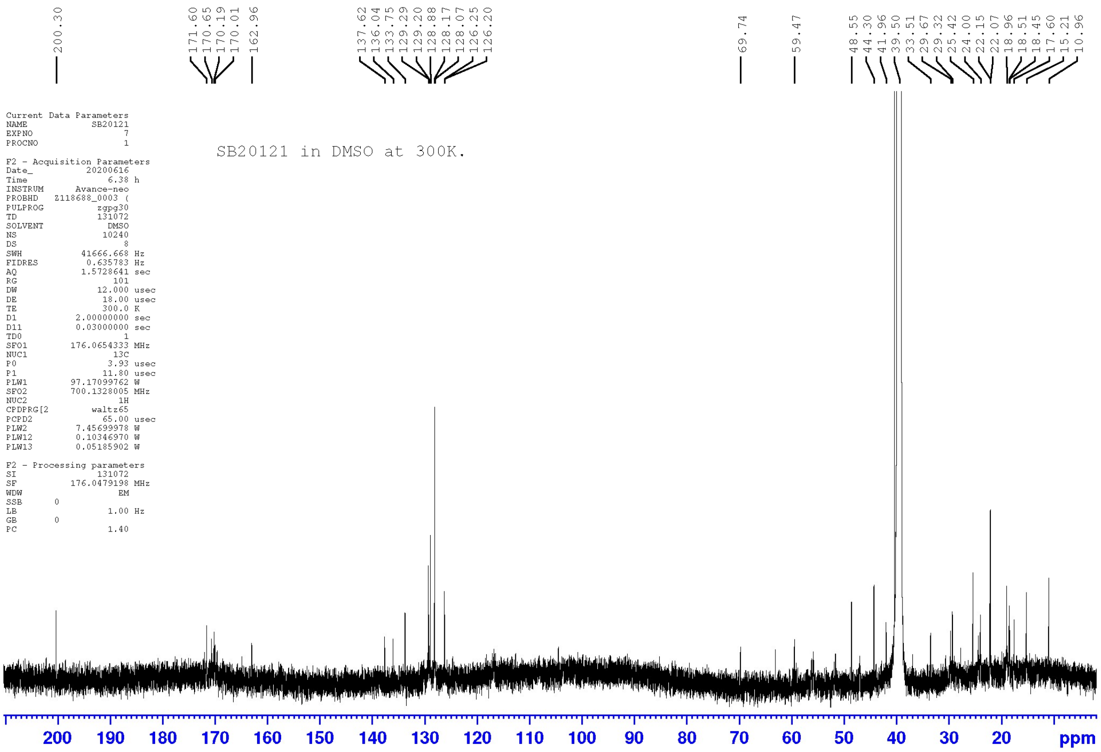

**Figure S20.** NMR spectra of natural isolated compound **2** in DMSO-$d_6$. NMR data can be found in Table 1.

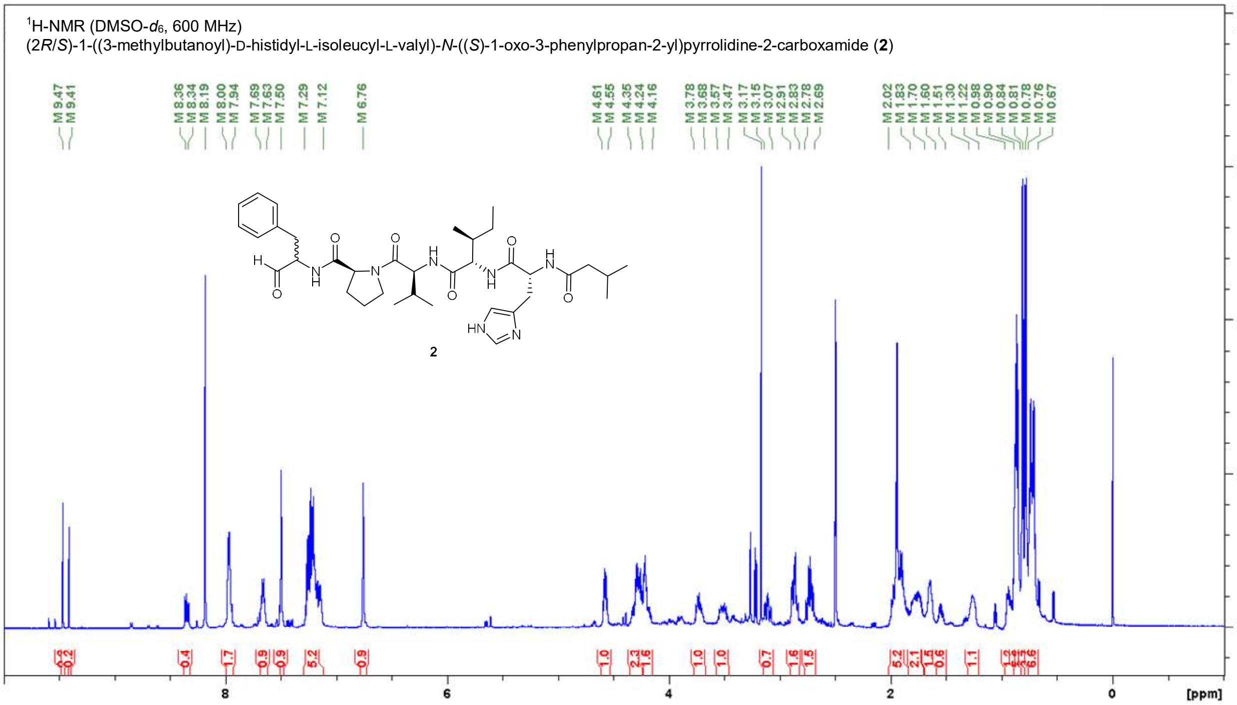

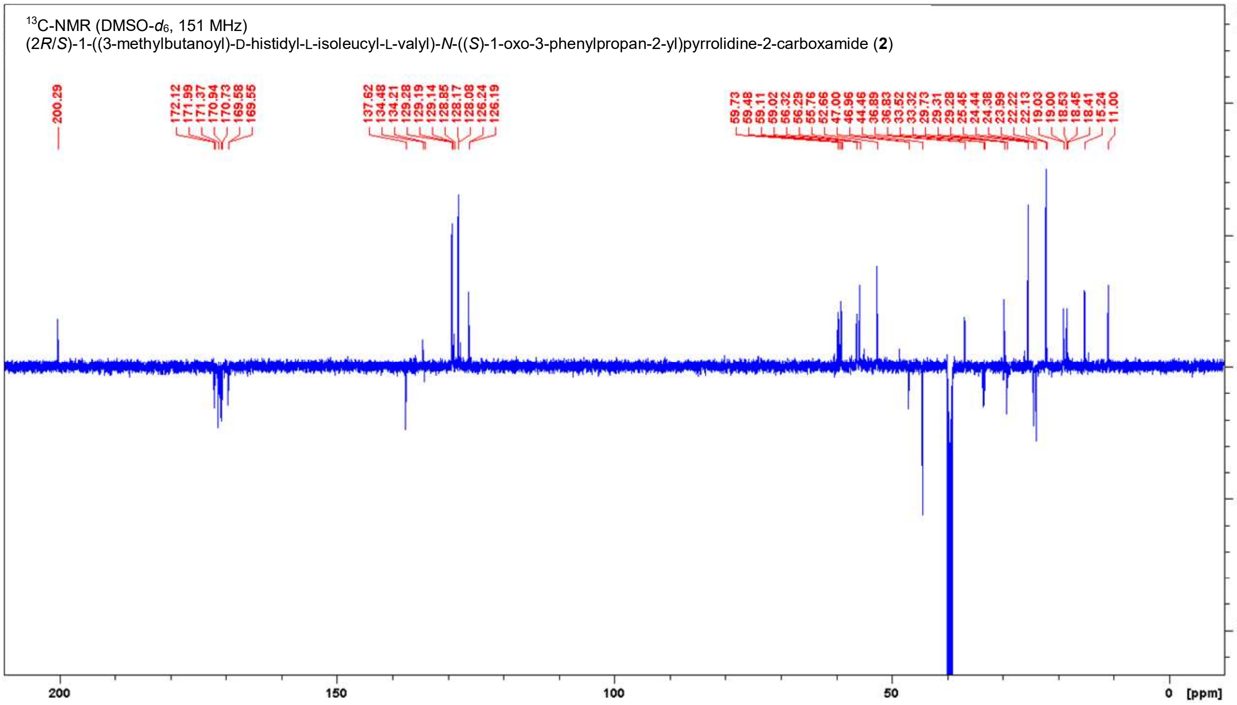

**Figure S21.** NMR spectra of **2** in DMSO-$d_6$.

**Table S13.** NMR data for **2** in DMSO-$d_6$ (600 MHz, 151 MHz).

| **Position** | **δ $^{13}$C [ppm]** | | **δ $^{1}$H [ppm]; (m, ∫, *J*)** |
|---|---|---|---|
| Phenylalaninal | | | |
| **CHO** | 200.3 | | 9.47 (s, 0.2H)<br>9.41 (s, 0.2H) |
| **NH** | - | | 8.36 (d, 0.2H, *J* = 7.7 Hz)<br>8.34 (d, 0.2H, *J* = 7.0 Hz) |
| **α-CH** | 59.7 | 59.5 | 4.35-4.24 (m, 2H, overlay)<br>4.24-4.16 (m, 2H, overlay) |
| **β-CH$_2$** | 33.6 | 33.3 | 3.15-3.07 (m, 1H)<br>2.91-2.83 (m, 2H, overlay)<br>2.78-2.69 (m, 2H, overlay) |
| **γ-C$_{quart}$** | 137.6 | | - |
| **CH$_{arom}$** | 129.3, 129.2, 129.1,<br>128.9, 128.2, 128.1,<br>126.24, 126.19 | | 7.29-7.12 (m, 5H) |
| L-Proline | | | |
| **CO** | 172.1 | 172.0 | - |
| **α-CH** | 59.1 | 59.0 | 4.35-4.24 (m, 2H, overlay) |
| **β-CH$_2$** | 29.31 | 29.28 | 2.02-1.83 (m, 5H, overlay)<br>1.83-1.70 (m, 2H, overlay)<br>1.60-1.51 (m, 1H) |
| **γ-CH$_2$** | 24.44 | 24.38 | 2.02-1.83 (m, 5H, overlay)<br>1.83-1.70 (m, 2H, overlay) |
| **δ-CH$_2$N** | 47.00 | 46.96 | 3.78-3.68 (m, 1H)<br>3.57-3.47 (m, 1H) |
| L-Valine | | | |
| **CO** | 169.58 | 169.55 | - |
| **NH** | - | | 8.00-7.94 (m, 2H, overlay) |
| **α-CH** | 55.7 | | 4.35-4.24 (m, 2H, overlay) |
| **β-CH** | 29.7 | | 2.02-1.83 (m, 5H, overlay) |
| **γ-CH$_3$** | 19.03 | 19.00 | 0.90-0.84 (m, 6H, overlay) |
| **γ-CH$_3$** | 18.45 | 18.41 | 0.90-0.84 (m, 6H, overlay) |
| L-Isoleucine | | | |
| **CO** | 170.7 | | - |
| **NH** | - | | 7.69-7.63 (m, 1H) |
| **α-CH** | 56.32 | 56.29 | 4.24-4.16 (m, 2H) |
| **β-CH** | 36.9 | 36.8 | 1.70-1.60 (m, 2H) |
| **γ-CH$_2$** | 24.0 | | 1.30-1.22 (m, 1H)<br>0.98-0.90 (m, 1H) |
| **γ-CH$_3$** | 15.2 | | 0.76-0.67 (m, 6H, overlay) |
| **δ-CH$_3$** | 11.0 | | 0.76-0.67 (m, 6H, overlay) |
| D-Histidine | | | |
| **CO** | 170.9 | | - |
| **NH** | - | | 8.00-7.94 (m, 2H, overlay) |
| **α-CH** | 52.7 | | 4.61-4.55 (m, 1H) |
| **β-CH$_2$** | 29.9* | | 2.91-2.83 (m, 2H, overlay)<br>2.78-2.69 (m, 2H, overlay) |
| **γ-C$_{quart}$** | 134.2 | | - |
| **δ-CH$_{arom}$** | n.o. | | 6.76 (s, 1H) |
| **ε-CH$_{arom}$** | 134.5 | | 7.50 (s, 1H) |
| **ε-NH** | - | | n.o. |
| Isovaleroyl | | | |

| | | |
|---|---|---|
| **CO** | 171.4 | |
| **$CH_2$** | 44.5 | 2.02-1.83 (m, 5H, overlay) |
| **CH** | 25.5 | 2.02-1.83 (m, 5H, overlay) |
| **$CH_3$** | 22.2 | 0.81 (d, 3H, *J* = 6.4 Hz) |
| **$CH_3$** | 22.1 | 0.78 (d, 3H, *J* = 6.4 Hz) |

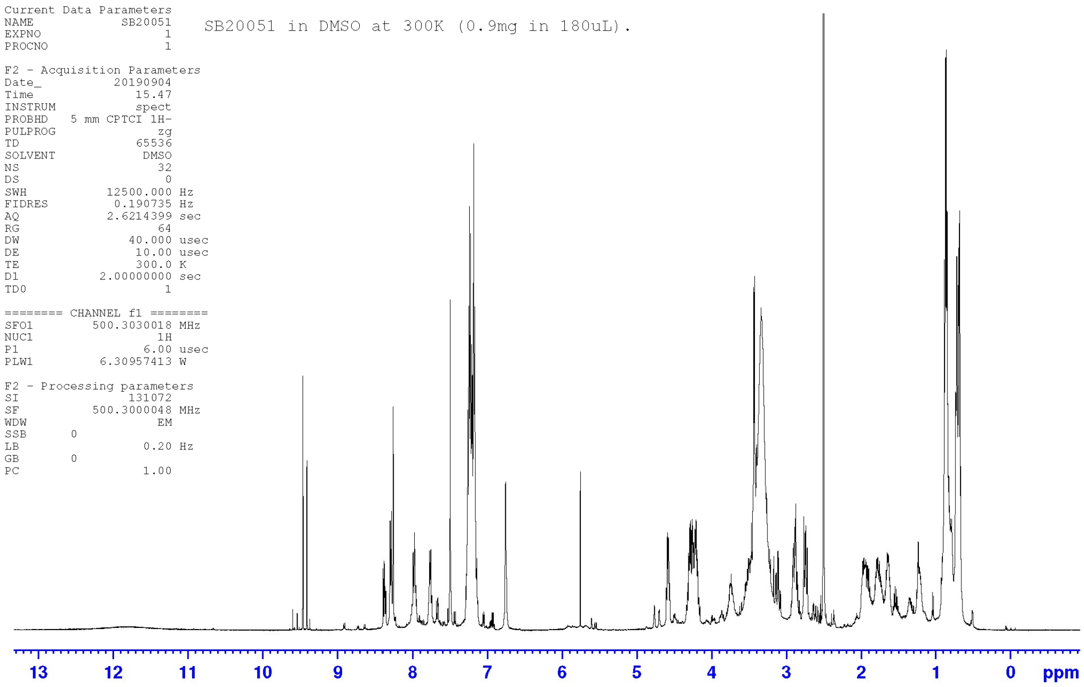

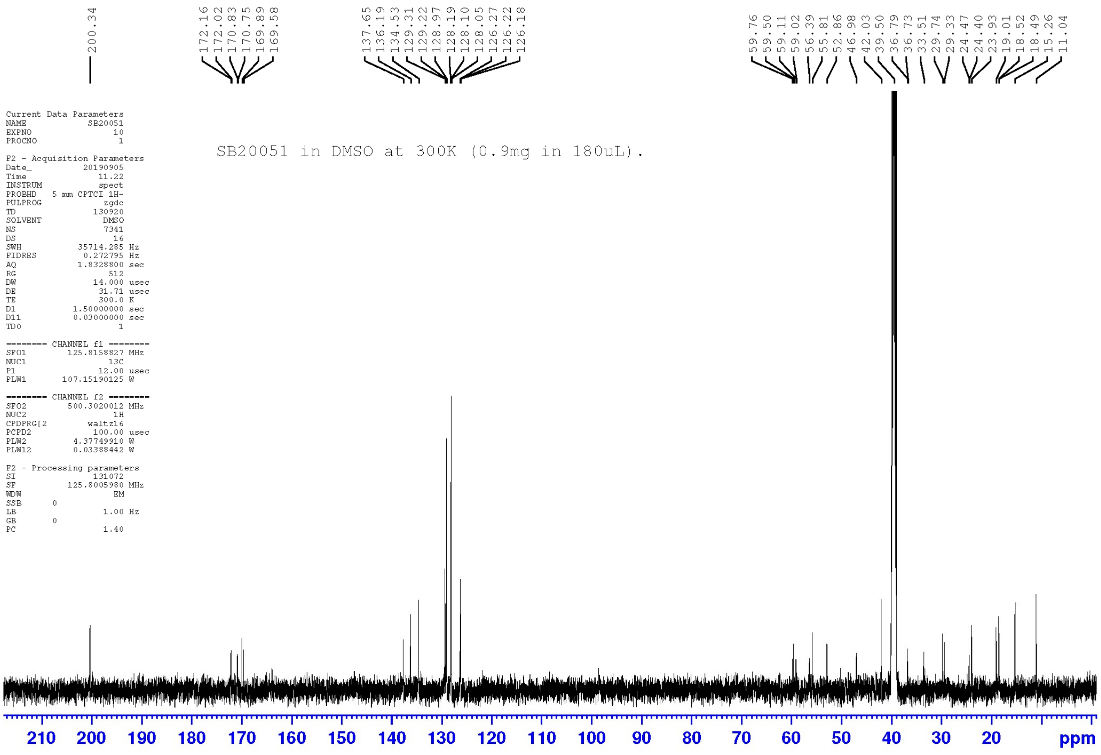

**Figure S22.** NMR spectra of natural isolated compound **3** in DMSO-$d_6$. NMR data can be found in Table 2.

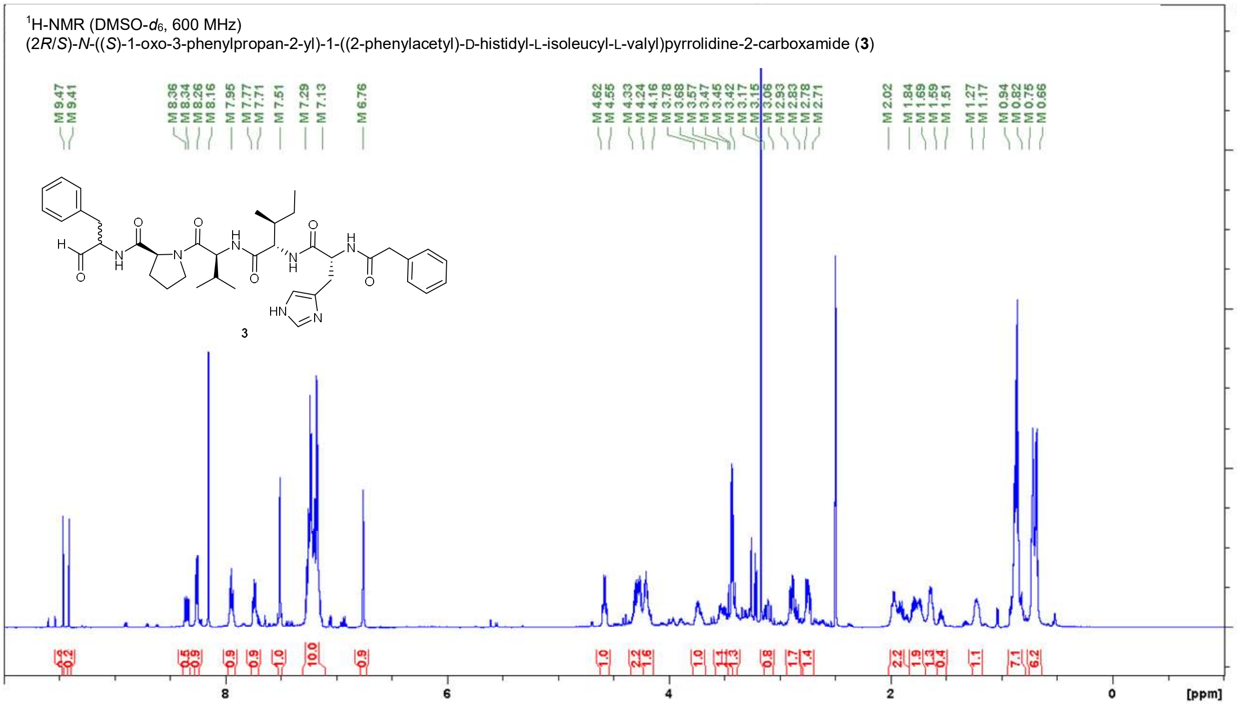

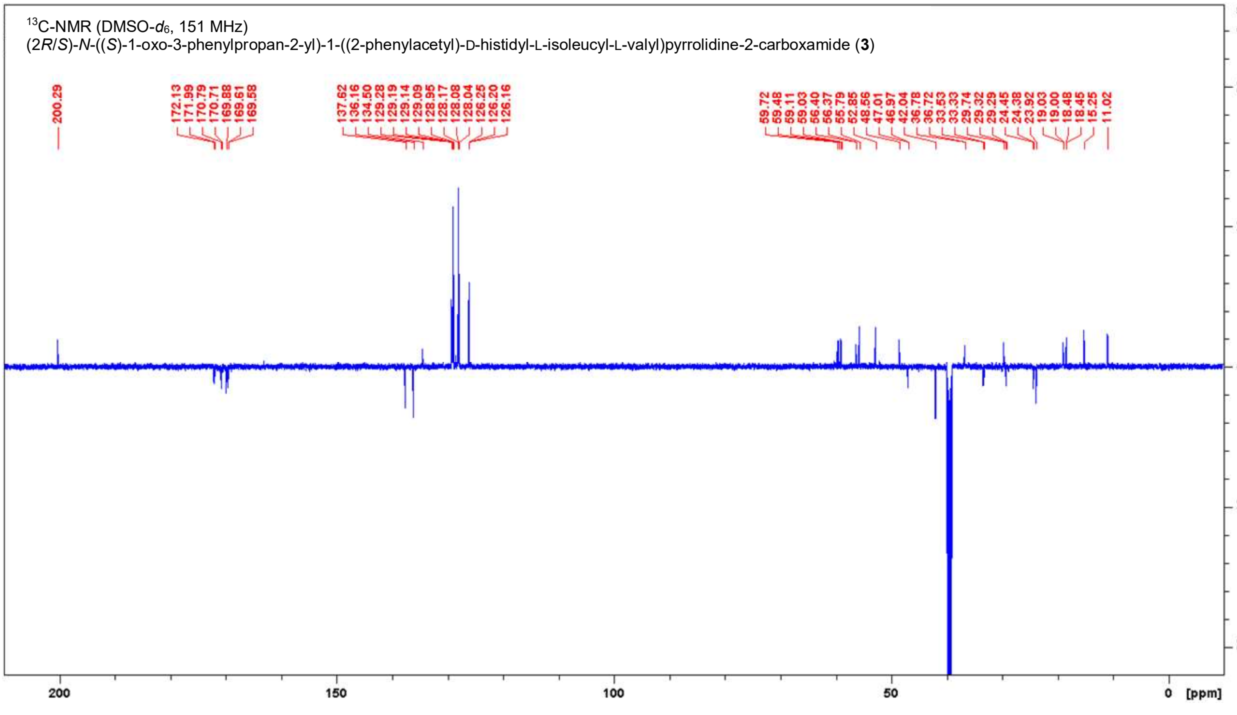

**Figure S23.** NMR spectra of **3** in DMSO-$d_6$.

**Table S14.** NMR data for **3** in DMSO-$d_6$ (600 MHz, 151 MHz).

| **Position** | **δ $^{13}$C [ppm]** | | **δ $^{1}$H [ppm]; (m, ∫, *J*)** |
|---|---|---|---|
| Phenylalaninal | | | |
| **CHO** | 200.3 | | 9.47 (s, .0.2H)<br>9.41 (s, .0.2H) |
| **NH** | - | | 8.36 (d, 0.25H, *J* = 7.7 Hz)<br>8.34 (d, 0.25H, *J* = 7.0 Hz) |
| **α-CH** | 59.1 | 59.0 | 4.24-4.16 (m, 2H, overlay) |
| **β-CH₂** | 33.5 | 33.3 | 3.15-3.06 (m, 1H, overlay)<br>2.93-2.83 (m, 2H, overlay)<br>2.78-2.71 (m, 1H, overlay) |
| **γ-C$_{quart}$** | 137.6 | | - |
| **CH$_{arom}$** | 129.3, 129.2, 129.14, 129.09, 129.0, 128.2, 128.1, 128.0, 126.25, 126.20, 126.16 | | 7.29-7.13 (m, 10H, overlay) |
| L-Proline | | | |
| **CO** | 172.1 | 172.0 | - |
| **α-CH** | 59.5 | | 4.33-4.24 (m, 2H, overlay) |
| **β-CH₂** | 29.32 | 29.29 | 2.02-1.84 (m, 2H, overlay)<br>1.84-1.69 (m, 2H, overlay)<br>1.59-1.51 (m, 1H) |
| **γ-CH₂** | 24.5 | 24.4 | 2.02-1.84 (m, 2H, overlay)<br>1.84-1.69 (m, 2H, overlay) |
| **δ-CH₂N** | 47.01 | 46.97 | 3.78-3.68 (m, 1H)<br>3.57-3.47 (m, 1H) |
| L-Valine | | | |
| **CO** | 169.61 | 169.58 | - |
| **NH** | - | | 7.95 (t, 1H, *J* = 9.0 Hz) |
| **α-CH** | 55.8 | | 4.33-4.24 (m, 2H, overlay) |
| **β-CH** | 29.7 | | 2.02-1.84 (m, 2H, overlay) |
| **γ-CH₃** | 19.03<br>18.5 | 19.00<br>18.4 | 0.94-0.82 (m, 7H, overlay) |
| L-Isoleucine | | | |
| **CO** | 170.7 | | - |
| **NH** | - | | 7.77-7.71 (m, 1H) |
| **α-CH** | 56.40 | 56.37 | 4.24-4.16 (m, 2H, overlay) |
| **β-CH** | 36.8 | 36.7 | 1.69-1.59 (m, 1H) |
| **γ-CH₂** | 23.9 | | 1.27-1.17 (m, 1H)<br>0.94-0.82 (m, 7H, overlay) |
| **γ-CH₃** | 15.3 | | 0.75-0.66 (m, 6H, overlay) |
| **δ-CH₃** | 11.0 | | 0.75-0.66 (m, 6H, overlay) |
| D-Histidine | | | |
| **CO** | 170.8 | | - |
| **NH** | - | | 8.26 (d, 1H, *J* = 8.1 Hz) |
| **α-CH** | 52.9 | | 4.62-4.55 (m, 1H) |
| **β-CH₂** | 30.1* | | 2.93-2.83 (m, 2H, overlay)<br>2.78-2.71 (m, 1H, overlay) |
| **γ-C$_{quart}$** | n.o. | | - |
| **δ-CH$_{arom}$** | n.o. | | 6.76 (s, 1H) |
| **ε-CH$_{arom}$** | 134.5 | | 7.51 (s, 1H) |
| **ε-NH** | - | | n.o. |
| Phenylacetyl | | | |

| | | |
|---|---|---|
| **CO** | 169.9 | - |
| **$CH_2$ a** | 42.0 | 3.45 (d, 1H, *J* = 14.1 Hz)<br>3.42 (d, 1H, *J* = 14.1 Hz) |
| **$C_{quart}$** | 136.2 | - |
| **$CH_{arom}$** | 129.3, 129.2, 129.14, 129.09, 129.0, 128.2, 128.1, 128.0, 126.25, 126.20, 126.16 | 7.29-7.13 (m, 10H, overlay) |

**Table S15**. Summary of molecular docking results.

| | Falcipain-2 | | Falcipain-3 | | Chymotrypsin | |
|---|---|---|---|---|---|---|
| **PDB-ID** | 3BPF | | 3BPM | | 1AFQ | |
| **Reference ligand** | E-64 | | Leupeptin | | D-Leucyl-L-phenylalanyl-p-fluorobenzylamide | |
| **Docking** scores[a] | non-covalent (distance)[b] | covalent | non-covalent (distance)[b] | covalent | non-covalent (distance)[b] | covalent |
| **Re-docking [RMSD, Å]** | -4.8 (3.5) [1.4][d] | -15.7 | -5.8 (3.6) [1.3] | -14.1 | -12.5 (-) [0.8] | - |
| ace-Val-Pro-Phe-alcohol | -3.2 | - | -4.5 | - | -7.1 | - |
| ace-Val-Pro-Phe-aldehyde | -2.0 (3.0) | -16.0 | -2.7 (3.0) | -12.3 | -8.4 (2.0) | -10.1 |
| ace-Val-Pro-**D-Phe-aldehyde** | -2.5 (7.3) | -14.6 | -2.5 (5.0) | -13.1 | -6.8 (3.4) | -12.3 |
| ace-Val-Pro-Phe-acid | -2.9 | - | -3.8 | - | -7.5 | - |
| ace-Val-Pro-amide[c] | -2.0 | - | -3.3 | - | -3.4 | - |
| ace-Val-Val-Pro-**amide**[c] | -3.6 | - | -4.1 | - | -4.2 | - |

[a]Values are docking scores in kcal/mol. [b]Describes distance between electrophilic carbon if present and nucleophilic sulfur of cysteine or oxygen of serine in Å. [c]Truncation of falcitidin. [d]For 3BPF non-covalent docking, potential field generation, and re-docking the binding mode of aligned leupeptin from PDB-ID 3BPM was used as reference.
